# Dissecting intratumor heterogeneity of nodal B cell lymphomas on the transcriptional, genetic, and drug response level

**DOI:** 10.1101/850438

**Authors:** Tobias Roider, Julian Seufert, Alexey Uvarovskii, Felix Frauhammer, Marie Bordas, Nima Abedpour, Marta Stolarczyk, Jan-Philipp Mallm, Sophie Rabe, Peter-Martin Bruch, Hyatt Balke-Want, Michael Hundemer, Karsten Rippe, Benjamin Goeppert, Martina Seiffert, Benedikt Brors, Gunhild Mechtersheimer, Thorsten Zenz, Martin Peifer, Björn Chapuy, Matthias Schlesner, Carsten Müller-Tidow, Stefan Fröhling, Wolfgang Huber, Simon Anders, Sascha Dietrich

**Affiliations:** Department of Medicine V, Hematology, Oncology and Rheumatology, University of Heidelberg, Heidelberg, Germany; Molecular Medicine Partnership Unit (MMPU), Heidelberg, Germany; European Molecular Biology Laboratory (EMBL), Heidelberg, Germany; Bioinformatics and Omics Data Analytics, German Cancer Research Center (DKFZ), Heidelberg, Germany; Faculty of Biosciences, University of Heidelberg, Heidelberg, Germany; Center for Molecular Biology of the University of Heidelberg (ZMBH), Heidelberg, Germany; Division of Molecular Genetics, German Cancer Research Center (DKFZ), Heidelberg, Germany; Department for Translational Genomics, University of Cologne, Cologne, Germany; Division of Chromatin Networks, German Cancer Research Center (DKFZ) & Bioquant, Heidelberg, Germany; Department of Translational Medical Oncology, National Center for Tumor Diseases (NCT) Heidelberg and German Cancer Research Center (DKFZ), Heidelberg, Germany; Department I of Internal Medicine, University Hospital of Cologne, Cologne, Germany; Institute of Pathology, University of Heidelberg, Heidelberg, Germany; Division of Applied Bioinformatics, German Cancer Research Center (DKFZ), Heidelberg, Germany; Department of Medical Oncology and Hematology, University of Zürich, Zürich, Switzerland; Clinic for Hematology and Medical Oncology, University Medicine Göttingen, Göttingen, Germany; German Cancer Consortium (DKTK), Heidelberg, Germany

## Abstract

Tumor heterogeneity encompasses both the malignant cells and their microenvironment. While heterogeneity between individual patients is well-known to affect the efficacy of anti-cancer drugs, most personalized treatment approaches do not account for intratumor heterogeneity. We addressed this issue by studying the heterogeneity of lymph node-derived B cell non-Hodgkin lymphoma (B-NHL) by single cell RNA-sequencing (scRNA-seq) and transcriptome-informed flow cytometry. We identified transcriptionally distinct malignant subclones and compared their drug response and genomic profiles. Malignant subclones of the same patient responded strikingly different to anti-cancer drugs ex vivo, which recapitulated subclone-specific drug sensitivity during in vivo treatment. Tumor infiltrating T cells represented the majority of non-malignant cells, whose gene expression signatures were similar across all donors, whereas the frequencies of T cell subsets varied significantly between the donors. Our data provide new insights into the heterogeneity of B-NHL and highlight the relevance of intratumor heterogeneity for personalized cancer therapies.

## Introduction

The genomic and transcriptional landscape of many cancer entities has been catalogued over recent years, documenting the range of tumor heterogeneity between individual patients [1]. In addition, it has long been appreciated that tumors within each patient consist of diverse, but phylogenetically-related subclones [2]. Bulk sequencing studies of tumor cells have been conducted to infer the genetic spectrum of intratumor heterogeneity from variant allele frequencies of somatic mutations [3]. While important insights were gained from these studies, further characterization on the single cell level is needed to more accurately dissect the pathway and molecular properties associated with distinct subclones.

Neoplastic cells alone do not manifest a malignant disease, but attract a battery of non-malignant bystander cells, which support tumor cell growth and survival. The diversity and plasticity of the microenvironment constitutes another layer of heterogeneity, beyond the heterogeneity of the cancer cells themselves [4]. There is solid evidence that intratumor heterogeneity among malignant and non-malignant cells, and their interactions within the tumor microenvironment are critical to diverse aspects of tumor biology, response to treatment, and prognosis [5].

While bulk genomic tissue profiling has only a limited ability to reconstruct the complex cellular composition of tumors, single cell DNA-sequencing [6, 7] and RNA-sequencing (scRNA-seq) methods [8–11] have emerged as powerful tools to study intratumor heterogeneity and reconstruct the full picture of malignant and non-malignant cells. These technologies further enable researchers to identify rare cell types such as cancer stem cells [12] and circulating tumor cells [13, 14], or to follow clonal dynamics during cancer treatment [15]. Most of these single cell studies have been used to describe distinct cell subpopulations on the transcriptional level, but their functional properties, such as drug response profiles, remain largely unexplored.

To address this, we used B cell non-Hodgkin lymphoma (B-NHL) as a model disease entity to dissect intratumor heterogeneity on the transcriptional, genetic, and functional (drug response) level. In parallel, we investigated the cellular heterogeneity of the B-NHL lymph node microenvironment. B-NHL are a heterogenous group of hematologic malignancies that most frequently grow in the lymph node compartment. Almost half of all B-NHL are classified as diffuse large B cell lymphoma (DLBCL) or follicular lymphoma (FL) [16]. Transformation of indolent FL into aggressive DLBCL is observed in approximately 10% of all FL cases [17]. Despite effective treatment options, 20-40% of B-NHL patients relapse multiple times and present with chemotherapy refractory disease [18, 19]. The response to single agent targeted therapy in these patient cohorts is surprisingly low [20, 21]. Intratumor heterogeneity might be a key factor contributing to therapeutic failure and low success rate of these single agent targeted therapies [3]. Understanding subclonal drug response patterns would therefore be an important asset for designing more effective personalized lymphoma therapies.

To dissect the complex cellular composition of the malignant lymph node niche, we profiled transcriptomes of malignant and non-malignant cells derived from 12 different reactive or B-NHL lymph node biopsies. We further studied the variation of the cellular composition of the malignant lymph node niche by flow cytometry in a larger cohort of 41 patients. Among malignant cells, we identified transcriptionally distinct malignant subclones and characterized these subclones further by ex-vivo drug perturbation and genome sequencing. This revealed new insights into intratumor heterogeneity of B-NHL and demonstrated substantially different drug responses between malignant subclones in the same patient.

## Results

### Study outline

We designed an experimental pipeline to dissect the heterogeneity of non-malignant and malignant lymph node-derived lymphocytes (Figure 1A). This involved first preparing single cell suspensions of B-NHL lymph node biopsies and performing scRNA-seq. These single cell transcriptomic data were then used to identify transcriptionally-distinct subclones by flow cytometry using distinguishing subclone-specific surface markers, and finally the subclones were functionally interrogated in drug perturbation assays with a comprehensive panel of 58 drugs in five concentrations, and further characterized by whole genome (WGS) and/or exome sequencing (WES).

**Figure 1.**
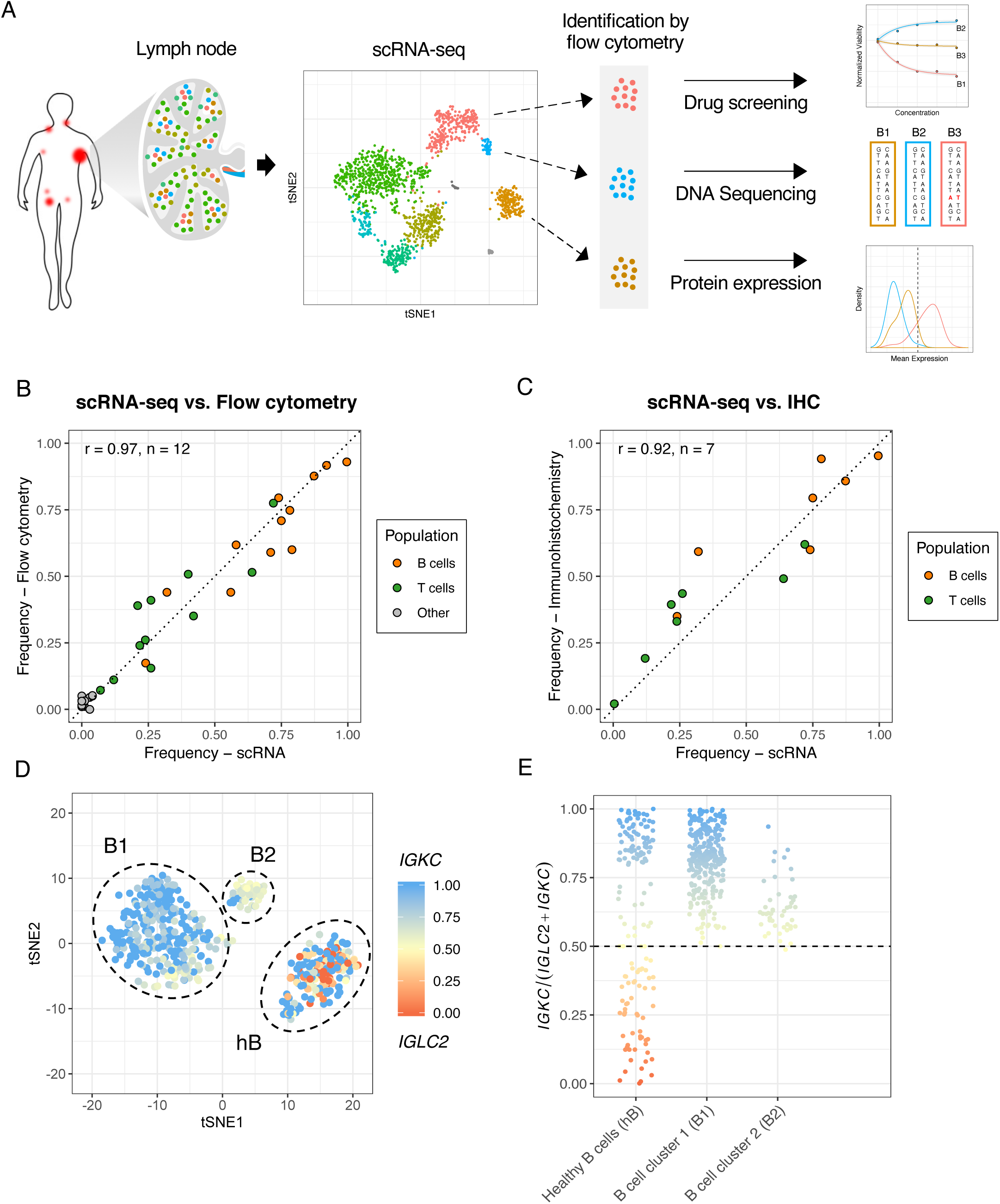
Identification of cell types using scRNA-seq. A) Schematic overview of the study design. B and C) Lymph node-derived B and T cells were quantified by scRNA-seq, flow cytometry and immunohistochemistry (IHC) of paraffin tissue sections (see Supplementary Figure 1 for details). The frequencies of B and T cells were correlated for B) scRNA-seq and flow cytometry or C) scRNA-seq and IHC. Pearson’s correlation coefficients (r) and the number of samples included (n) are given in the left top corner. D and E) Illustration of the strategy to identify malignant B cells. Single cell RNA expression profiles of B cells derived from the tFL1 sample were visualized by t-SNE. The different B cell clusters are circled and labeled with hB (healthy B cells), B1 (B cell cluster 1) and B2 (B cell cluster 2). For each single B cell we calculated the the kappa light chain (IGKC) fraction *IGKC*/(*IGKC*+*IGLC2)* (see color code D and E). If this IGKC-fraction was > 0.5, we classified a B cell as a kappa positive and if this ratio was below 0.5 we classified the B cell as a lambda positive. The percentage of B cells either expressing kappa or lambda per transcriptionally distinct B cell cluster was calculated. The non-malignant healthy B cell (hB) cluster contained approximately 50% kappa and 50% lambda expressing B cells while the tow malignant cluster (B1, B2) contained B cells homogeneously expressing the kappa light chain.

### Dissecting the cellular composition of nodal B cell lymphomas

We assayed single cell suspensions of a total of 12 samples: four germinal center-derived diffuse large B cell lymphoma (DLBCL) samples, of which two were transformed from FL (DLBCL1, DLBCL2, tFL1, tFL2), one non-germinal center-derived DLBCL (DLBCL3), four follicular lymphoma samples (FL1, FL2, FL3, FL4), and three reactive non-malignant lymph node sample (rLN1, rLN2, rLN3) by flow cytometry and droplet-based scRNA-seq (Supplementary Table 1). After removal of low-quality cells, we analyzed scRNA-seq profiles of 13,259 malignant and 9,296 non-malignant cells with an average sequencing depth of 1,409 genes per cell.

First, we verified that the lymph node-derived single cell suspensions were representative for the cellular composition (B and T cells) of the lymphoma and its microenvironment in vivo. We used sections of paraffin-embedded samples of the same lymph nodes, which were formalin fixed directly after surgical excision and therefore represent the in vivo cellular composition, and quantified B and T cell frequencies by immunohistochemistry (IHC, Supplementary Figure 1A). In parallel, we calculated B and T cell frequencies by flow cytometry and scRNA-seq in single cell suspensions (Supplementary Figure 1A, B). The frequencies of B and T cells derived from scRNA-seq correlated very well with the frequencies determined by flow cytometry (r = 0.97, n = 12, Figure 1B) and IHC (r = 0.92, n = 7, Figure 1C).

Next, we aimed to distinguish malignant from non-malignant B cells and delineate these populations in our single cell experiments. We took advantage of the fact that malignant B cell populations express only one type of immunoglobulin light chain (LC), either κ or λ [22]. We calculated the LC-ratio (κ/λ) based on RNA expression of the genes *IGKC* (coding for the constant part of the κ LC) and *IGLC2* (λ LC) for each single B cell and color-coded this ratio in a t-distributed stochastic neighbor embedding (t-SNE) plot (Figure 1D, E). In the malignant lymph nodes, we could either identify both a non-malignant and malignant or only malignant B cell clusters (Supplementary Figure 2). In contrast, reactive lymph node samples contained only non-malignant B cells (see method section for details).

We further evaluated the frequencies of these subsets in a larger cohort of 41 lymph node samples by flow cytometry, including those samples used for scRNA-seq. Both approaches showed very similar frequencies of these cell subsets (r = 0.97, n = 12, Supplementary Figure 3A, B). We found that the proportion of malignant cells was highly variable across samples. It ranged from 14.6 to 97.2 % (median 79.3 %, n = 9) in DLBCL, 23.7 to 85.4 % (median 79.9 %, n = 12) in FL, 48.4 to 95.5 % (median 88.0 %, n = 4) in mantle cell lymphoma, and 65.4 to 91.4 % (median 83.1 %, n = 7) in chronic lymphocytic leukemia (Supplementary Figure 3C). This substantial cellular heterogeneity complicates bulk sequencing approaches of unsorted lymph node samples, and highlights the value of single cell sequencing to simultaneously study the full spectrum of malignant and non-malignant lymph node cells.

### Characterization of lymph node-derived T cell populations

T cells are key players of the host-specific tumor immunosurveillance [23]. B-NHL exhibit genetic immune escape strategies that can be targeted using current therapeutic strategies [24], including checkpoint inhibitors [25] and bispecific antibodies [26]. Notably, lymphoma cells can also orchestrate their tumor microenvironment so that certain T cell subsets support the growth and proliferation of the tumor cells [27]. Even though these subsets have been extensively studied by immunophenotyping, their transcriptional heterogeneity in B-NHL lymph nodes, in particular at the single cell level, still remains to be elucidated.

We combined single cell RNA expression profiles of T cells from all 12 donors and jointly visualized them by Uniform Manifold Approximation and Projection (UMAP), a dimension reduction algorithm alternative to t-SNE [28]. Many well-established surface markers, which are used to distinguish T cell subsets in flow cytometry studies, are insufficiently expressed on the scRNA-seq level. We therefore chose unsupervised clustering to partition T cells into transcriptionally distinct subsets, which were then annotated by differentially expressed marker genes. All T cells from either reactive or malignant lymph nodes distributed to only four major T cell subpopulations (Figure 2 A, B). Note, that clusters were not driven by patients or disease entity, suggesting only limited transcriptional heterogeneity across all donors. Apart from conventional T helper cells (T_H;_ *CD4, IL7R, PLAC8, KLF2*) and regulatory T cells (T_REG;_ *CD4, IL2RA, FOXP3, ICOS*), we identified a third T helper cell population, which was characterized by overexpression of *PDCD1* (PD1)*, ICOS, CXCR5, TOX, TOX2* and *CD200* (Figure 2C, Supplementary Table 2), suggesting a T follicular helper cell (T_FH_) phenotype [29–33]. In contrast to the diversity of T helper cells, we observed only one major cluster of cytotoxic T cells (T_TOX_, *GZMK*, *CCL4/5*, *GZMA*, *NKG7*, *CD8A*). However, the frequencies of the four identified T cell subsets were highly variable between different B-NHL donors (Figure 2D).

**Figure 2.**
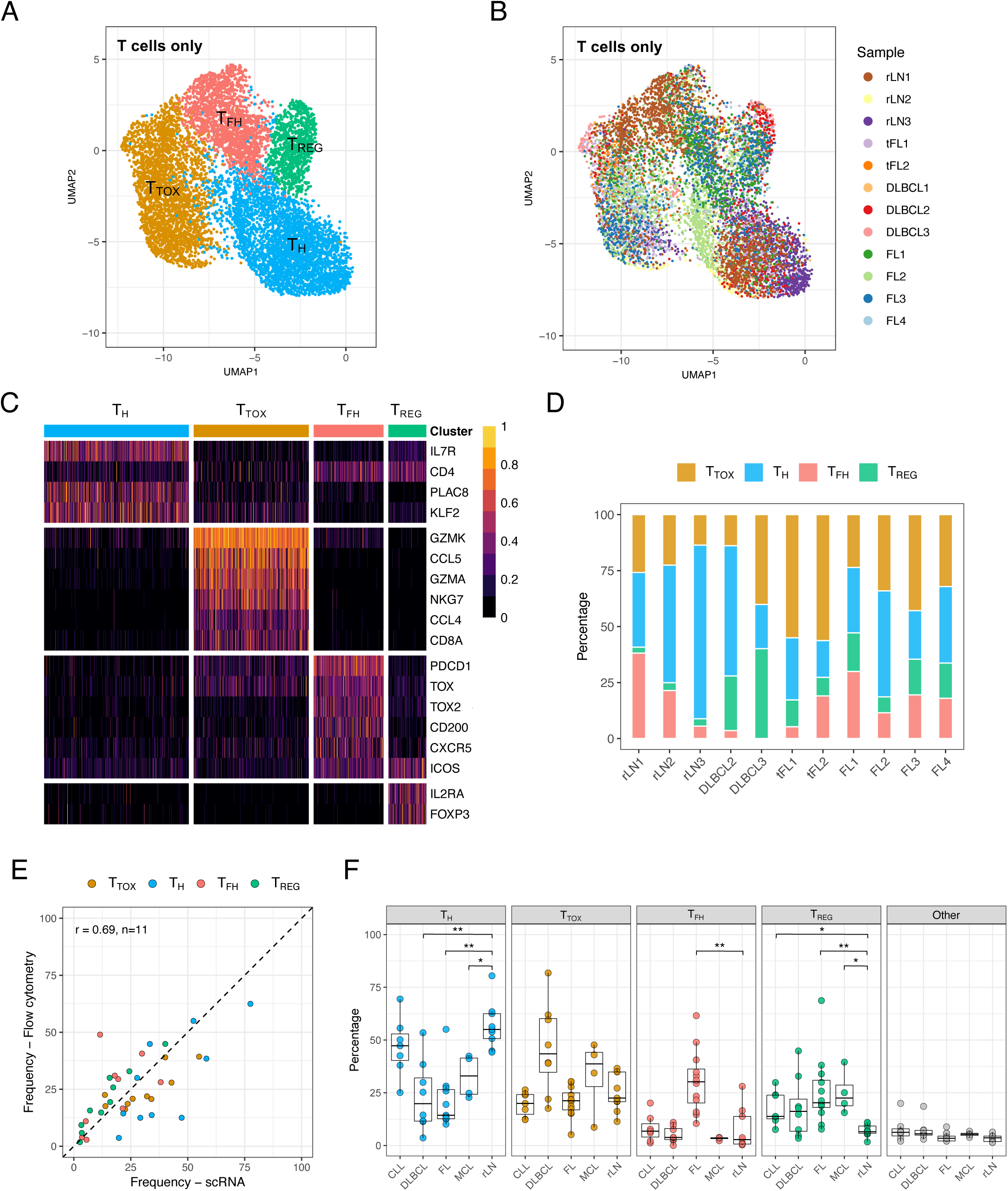
The transcriptional heterogeneity of lymph node-derived T cells. A and B) T cells from all samples were combined and jointly visualized using UMAP. Cells were colored with respect to their cluster of origin (A) or to their sample of origin (B). C) The heatmap shows differentially expressed genes, which were used to identify the T cell subsets: Cytotoxic T cells (T_TOX_), conventional T helper cells (T_H_), T follicular helper cells (T_FH_), regulatory T cells (T_REG_). Gene expression values were scaled to the the maximum of each row. D) Stacked bar chart displaying the proportion of T cell subpopulations based on scRNA-seq identified in each sample. Note that the DLBCL1 sample is not shown here due to only five T cells identified in this sample. E and F) Single cell suspensions of lymph nodes derived from 39 different patients, including those passed to scRNA-seq, were characterized by flow cytometry. The four different T cell populations identified by single cell RNA-Seq were distinguished using the following marker panel: CD3, CD4, CD8, PD1, ICOS and FoxP3. Specifically, T_H_ were identified based on CD3^+^CD4^+^ without the phenotype of T_FH_ or T_REG_; T_TOX_were identified based on CD3^+^CD8^+^; T_FH_ were identified based on CD3^+^CD4^+^ ICOS^High^PD1^High^; and T_REG_ were identified based on CD3^+^CD4^+^FoxP3^+^. E) The frequencies based on flow cytometry were correlated with the frequencies based on scRNA-seq. Pearson’s correlation coefficients (r) and the number of samples included (n) are given in the top left corner. Note that the DLBCL1 and tFL2 samples are not shown here due to the low number of T cells in the scRNA-seq data (DLBCL1) or the lack of material (tFL2). F) Frequencies for each subpopulation with regard to the sum of all T cells are shown. P values were calculated by the two-sided Wilcoxon’s test comparing each Entity with rLN group, and corrected by Bonferroni method. Only significant differences are shown; ** ≙ p value ≤ 0.01, * ≙ p value ≤ 0.05. SNN: Shared-nearest-neighbor-based. UMAP: Uniform Manifold Approximation and Projection. rLN: Reactive lymph node. MCL: Mantle cell lymphoma. FL: Follicular lymphoma. DLBCL: Diffuse large B cell lymphoma, CLL: Chronic lymphocytic leukemia.

To study this variation in a larger cohort, we quantified the abovementioned T cell populations in 39 lymph node samples of DLBCL, FL, mantle cell lymphoma and chronic lymphocytic leukemia by flow cytometry using the most distinctive markers (CD3, CD4, CD8, CD25, FoxP3, ICOS, PD1), as seen in Figure 2C. The frequencies of all T cell subsets derived from scRNA-seq correlated well with the frequencies determined by flow cytometry (r = 0.69, n = 10, Figure 2E). We found that T_FH_ cells were significantly increased in FL (two-sided Wilcoxon test: p = 0.006, Figure 2F), and T_REG_ cell frequencies were significantly increased in malignant lymph nodes, compared to the reactive ones (two-sided Wilcoxon test: p values as indicated, Figure 2F).

Taken together, we demonstrated that T cells derived from malignant B-NHL lymph nodes are transcriptionally similar to those derived from non-malignant reactive lymph nodes. In contrast, the proportion of individual T cell subsets differed significantly between lymphoma entities and individual patients. This finding indicates that B-NHL shape their microenvironment by influencing the recruitment of certain T cell subpopulations, but have less effect on their transcriptional programs. Therefore, studying the frequencies of lymphoma infiltrating T cell subsets and their effect on the outcome after immunotherapies might be highly relevant for the development of biomarkers.

### Identification of gene expression signatures driving B cell heterogeneity by scRNA-seq

Next, we examined the heterogeneity of the malignant and non-malignant B cells. To gain a global overview of the gene expression pattern across all malignant and non-malignant B cells from the 12 different donors, we combined their single cell RNA expression profiles, clustered them jointly and visualized them by UMAP (Figure 3A, B).

**Figure 3.**
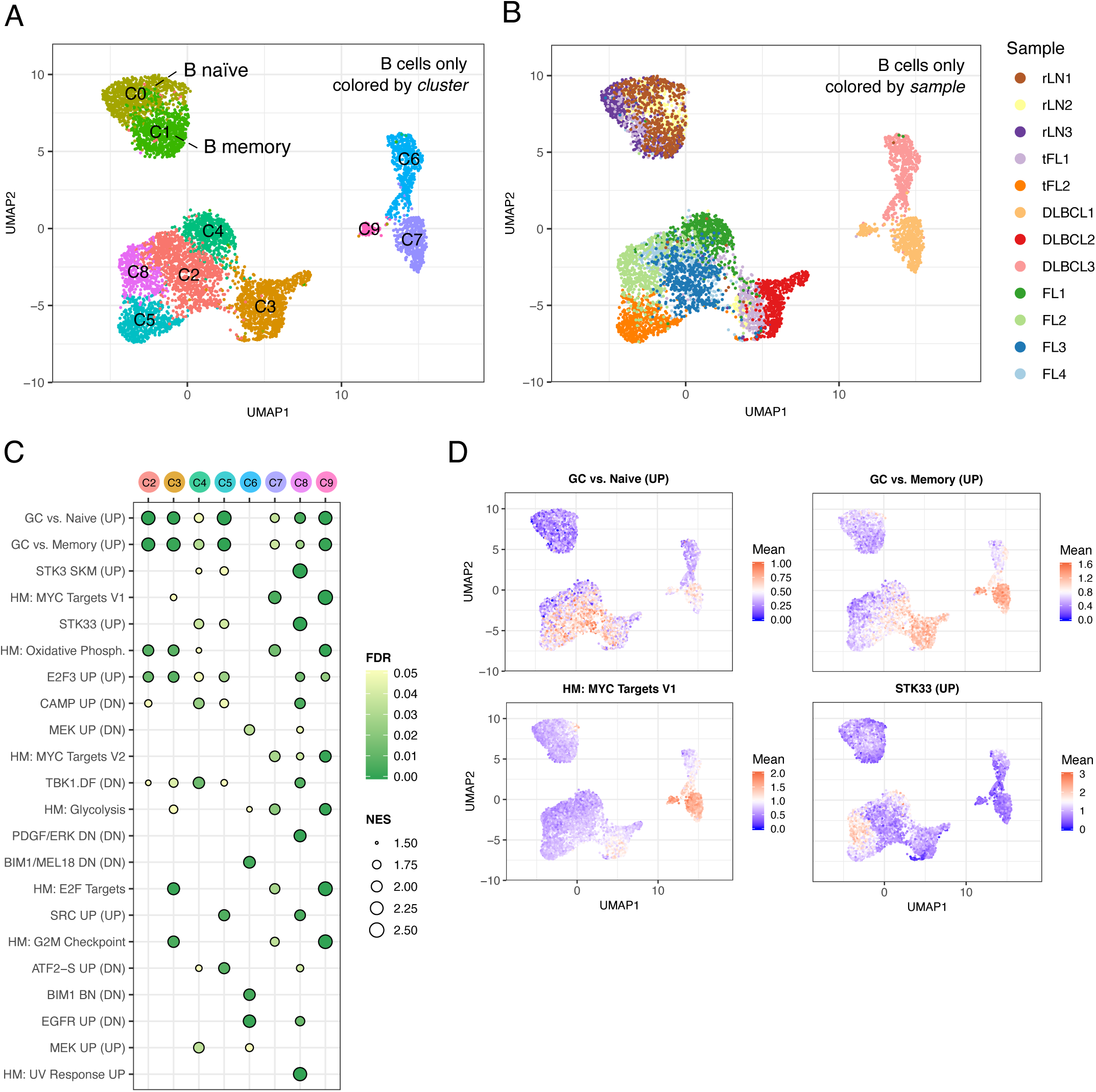
Gene expression signatures driving B cell heterogeneity. A and B) Single cell RNA expression profiles from all B cells were combined and jointly visualized using UMAP. Cells are colored either by SNN-based clusters (A) or by sample (B). C) A gene set enrichment analysis was performed separately for each malignant cluster (C3 to C9) versus all healthy B cells (C0, C1). The four most enriched gene sets per sample are shown. Columns refer to cluster. Circles are coded by color (nominal FDR) and size (NES). Gene sets with NES > 1.5 are shown. D) Cells in UMAP plot were colored by the mean expression of enriched genes for four representative gene expression signatures. UMAP: Uniform Manifold Approximation and Projection. SNN: Shared-nearest-neighbor. FDR: False-positive detection rate. NES: Normalized enrichment score.

Clustering partitioned the non-malignant B cells into two distinct subpopulations (C0-C1, Figure 3A). Among multiple differentially expressed genes between these two subsets (Supplementary Table 3), we found *IGHM* and *CD72* to be overexpressed in cluster C0, which characterizes naïve B cells [34], and *CD27* and *IGHG1* to be overexpressed in cluster C1, which characterizes memory B cells [35].

Of the eight transcriptionally distinct clusters formed by the malignant B cells (C2-C9, Figure 3A), six exclusively contained cells of only one donor (Figure 3A, B). This suggests a higher degree of inter-patient heterogeneity for malignant than for non-malignant B cells. We performed a gene set enrichment analysis (GSEA) on the mean expression differences between each malignant B cell cluster and all non-malignant cells, which revealed multiple cluster-specific gene sets (Figure 3C). Germinal center (GC)-associated gene expression signatures were significantly enriched in all clusters except for cluster 6, which exclusively contained malignant B cells of DLBCL3. This finding supports the classification of all B-NHL cells as either GCB type DLBCL or FL, except for the remaining DLBCL3 sample, which was classified as a non-GCB type DLBCL based on the Hans-classifier (Supplementary Table 1) [36]. Individual clusters were characterized by oncogenic transcriptional programs, which indicated activation of oncogenic MYC or STK33 signaling (Figure 3D).

Inter-patient heterogeneity of B cell lymphomas also comprises their proliferative capacity, which can vary from very low in FL to very high in DLBCL. We determined the proportion of B cells in S, G_2_ or M phase based on their single cell RNA profile (Supplementary Figure 4A) and observed a high correlation with flow cytometry- and IHC-based staining of Ki67 (R = 0.83 scRNA-seq to flow cytometry, R = 0.92, scRNA-seq to IHC, Supplementary Figure 4B).

In summary, these results indicate that inter-patient heterogeneity of malignant B cells, including their diverse proliferative activity, can be captured by the scRNA-Seq and can be linked to lymphoma-specific transcription signatures. Non-malignant B cells, however, had similar transcriptional profiles across different donors.

### Decoding the crosstalk between T cells and malignant B cells in the lymph node microenvironment

Above, we concluded that B cell lymphomas shape their microenvironment by modulating the frequency of different subsets of lymphoma infiltrating T cells. We now aimed to understand through which potential ligand-receptor interactions malignant B cells could benefit from their specific T cell microenvironment. For this purpose, we adopted a computational approach described by Vento-Tormo et al. [37] and analyzed 760 known ligand-receptor combinations (Supplementary Table 4) to identify the most significant interactions between malignant B cells and lymphoma infiltrating T cells within the lymph node microenvironment (Figure 4A).

**Figure 4.**
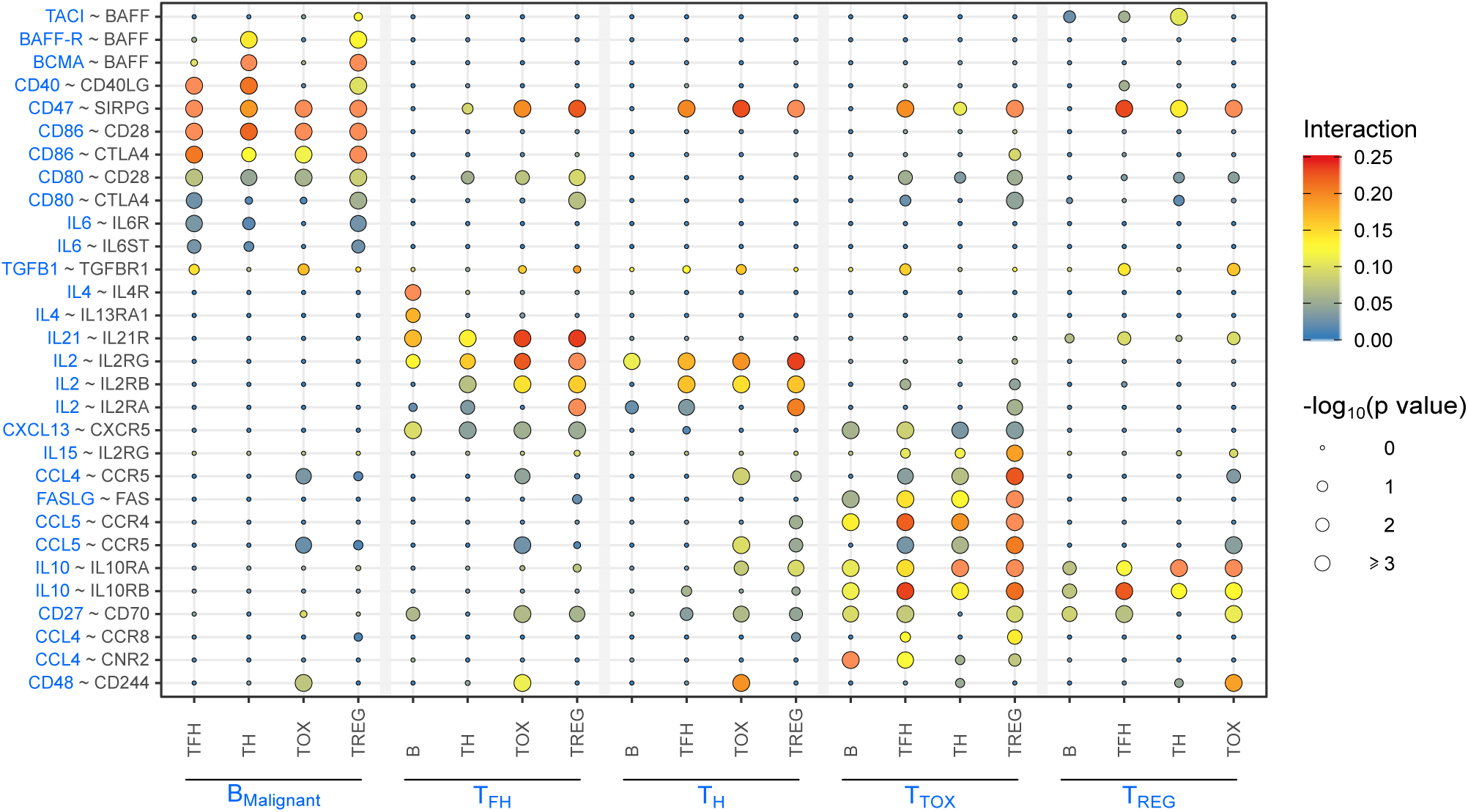
Cellular crosstalk in B cell lymphoma within the lymph node microenvironment. Overview of most significant ligand-receptor interactions across all lymphoma samples, excluding DLBCL1 due to the low number of T cells. Circle size indicates negative log_10_ of adjusted p values which were determined by permutation test (see Methods for details). Color scheme visualizes interaction scores which were calculated by the mean expression of molecule 1 (blue) in cell type A (blue) and the mean expression of molecule 2 (black) in cell type B (black). Protein names instead of gene names were used for TACI (*TNFRSF13B*), BAFF-R (*TNFRSF13C*), BCMA (*TNFRSF17*) and BAFF (*TNFSF13B*).

This analysis suggested that malignant B cells could receive costimulatory and coinhibitory signals by all four major T cell subsets, via CD80/CD86-CD28 and CD80/CD86-CTLA, while interactions via BCMA-BAFF, BAFF-R-BAFF and CD40-CD40LG could predominantly be mediated by T_H_ or T_REG_ cells. Significant interaction scores of IL4-IL4R and IL4-IL13RA1 were exclusively observed between T_FH_ and malignant B cells, providing further evidence that T_FH_ cells represent the most important source of IL4 production in B cell lymphoma [38]. This observation might be of clinical relevance because the IL4/IL4R interaction is discussed as potential resistance mechanism against Bruton’s tyrosine kinase (BTK) inhibitors [39, 40]. In line with the current state of knowledge [41–43], we also observed strong interaction scores for T_FH_ via IL21-IL21R with malignant B cells and via IL2-IL2R with other T cell subsets. This analysis supports the classification of T_FH_ cell as one of the four main T cell subsets within the lymph node microenvironment and reveals that each subset may provide a distinct panel of stimuli to interact with malignant B cells.

### Dissecting transcriptional intratumor heterogeneity using multicolor flow cytometry

Intratumor heterogeneity of nodal B cell lymphoma is a well-known phenomenon, however, most available studies infer intratumor heterogeneity from variant allele frequencies of genetic alterations corrected for purity, ploidy and multiplicity of local copy number [44, 45].

Here, we aimed to investigate the genomic, transcriptomic and functional (drug response) layers of intratumor heterogeneity from single cells. Unsupervised clustering of scRNA-seq profiles of malignant and non-malignant B cells revealed that all malignant samples were composed of at least two or more transcriptionally distinct subclusters (Supplementary Figure 5). We aimed to validate scRNA-based clusters at the cellular level to understand if this clustering represents biologically and clinically relevant differences. Therefore, we selected three samples (FL4, tFL1, DLBCL1) based on the availability of material for follow-up studies. We inferred differentially expressed surface markers from single cell expression profiles and first validated the distinction of scRNA-based clusters by flow cytometry. In a second step, we cultured lymph node derived lymphocytes with 58 different drugs in 5 concentrations (Supplementary Table 5) and stained them with specific antibody combinations to assess their drug response profiles by flow cytometry. In a third step, we sorted subpopulation and performed genome sequencing for each subclone (tFL1, DLBCL1).

### Verifying five transcriptionally distinct clusters in follicular lymphoma sample

The FL4 sample was collected at initial diagnosis. Based on single cell gene expression profiling, we identified five different B cell subpopulations (Supplementary Figure 6A). We aimed to validate all five clusters (C1 to C5) at the cellular level by flow cytometry and hence, we stained the differentially expressed surface markers CD44, CD24, CD22, CD27, kappa and lambda light chain (encoded by IGKC and IGLC2, Supplementary Figure 6B). Using the ratio of *IGKC* and *IGLC2* (see Methods for details), we found benign B cells in C1, lambda-restricted malignant B cells in C2, and malignant B cells with only marginal expression of *IGKC* and *IGLC2* in C3 to C5 (Supplementary Figure 6C). The pattern of light chain expression could be perfectly comprehended using flow cytometry (Supplementary Figure 6D), enabling us to differentiate C1 versus C2 versus C3, C4 and C5. Cluster C3 could then be recognized by a high expression of CD44 (Supplementary Figure 6D, 6E). To further distinguish C4 and C5 among the CD44^Low^ cells, we combined CD22, CD27 and CD24 and detected a subpopulation with CD22^High^, CD27^High^ and CD24^Low^, which corresponded to the expression pattern of cluster C5 (Supplementary Figure 6F). This approach allowed us to proof all five scRNA-based clusters by flow cytometry with comparable frequencies.

To assess subclone-specific drug response, we stained for kappa and lambda light chains and focused on the two major populations (C2 ≙ lambda^+^, C3-C5 ≙ kappa/lambda^-^). We did not observe differential responses for the majority of targeted drugs, but we found that only the kappa/lambda^-^ cluster was sensitive to chemotherapeutics (Supplementary Figure 6G). Interestingly, this patient received doxorubicin-based immunochemotherapy as first line treatment after sample collection and achieved only a partial remission.

### The indolent and aggressive component of transformed follicular lymphoma exhibit a distinct transcriptional, genomic and drug response profile

For the tFL1 sample, we detected three transcriptionally distinct clusters of B cells based on single cell RNA expression profiling (Figure 5A, B). Two clusters exclusively contained malignant B cells, and one cluster contained non-malignant B cells. We assessed the proliferative activity of both malignant populations based on their gene expression profiles, and observed that only one malignant cluster contained cells in S phase (Supplementary Figure 7A), with no cells in G_2_ or M phase (Supplementary Figure 7B). This suggests that this cluster represents a proliferating, thus aggressive component of the transformed FL. We performed a GSEA on the mean expression differences between the two malignant clusters, which revealed that gene expression signatures associated with MYC, MTORC1, and the G_2_M transition [46] were significantly enriched in the presumptively aggressive malignant B cell subclone (Supplementary Figure 7D-F).

**Figure 5.**
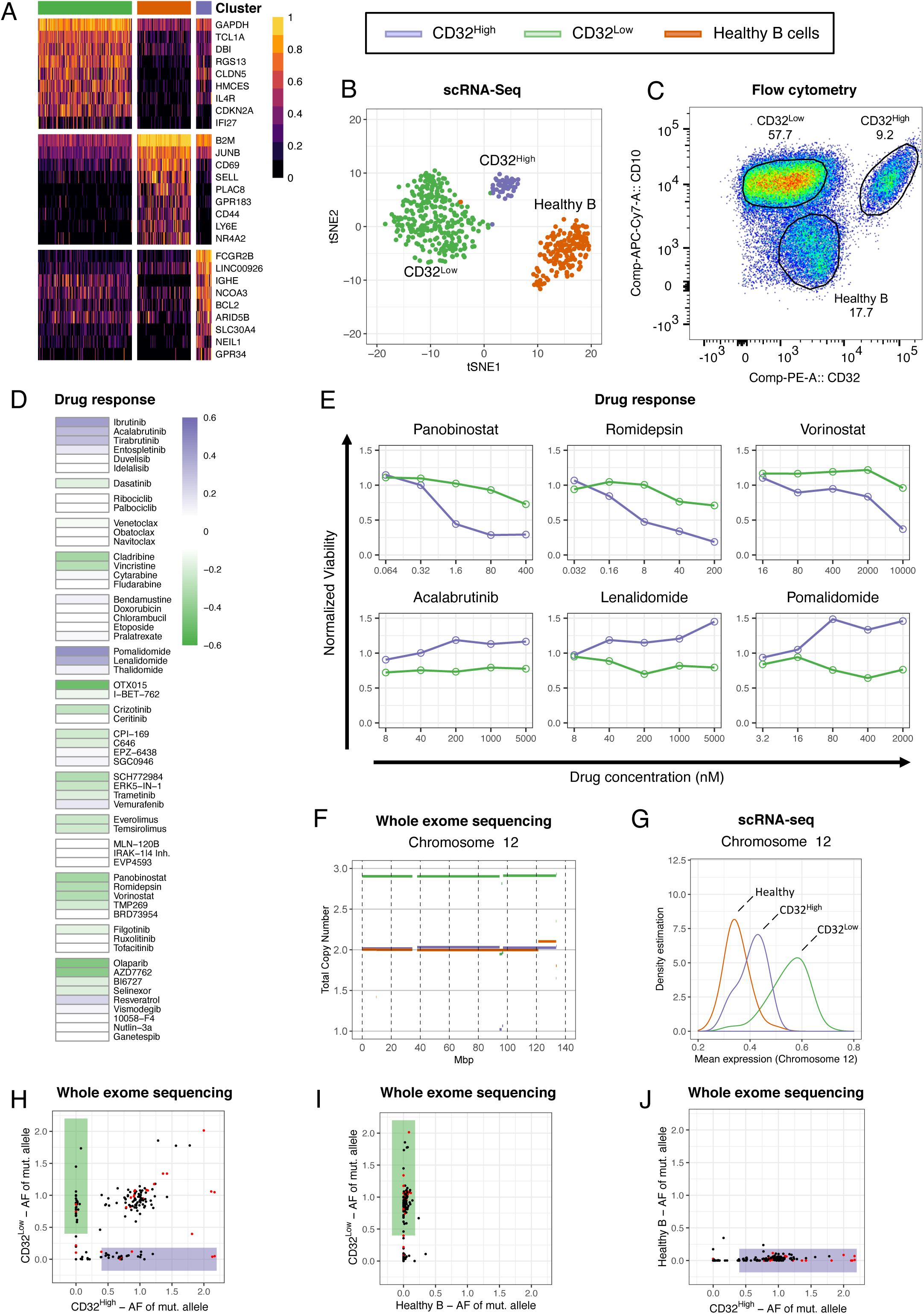
In depth-analysis of sample tFL1. A-B) Single cell RNA expression profiles of B cells derived from the tFL sample were subjected to SNN-based clustering. Three transcriptionally distinct clusters emerged. A) The heatmap illustrates the top 30 differentially expressed genes between all three identified clusters. Gene expression values were scaled to the maximum of each row. B) Clusters were colored and visualized in t-SNE projections of scRNA-seq expression profiles of malignant B cells. C) tFL1 derived lymph node cells were stained for viability, CD19, CD32, and CD10. The gates highlight three CD19^+^ populations which correspond to the subclusters shown in panel B. D and E) Unsorted single cell suspensions from the tFL sample were incubated for 48 hours with 58 different drugs and five concentrations, and stained as described in panel C. Viability was normalized to vehicle control for each subpopulation separately. D) The mean difference of viabilities between the two malignant subclones is shown. White indicates that both malignant clones responded equally to this drug. Purple or green indicates that the viability of the CD32^High^ or CD32^Low^ subpopulation was superior. E) Six representative subclone specific responses to the following drugs are shown: panobinostat, vorinostat, romidepsin (HDAC inhibitors), acalabrutinib (BCR signaling inhibitor), lenalidomide and pomalidomide (immunomodulatory imide drugs). F) Whole exome sequencing was performed on FACS sorted CD32^High^-, CD32^Low^- and the non-malignant CD10^-^ B cell subset. The line plot shows the total copy number estimation for chromosome 12 for all three sorted populations. The CD32^Low^ clone harbors an additional copy of chromosome 12. G) Density curves of single cell expression values for all genes located on chromosome 12 are shown for each subclone. H-J) The scatter plots show the allele frequency (AF) of the mutated allele for exonic SNVs in both malignant subclones (H), in CD32^Low^ versus healthy B cells (I) and in CD32^High^ versus healthy B cells (J). Shaded purple or green boxes highlight SNV that are exclusive to one of the malignant B cell subclones. Red dots mark immunoglobulin-associated mutations. SNN: Shared-nearest-neighbor. HDAC: Histone deacetylase. BCR: B cell receptor. SNV: Single nucleotide variant.

Among the genes differentially expressed between both subclones, we found *FCGR2B* (Figure 5A), which encodes a surface receptor protein (CD32B), to be exclusively expressed in the presumptively indolent subclone. Thus, we confirmed the existence of three B cell populations by flow cytometry (Figure 5C, see Supplementary Figure 8A for complete gating strategy). CD10 was strongly positive in both malignant B cell populations (CD32^High^, CD32^Low^), but not in non-malignant B cells.

As described above, we measured the ex vivo drug responses separately for each subclone (Figure 5D, E) and observed very different drug response profiles for the two malignant subclones. The BTK inhibitors, ibrutinib, acalabrutinib, and tirabrutinib, and the immunomodulatory imide drugs (pomalidomide, lenalidomide), were exclusively active in the CD32^Low^ subclone, whereas HDAC inhibitors (panobinostat, romidepsin, vorinostat) were more active in the CD32^High^ subclone.

Based on CD32 and CD10 expression, we sorted the three B cell subclones by flow cytometry (Supplementary Figure 8A) and performed WES of peripheral blood-derived normal control DNA, whole tumor DNA, DNA of both malignant subclones, and DNA of the non-malignant B cell population. Copy number profiles of both malignant subclones were very different, including exclusive aberrations of chromosomes 3, 4, 6, 10, 12, 15, 18 and X (Supplementary Figure 7F). Only the CD32^Low^ subclone harbored a trisomy 12 (Figure 5F), which was confirmed by scRNA-seq data (Figure 5G). Trisomy 12 has been associated with a better response to B cell receptor (BCR) signaling inhibitors [47], which was consistent with our observation that this subclone was more responsive to these drugs (Figure 5D, E). We also detected 157 somatic single nucleotide variants (SNV) in exonic regions, of which 25 (15.9 %) or 24 (15. 2%) were exclusively detected in the CD32^High^ or CD32^Low^ subclone, respectively (Figure 5H, I, Supplementary Table 6). However, the majority of somatic SNVs were equally represented in both subclones, indicating a phylogenetic relationship. We compared the allele count of all exonic SNVs between all three B cell populations and did not detect somatic SNV in healthy B cells (Figure 5J), which supports the validity of our sorting approach.

Taken together, scRNA-seq allowed us to identify different subclones within the same lymph node, which were genetically and functionally distinct in clinically-relevant aspects.

### A subclone-specific copy number variation of MYC drives a distinct gene expression and drug response program

The DLBCL1 sample was collected from a patient with a chemotherapy refractory disease during progression, but before retreatment. Using scRNA-seq, we identified two distinct clusters of malignant B cells, which exhibited a high number of differentially expressed genes associated with diverse cellular programs (Figure 6A, B), such as BCR signaling (*PRKCB*, *NFKB1I*), cytokine signaling (*LGALS9*, *IFITM1*), MAPK signaling (*RGS13*, *FBLN5*) and antigen processing (*PTPN22*, *SELL*, *CD48)*. Among the differentially expressed genes, we found *CD48* and *SELL* (Supplementary Figure 9A, B), which encode for the surface markers CD48 and CD62L respectively. Staining for CD48 and CD62L by flow cytometry validated the existence of the two distinct subclones (Figure 6C). The proportions of both clusters (CD48^High^CD62L^+^, CD48^Low^CD62L^-^) calculated based on flow cytometry and scRNA-seq were comparable, indicating good concordance between RNA and protein expression. We measured again the ex vivo drug responses for each subclone (Figure 6D, E) and observed a strikingly different drug response profile between the two subclones: B cell receptor (BCR) signaling inhibitors (acalabrutinib, tirabrutinib, ibutinib, duvelisib, idelalisib, entospletinib) and CDK inhibitors were exclusively effective in the CD48^Low^CD62L^-^ subclone, whereas Bromodomain and Extra-Terminal motif (BET) inhibitors (I-BET-762, OTX015), nucleoside analogues (cytarabine, fludarabine, cladribine) and vincristine were exclusively efficacious in the CD48^High^CD62L^+^ subclone.

**Figure 6.**
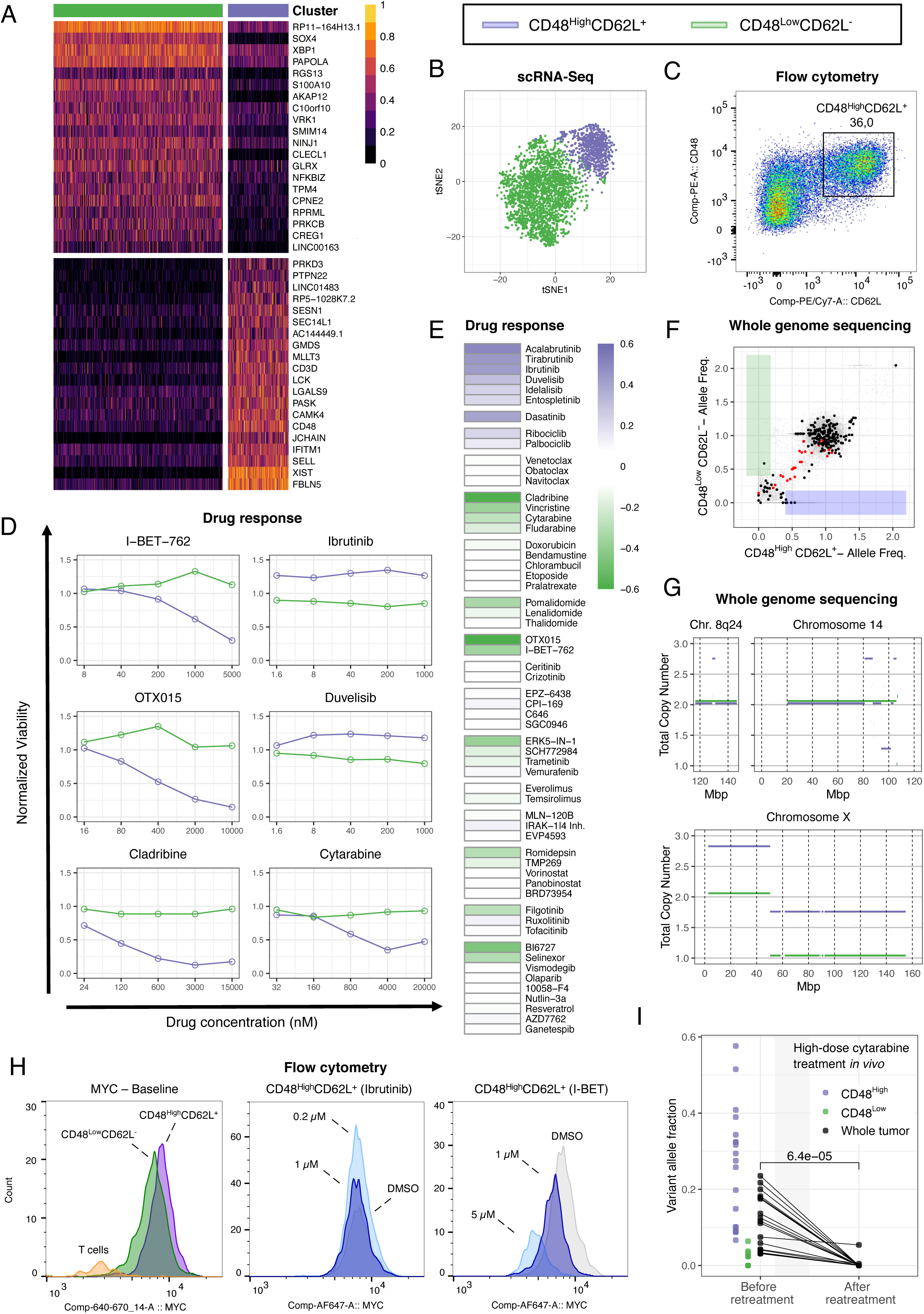
In depth-analysis of sample DLBCL1. A-B) Single cell RNA expression profiles of malignant B cells derived from the DLBCL1 sample were subjected to SNN-based clustering. Two transcriptionally distinct clusters emerged. A) Top 40 differentially expressed genes between the two identified clusters are shown in the heatmap. Gene expression values were scaled to the maximum of each row. B) Clusters were colored and visualized in t-SNE projections of scRNA-seq expression profiles of malignant B cells. C) DLBCL1 derived lymph node cells were stained for viability, CD19, CD48 and CD62L (=SELL). The gate highlights a population which co-expresses CD62L and CD48, which represents the identified subclones. D and E) Lymph node derived cells from the DLBCL1 sample were incubated for 48 hours with 58 different drugs and 5 concentrations. Cells were stained as described in C. Viability was normalized to DMSO controls for each subpopulation separately. D) Six representative subclone-specific responses to the following drugs are shown: I-BET-762, OTX015 (BET inhibitors), ibrutinib, duvelisib (BCR signaling inhibitors), cytarabine and cladribine (chemotherapy). E) The mean difference of viabilities between the two subpopulations is shown. White indicates that both clusters responded equally to this drug. Purple or green indicates that the viability of the CD48^High^CD62L^+^ or CD48^Low^CD62L^-^ subpopulation was superior. F-G) Whole genome sequencing was performed on both FACS sorted populations (CD48^High^CD62L^+^, CD48^Low^CD62L^-^). F) The scatter plot shows the allele frequency (AF) of the mutated allele for non-synonymous exonic SNV in bold black and synonymous or intronic SNV in faded grey of both subclones. Shaded purple or green boxes highlight SNV that are exclusive to one or the other subclone. Red dots mark immunoglobulin-associated non-synonymous exonic mutations. G) Line plots show total copy number estimations for chromosome 8q24, 14 and X for both clones. H) DLBCL1 derived lymph node cells were incubated with DMSO control, I-BET-762 at two concentrations (1 µM, 5 µM) or ibrutinib at two concentrations (0.2 µM, 1 µM). At baseline and after 24 hours cells were harvested and stained for viability, CD19, CD3, CD48, CD62L and MYC or respective isotype control. Histograms show the fluorescence intensity of MYC at baseline for T cells, CD48^High^CD62L^+^ and CD48^Low^CD62L^-^ subclone, after 24 hours incubation with I-BET-762 and DMSO control or Ibrutinib and DMSO control. I) Shown are SNV with high variant allele frequencies in the CD48^High^CD62L^+^ subpopulation (purple) and low or undetectable in CD48^Low^CD62L^-^ subpopulation (green). Black circles show corresponding variant allele frequencies of whole tumor samples before and after retreatment with high-dose cytarabine. P value was calculated by the paired Wilcoxon-text. SNN: Shared-nearest-neighbor. BET: Bromodomain and Extra-Terminal motif protein. BCR: B cell receptor. SNV: Single nucleotide variant. DMSO: Dimethyl sulfoxide.

We sorted viable tumor cells based on surface markers (CD48, CD62L, Supplementary Figure 8B) and performed WGS on each subclone separately, as well as on the whole tumor sample. In total, we detected 240 non-synonymous SNV located in exonic regions (Supplementary Table 7), however, only 1 (0.4 %) or 5 (2.1 %) SNV were exclusively detected in the CD48^Low^CD62L^-^ or the CD48^High^CD62L^+^ cluster, respectively (Figure 6F). We further compared CNV profiles of the two subclones and detected a number of differences: the CD48^High^CD62L^+^ cluster carried an additional copy of *MYC* (8q24, Figure 6G), which was reflected by increased *MYC* expression levels (Supplementary Figure 9C). The q arm of chromosome 14 harbored two copy number gains and one copy number loss in the CD48^High^CD62L^+^ cluster (Figure 6G). Moreover, chromosome X exhibited a copy number gain of the p arm in the CD48^High^CD62L^+^ cluster, and a copy number loss of the q arm in the CD48^Low^CD62L^-^ cluster (Figure 6G).

Since pathologic activation of MYC renders cells sensitive to BET inhibitors [48, 49], we performed intracellular flow cytometry-based staining of MYC at baseline and after 24 hours incubation with and without the two BET inhibitors, I-BET-762 or OTX015. We confirmed the increased MYC expression level of the CD48^High^CD62L^+^ subclone at baseline (Figure 6H, Supplementary Figure 9D), and, as expected, found that MYC was downregulated upon incubation with I-BET-762 and OTX015, but not upon incubation with the BTK inhibitor ibrutinib (Figure 6H, Supplementary Figure 9E-G).

### In vivo retreatment confirms ex vivo prediction of subpopulation-specific drug response

To exemplify the translational relevance of subclone-specific drug responses, we performed WES of DLBCL1 during the second relapse after retreatment with high-dose cytarabine. Based on ex vivo drug perturbation we had predicted that the CD48^High^CD62L^+^ but not the CD48^Low^CD62L^-^ subpopulation would respond to cytarabine (Figure 6D). We compared several synonymous SNV exclusive to the CD48^High^CD62L^+^ subpopulation before retreatment and during second relapse, and observed that the cytarabine-sensitive subpopulation was successfully eradicated (Figure 6I). Due to the lack of sufficiently exclusive SNV in the resistant subclone, we took advantage of the loss of heterozygosity (LOH) on chromosome Xq (Figure 6G) to determine the aberrant fraction of cells harboring a loss of Xq before and after retreatment. We found that the fraction of chemotherapy-resistant cells, harboring the loss of Xq, increased from 72 % to 93 % (see methods section for details).

In summary, we dissected the intratumor heterogeneity of the DLBCL1 sample on the transcriptional, genomic, and drug response level. This clinically relevant example highlights the huge translational relevance of tumor subpopulations and their specific drug response profile for personalized cancer treatment.

## Discussion

Intratumor heterogeneity poses a significant challenge for the clinical management of cancer patients. Advances of single cell technologies facilitated the profiling of intratumor heterogeneity at an unprecedented resolution [50]. Most of these studies comprehensively describe intratumor heterogeneity on the transcriptional level, but do not explore its functional consequences such as response or resistance to drugs. In this study, we address this limitation and identify transcriptionally distinct malignant subclones in B-NHL lymph node biopsies. We study differential drug response patterns of these subclones and genetic events which likely drove these differences.

Our analysis revealed the coexistence of up to four transcriptionally distinct subpopulations of malignant cells within individual B-NHL lymph node samples. This result recapitulates similar observations in follicular lymphoma [45], multiple myeloma [51] and other cancer entities [8, 12, 52]. We and others attributed this heterogeneity to differentially enriched gene sets, which indicate, for instance, activity of *MYC*, proliferation, or germinal center experience. However, we went further and established a straightforward strategy to prove the coexistence of up to four different tumor subpopulations at the cellular level. We subsequently performed perturbation assays with a comprehensive panel of clinically relevant drugs and observed that tumor subclones within the same lymph node responded strikingly different both to targeted compounds, such as ibrutinib, but also chemotherapeutics. The study by de Boer and colleagues supports our observation by demonstrating that acute myeloid leukemia subclones, which were identified on the basis of 50 leukemia-enriched plasma membrane proteins, had distinct functional properties including a differential sensitivity to FLT3-inhibition driven by a subclonal FLT3-ITD mutation [53]. Most preclinical in vitro and in vivo drug screens do not address such clonal heterogeneity, which may explain the failure of numerous drug candidates in the clinic [47]. For a single patient, we even demonstrated that the ex vivo drug response profiling correctly predicted the treatment sensitivity of tumor subclones in vivo. The prospective identification of rational combinations of cancer drugs that effectively target co-existing tumor subclones separately could avoid the outgrowth of resistant tumor clones under therapeutic pressure of a single drug, and would thereby improve efficacy of cancer treatments. Our study addresses this limitation of many ex vivo drug perturbation studies, and, due to its unbiased approach to prospectively dissect the malignant substructure, it is also generalizable to other cancer entities. However, due to the limitation of lymph node derived primary cells, we have to acknowledge that we could not apply our approach to all samples. Further studies are necessary to expand this approach and to address also spatial heterogeneity of malignant tumors.

Our approach enabled us to directly identify genetic factors that underlie the transcriptional and drug response differences between subclones. This distinguishes our work from a previous scRNA-seq study in FL, which indirectly compared allele frequencies of bulk WES with the size of transcriptionally distinct subclones [45]. The authors found a correlation between genomic alterations and subclonal fractions and concluded that somatic mutations are associated with transcriptional differences. These findings are in contrast to another study, which correlated subclusters derived from targeted single cell expression profiling of 91 genes with subclusters derived from single cell immunoglobulin heavy-chain (IGH) sequencing. In this study, the authors concluded that distinct gene expression clusters were not associated with subclones derived from IGH hypermutations [54]. While these studies provide only indirect evidence, we physically sorted tumor subclones and normal B cells, and performed WGS or WES separately for transcriptionally distinct lymphoma subpopulations. With regard to somatic mutations, we observed two different scenarios: in the DLBCL1 sample we identified almost no somatic SNVs to be exclusive for one or the other subclone, whereas in the tFL1 sample we found up to 15% exclusive somatic SNVs in each subclone. However, both examples represent scenarios where subclone specific drug profiles could not have been predicted by means of gene mutation sequencing. We further compared CNV profiles of the same tumor subclones, and found that all subclones harbored significantly different CNV profiles, suggesting that copy number alterations represent an important layer of genetic events which can drive differential gene expression programs and drug response profiles. Although our results support the general notion that genetic events drive subclone specific differences in drug response, they also highlight the difficulty to predict drug responses based on only genome sequencing in clinical practice. It might therefore be beneficial to obtain both genetic- and drug response profiles for personalized treatment decisions.

Exploring the heterogeneity of the immune microenvironment in B-NHL has the potential to better reveal how lymphomas shape their microenvironment and how lymphoma patients could be better stratified for the treatment with immunotherapies. T cells represented the largest non-malignant population in B NHL lymph node biopsies. We identified four major, transcriptionally distinct T cell subpopulations, which were annotated as cytotoxic T cells, regulatory T helper cells, conventional T helper cells and T follicular helper cells [29–33]. We measured the frequency of these T cell subsets in an extended cohort of malignant lymph node biopsies and found T follicular helper cells to be enriched in FL, which is in line with previous flow cytometry-based studies [33, 55]. These T cell subsets displayed only limited transcriptional heterogeneity with less variability between lymph nodes compared to malignant cells. However, the frequencies of these T cell subsets varied significantly across donors, which suggests that B-NHL shape their microenvironment by regulating the recruitment of different T cell subsets. This observation might be of clinical relevance, because cold tumors with very few infiltrating T cells have been reported to respond less well to immunotherapies [56].

Despite the rather small number of analyzed B-NHL patients, our study is of high clinical relevance. We demonstrated that the prospective identification of pre-existing transcriptionally distinct malignant subclones might be of diagnostic value to detect difficult to treat tumor subclones. In addition, our research establishes scRNA-seq as a new key technology for precise molecular profiling of relapsed and refractory nodal B cell lymphomas, and facilitates the design of new and molecularly-informed diagnosis and treatment strategies.

## Online Methods

### Patients samples and lymph node procession

Our study was approved by the Ethics Committee of the University of Heidelberg. Informed consent was obtained in advance. Immediately after the excision, the lymph node was cut in small pieces and put into Roswell Park Memorial Institute (Gibco) medium supplemented with 10 % fetal bovine serum (FBS, Gibco), penicillin and streptomycin (Gibco) at a final concentration of 100 U/ml and 100 µg/ml and L-Glutamine (Gibco) at a final concentration of 2 mM. After filtering by a 40 µm strainer, cells were washed once with phosphate-buffered saline (PBS, Gibco and put into RPMI medium (Gibco) medium supplemented with 20 % FBS (Gibco) and 10 % dimethyl sulfoxide (DMSO, Serva), and then cryopreserved in liquid nitrogen until further analysis. an

### Quantification of immunohistochemical staining

Formalin fixed lymph node tissue were processed through the hospital’s routine immunohistochemistry pipeline and thereby stained for CD3, PAX5 and Ki67 (all Ventana). After completion of diagnostics, the corresponding slides were scanned for a subset of patients (n = 7). To quantify the frequencies of B and T cells, the open source software QuPath (v0.1.2) was used for PAX5 or CD3 stained slides according to the recommended workflow [57]. After detection of about 100.000 cells per slide, the measurements were exported and further analyzed using R. We visualized the intracellular signal of diaminobenzidine staining of all detected events in a histogram. For the staining of PAX5 and CD3 we observed two clear peaks for all samples and set a threshold in between. Cells with an intracellular signal of CD3 or PAX5 greater than this threshold were regarded as T cells or B cells, respectively. The proportion of Ki67^+^ cells was obtained from routine pathology reports.

### Surface and intracellular staining by flow cytometry

As described above, lymph node derived cells were thawed and stained for viability using a fixable viability dye e506 (Thermo Fisher Scientific) and for different surface markers depending on the experimental setup. The following surface antibodies were used: anti-CD3-PerCP/Cy5.5, anti-CD3-APC, anti-CD19-BV421, anti-kappa-PE, anti-kappa-FITC, anti-lambda-PE/Dazzle, anti-CD22-APC, anti-CD24-BV785, anti-CD27-PE-Cy7, anti-CD32-PE, anti-CD44-PE, anti-CD48-PE, anti-CD62L-PE/Cy7, anti-CD10-APC-Cy7, anti-CD4-AF700, anti-CD8-FITC, anti-PD1-BV421 and anti-ICOS-PE/Dazzle (all Biolegend). In case of subsequent intracellular staining, cells were fixed and permeabilized with the intracellular fixation/permeabilization buffer set (Thermo Fisher Scientific) and stained with anti-MYC-AF647 (Thermo Fisher Scientific), anti-FoxP3-AF647 (BD Biosciences), or adequate isotype controls (Thermo Fisher Scientific, BD Biosciences). Cells were then analyzed with an LSR Fortessa (BD Biosciences) and FACSDiva (BD Biosciences, Version 8)

### Estimating the proportion of malignant and non-malignant B cells by flow cytometry

Staining for expression of the light chains (kappa, lambda) is a well-established tool to identify the accumulation of light chain restricted, malignant B cells [58]. Lymph node derived cells were stained as described above. In case of a kappa^+^ or lambda^+^ B cell population greater than 80 %, we regarded this population as light chain restricted and therefore as malignant. We further assumed that the ratio of kappa^+^ versus lambda^+^ B cells among the potentially remaining non-malignant B cells is still balanced. Therefore, there must be roughly the same proportion of non-malignant B cells among those carrying the restricted type of light chain. This ends up in the following formula to estimate the proportion of malignant cells:

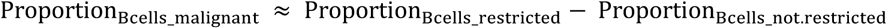

In addition, cells without detectable expression of kappa or lambda light chain on protein level were regarded as malignant cells because a loss of light chain expression is not observed in non-malignant lymph nodes [59].

### Single cell sample preparation and RNA sequencing

After thawing, cells were washed to remove DMSO as quickly as possible. We used the dead cell removal kit (Miltenyi Biotec) for all samples to achieve a viability of at least 90%. The preparation of the single cell suspensions, synthesis of cDNA and single cell libraries were performed using the Chromium single cell v2 3’ kit (10x Genomics) according to the manufacturer’s instructions. Each was sequenced on one NextSeq 550 lane (Illumina).

### Subclone specific drug screening

58 different drugs at 5 different concentrations (Supplementary Table 5) and a suitable number of DMSO controls were prepared in 384 well plates. DMSO concentration was kept equally at 0.2 % in all wells. Lymph node cells were thawed in a 37°C water bath and DMSO containing freezing medium was removed as quickly as possible to reduce cytotoxic effects. Afterwards, lymph node cells were rolled for 3 hours in RPMI medium supplemented with penicillin and streptomycin (Gibco) at a final concentration of 100 U/ml and 100 µg/ml, L-glutamine (Gibco) at a final concentration of 2 mM and with 10 % human AB male serum (Sigma). Cells were seeded at a cell count of 50,000 in 50 µl per well. After 48 hours, cells were washed once with staining buffer [PBS (Gibco) supplemented with 1% FBS and 0.5 % ethylenediaminetetraacetic acid (EDTA, Sigma Aldrich)]. Cells were subsequently stained with fixable viability dye e506 (Thermo Fisher Scientific), anti-CD3-APC, anti-CD19-BV421 and anti-CD48-PE, anti-CD62L-PE/Cy7 or anti-kappa-FITC, anti-lambda-PE/Dazzle, anti-CD10-APC/Cy7, anti-CD27-PE/Cy7, anti-CD32-PE (all Biolegend). After staining the microtiter plate was washed twice with staining buffer. Then, cells were fixed using paraformaldehyde at a final concentration of 2 % for 15 min at room temperature and washed with staining buffer. Fixed cells were analyzed with an LSR II and FACSDiva (BD Biosciences, Version 8) equipped with a high throughput sampler (HTS) system (BD Biosciences). Approximately 5,000 to 10,000 events were recorded per well. Flow cytometry data was analyzed using FlowJo software (Tree Star). The gating strategy is illustrated in Supplementary Figure 8. We ruled out that significant up- or downregulation of subclone-discriminating surface antigens confound subclone-specific drug response assessment by evaluating the fluorescence intensity of corresponding markers before and after drug treatment (Supplementary Figure 10).

### Fluorescence-activated cell sorting of B cell subclones

Lymph node cells were stained as described above. Sorting was performed at a FACS Aria Fusion (BD Biosciences). We sorted either for e506^−^ CD3^−^ CD19^+^ CD48^−^ CD62L^−^ and e506^−^ CD3^−^ CD19^+^ CD48^−^ CD62L^−^ (DLBCL1) or for e506^−^ CD3^−^ CD19^+^CD10^−^, e506^−^ CD3^−^ CD19^+^ CD10^+^ kappa^+^, CD32^low^ and e506^−^ CD3^−^ CD19^+^ CD10^+^ kappa^+^, CD32^high^ cells (tFL). The gating strategy is illustrated in Supplementary Figure 8. All relevant fractions were analyzed post-sorting to confirm a purity of at least 95 %.

### Whole genome and whole exome sequencing

DNA was extracted using the DNeasy mini kit (Qiagen) according to the manufacturers protocol, followed by quality control using gel electrophoresis and a TapeStation 2200 system (Agilent). Samples were prepared either for WGS or WES, as previously described [60]. Exome capturing was performed using SureSelect Human All Exon V5 in-solution capture reagents (Agilent). If samples were destined for WES on an Illumina HiSeq 2500 instrument, then 1.5 µg genomic DNA were fragmented to 150 to 200 bp insert size with a Covaris S2 device, and 250 ng of Illumina adapter-containing libraries were hybridized with exome baits at 65°C for 16 hours. If samples were destined for WES on an Illumina HiSeq 4000 instrument, then 200 ng genomic DNA were fragmented to 300 bp insert size with a Covaris LE220 or E220 device, and 750 ng of adapter-containing libraries were hybridized with exome baits at 65°C for 16 hours. If samples were destined for WGS on an Illumina HiSeq X instrument, then 100 ng of genomic DNA were fragmented to 450 bp insert size with a Covaris LE220 or E220 device, and libraries were prepared using the TruSeq Nano Kit (Illumina). On all platforms paired-end sequencing was carried out according to the manufacturer’s recommendations, yielding read lengths of 101 bp (4000) or 151 bp (HiSeq X).

### Single cell RNA sequencing data processing

The Cell Ranger analysis pipeline (v2.1, 10x Genomics) was used to demultiplex the raw base call files and to convert them into FASTQ files. FASTQ files were aligned to the reference genome (hg38) and filtered. Final numbers of cell barcodes, unique molecular identifiers (UMI) per cell, median genes and sequencing saturation are summarized in Supplementary Table 8.

### Filtering and normalizing single cell RNA sequencing data

The R package Seurat [61] (v2.3.3) was used to perform quality control and normalization. Gene count per cell, UMI count per cell and the percentage of mitochondrial and ribosomal transcripts were computed using the functions of the Seurat package. Genes expressed in three cells or fewer were excluded from downstream analysis. Libraries with a percentage of mitochondrial transcripts greater than 5%, along with those with less than 200 genes were filtered out prior to further analysis. Since aggressive lymphomas displayed higher gene and UMI count, the upper limit was set with regard to each sample. Counts were adjusted for cell-specific sampling (“normalized”) using the LogNormalize function with the default scale factor of 10,000.

### Assessing the cell cycle state using scRNA-seq data

The cell cycle state was assessed using the gene set and scoring system, described by Tirosh and colleagues [8]. Briefly, the S-Score and the G_2_M-Score were calculated based on a list of 43 S phase-specific and 54 G_2_ or M phase-specific genes. The calculation of the actual scores was performed using the CellCycleScoring function of the Seurat R package.

### Analysis of ligand-receptor interactions in scRNA-Seq data

We used the CellPhoneDB database [37] as basis for potential cell-cell interactions, but expanded the list by important B to T cell interactions (Supplementary Table 4). To assess the significance of each interaction, we adapted a statistical framework recently described by Vento-Tormo and colleagues [37] to our purpose. Importantly, we considered only genes which were expressed in 5 % of at least one cell type.

Briefly, we performed pairwise comparisons between the different T and B cell subtypes for each ligand-receptor pair and sample. For each combination of two different cell types and each ligand-receptor-pair, we permuted the cluster labels of cells at least 1,000 times and determined the mean interaction score (mean expression of ligand in cell type A times mean expression of receptor in cell type B). A p value was determined by calculating the proportion of permuted interaction scores which were by hazard higher than the actual interaction score. All interactions were calculated sample-wise. To determine which interactions were most relevant across different samples, we calculated the mean interaction scores and combined the different p values using the Fisher’s method. Then, p values were corrected using the Benjamini-Hochberg method. The R code is available on our GitHub repository (see code availability statement below).

### Combining data from different samples and batch correction

After identification of the different cell types the data sets were split into non-B cells or B cells using the SubsetData function. Then the respective subsets were combined using the MergeSeurat function. Putative batch effects between two runs, were corrected by the mutual nearest neighbors (MNN) technique [62] which is implemented in the scran Bioconductor package (v1.10.2).

### Clustering and dimensionality reduction techniques

SNN (Shared-nearest neighbor)-based clustering, t-SNE and UMAP visualization were performed using the FindClusters, RunTSNE and RunUMAP functions within the Seurat package [61]. Each of these were performed on the basis of a principal component analysis which was performed using the RunPCA function of the Seurat package. The same parameters were applied to all samples. UMAP was used instead of t-SNE for combined data sets because it is significantly faster than t-SNE and better preserves aspects of global structure in larger data sets [28]. Differentially expressed genes between the clusters were identified using the FindMarkers or FindAllMarkers functions within the Seurat package [61]. Differentially expressed genes between malignant B cell clusters can be browsed interactively using an html file (see data sharing statement below).

### Gene set enrichment analysis

Gene set enrichment analysis (GSEA) was performed using the GSEA java desktop application [63, 64] and the Molecular Signatures Database (MSigDB, v6.2) provided by the Broad Institute [63, 65]. Differentially expressed genes of two groups were used to determine significantly-enriched gene sets.

### WES and WGS data processing

Alignment of sequencing read pairs and variant calling were performed as recently described [66]. Briefly, reads were mapped to human reference genome (hg19) with bwa-mem (version 0.7.8, minimum base quality threshold set to zero [-T 0], remaining settings left to default) [67]. Subsequently, reads were coordinate-sorted with bamsort (compression option set to fast) and duplicate read pairs were marked with bammarkduplicates (compression option set to best) (both part of biobambam package version 0.0.148).

SNV and indels in matched tumor normal pairs were identified using the internal DKFZ variant calling workflows based on samtools/bcftools 0.1.19 with additional custom filters (optimized for somatic variant calling by deactivating the pval-threshold in bcftools) and Platypus 0.8.1, respectively, as described previously [66]. Gene annotation of variants was done with Annovar [68]. The variants were annotated with dbSNP141, 1000 Genomes (phase 1), Gencode mapability track, UCSC High Seq Depth track, UCSC Simple-Tandem repeats, UCSC Repeat-Masker, DUKE-Excluded, DAC-Blacklist, UCSC Selfchain. These annotation tracks were used to determine a confidence score for each variant by a heuristic punishment scheme and only high confidence variants were kept for further analysis. In addition, variants with strong read biases according to the strand bias filter were removed.

Genomic structural rearrangements (SVs) were identified using the SOPHIA algorithm (unpublished, source code available at https://bitbucket.org/utoprak/sophia/). Briefly, supplementary alignments as produced by bwa-mem are used as indicators of potential underlying SVs. Candidates are filtered by comparing them to a background control set of sequencing data obtained using normal blood samples from a background population database of 3261 patients from published TCGA and ICGC studies as well as published and unpublished studies of the German Cancer Research Center (DKFZ).

Allele-specific CNV were detected using ACEseq (allele-specific copy number estimation from WGS) [69] for WGS data and CNVkit for WES data [70]. ACEseq determines absolute allele-specific copy numbers as well as tumor ploidy and tumor cell content based on coverage ratios of tumor and control as well as the B-allele frequency (BAF) of heterozygous single-nucleotide polymorphisms (SNPs). SVs called by SOPHIA were incorporated to improve genome segmentation.

### Multi tumor comparison

To compare multi tumor samples of the same donor, every SNV position in each sample was determined using samtools mpileup 1.6. At each of these SNV positions, the variant allele fraction was determined by calculating the ratio between the number of variant reads and the total coverage at that position. To correct the variant allele fraction for actual tumor cell content, a scaling factor was incorporated, comprising ploidy and total copy number (TCN) estimates obtained from ACEseq/CNVkit. Specifically, the scaling factor is obtained as the ratio between purity corrected number of alleles in the tumor (TCN_tumor purity_tumor) and purity corrected total number of alleles in the sample ((TCN_tumor * purity_tumor) + 2 * (1 - purity_tumor)).

### Aberrant cell fraction estimation from LOH

To determine aberrant cell fractions, the minor allele-frequency (MAF, ratio between number of reads of minor allele and total coverage at given position) of single nucleotide polymorphism (SNP) was estimated for selected regions harboring a loss of heterozygosity (LOH) or a copy number neutral LOH (CN-LOH) in the tumor sample. Information on SNP location was received from matched-control SNV calling. To select heterozygous SNP, only SNP with a MAF ≥ 0.3 in the control were retained. Subsequently, MAF values of the selected SNP were calculated for the tumor samples. For exome samples, only SNP within the targeted capture regions were kept. The mean of the respective tumor MAF values was calculated and the aberrant cell fraction (ACF) was estimated as follows:

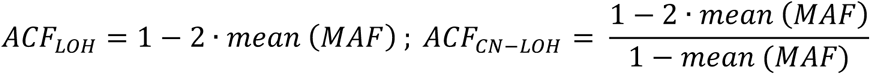

### Data sharing statement

The single cell expression data of merged B and T cell UMAP plots (Figure 2A/B and Figure 3A/B) are available for easy-to-use interactive browsing: https://www.zmbh.uni-heidelberg.de/Anders/scLN-index.html.

The raw single cell count tables can be downloaded here doi.org/10.11588/data/VRJUNV. This link will be activated upon publication and is accessible without further restriction. Differentially expressed genes between B cell clusters can be browsed in an interactive html file (Supplementary File 1).

### Code availability statement

R codes used for data analysis are available at our GitHub repository without further restriction (www.github.com/DietrichLab/scLymphomaExplorer).

## Acknowledgements

T.R. was supported by a physician scientist fellowship of the Medical Faculty of University Heidelberg. M.Se. was supported by a grant of the Deutsche Forschungsgemeinschaft (DFG). S.D. was supported by a grant of the Hairy Cell Leukemia Foundation, the Heidelberg Research Centre for Molecular Medicine (HRCMM) and an e:med BMBF junior group grant. For the data management we thank the Scientific Data Storage Heidelberg (SDS@hd) which is funded by the state of Baden-Württemberg and a DFG grant (INST 35/1314-1 FUGG). We thank Carolin Kolb (University Hospital Heidelberg) and Mareike Knoll (German Cancer Research Center Heidelberg) for their excellent technical assistance. We also thank the DKFZ Single-Cell Open Lab (scOpenLab) for the experimental assistance in terms of scRNA-seq. Also, this study was supported by the Heidelberg Center for Personalized Oncology (DKFZ-HIPO). We thank the DKFZ Omics IT and Data Management Core Facility (ODCF) and the DKFZ Genomics and Proteomics Core Facility (GPCF) for their technical support.

