## Supplementary Figures for "Dissecting intratumor heterogeneity of nodal B cell lymphomas on the transcriptional, genetic, and drug response level"

### Supplementary Figure 1

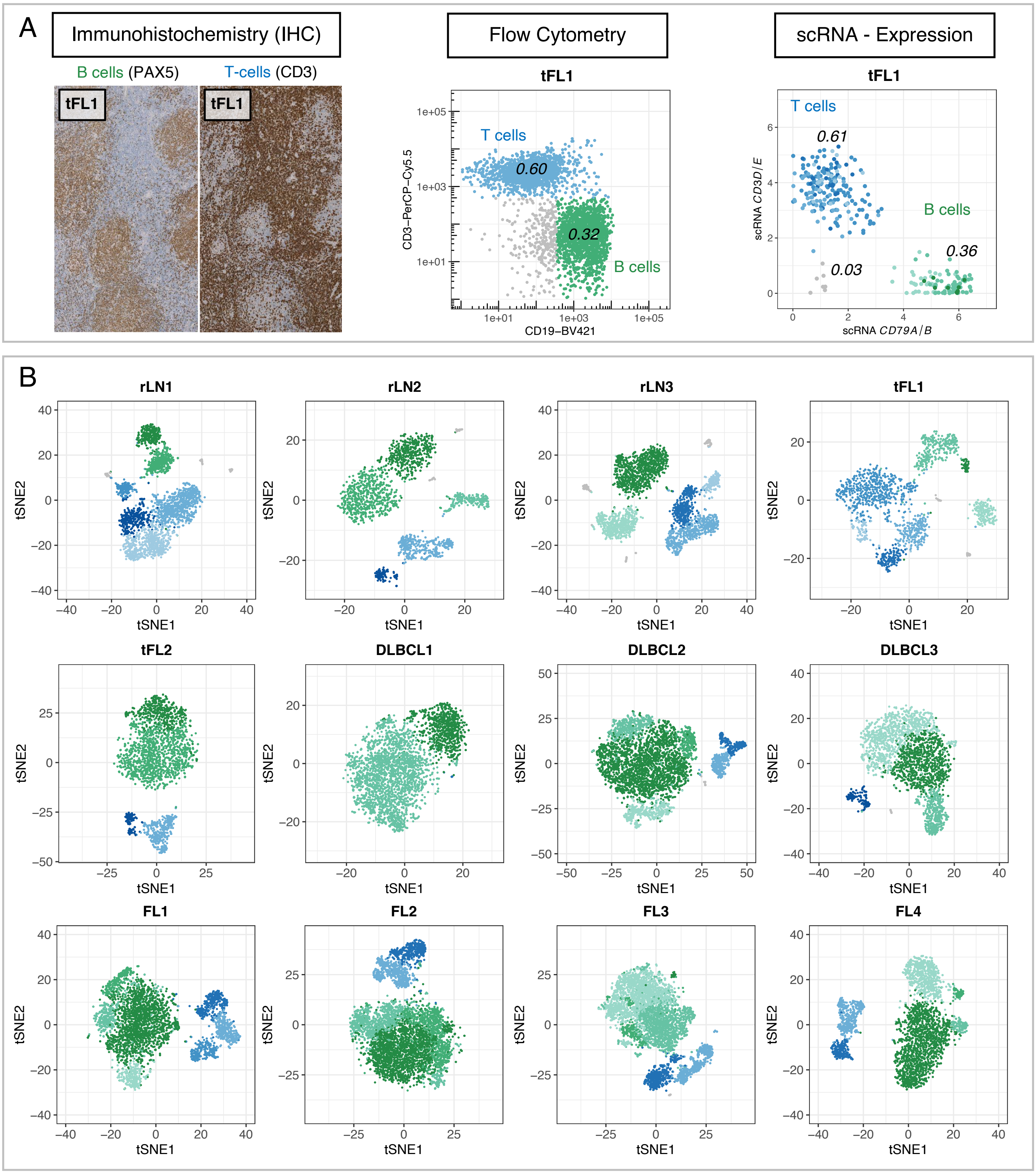


**Supplementary Figure 1.**

A) Example lymph node sample demonstrating how lymph node-derived B and T cells were classified by scRNA-seq (*CD79B* vs. *CD3*), flow cytometry (CD19 vs. CD3), or immunohistochemistry (PAX5 vs. CD3). Frequencies of B and T cells were calculated for each approach and correlated pairwise with each other (main Figure 1B and Figure 1C). B) Cell populations for all lymph node samples visualized in t-SNE plots. scRNA-seq data of all cells were subjected to SNN-based clustering and visualized by t-SNE. Each t-SNE represents one individual sample as indicated. SNN: Shared-nearest-neighbor.

### Supplementary Figure 2

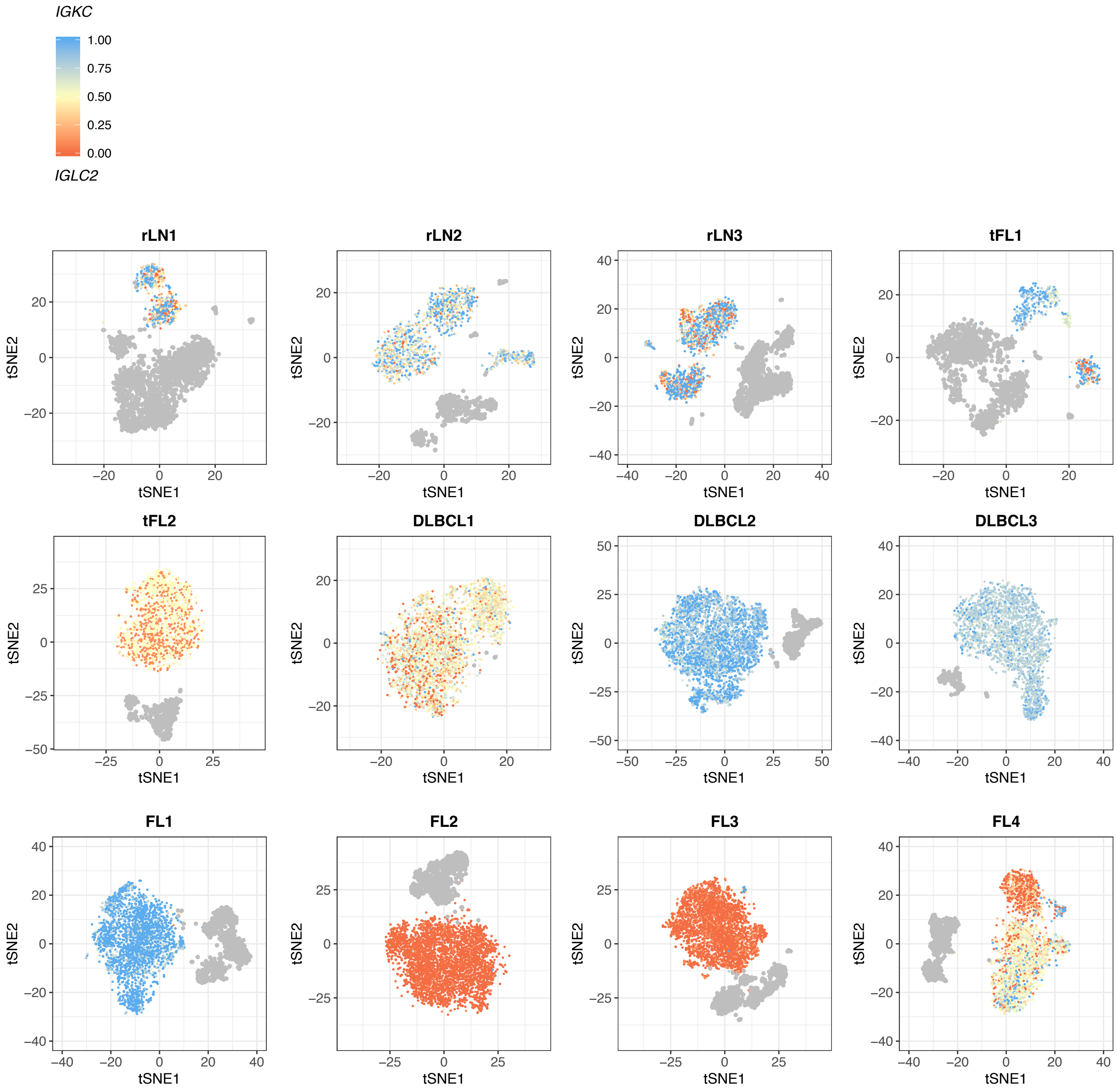


**Supplementary Figure 2.**

Cell populations for all lymph node samples visualized in t-SNE plots. scRNA-seq data of all cells were subjected to SNN-based clustering and visualized by t-SNE. Each t-SNE represents one individual sample as indicated. Cells are colored by light chain kappa fraction *IGKC*/(*IGKC*+*IGLC2)* of each single cell to demonstrate light chain restriction of each cluster. Non-B cells are colored in grey. The sample DLBCL1 showed only marginal light chain expression on single cell RNA level. Therefore, these cells were regarded as malignant cells (see Method section for details). The same is true for the larger cluster of sample FL4. SNN: Shared-nearest-neighbor.

### Supplementary Figure 3

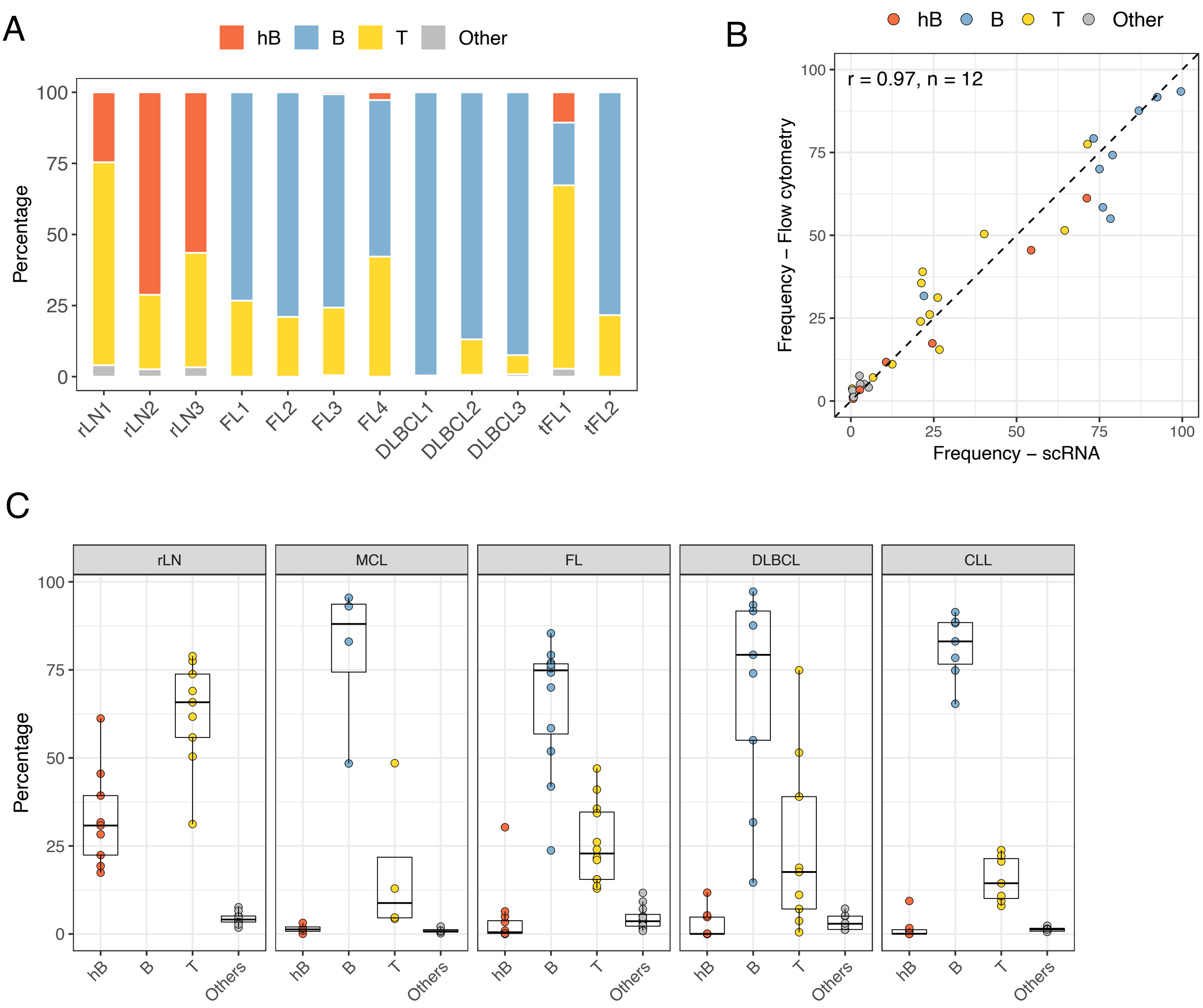


**Supplementary Figure 3.**

A) Stacked bar chart of the relative frequencies of malignant B cells (B), T cells (T), healthy B cells (hB) and myeloid cells (Other) for each sample calculated on the basis of single RNA expression profiles. B) Lymph node derived cells derived from those samples passed to scRNA-seq were stained for viability, CD19, CD3, kappa light chain and lambda light chain. The proportion of healthy (hB) and malignant B cells (B) were estimated based on the ratio of light chain restricted $CD19^+$ tumor cells and $CD19^+$ non-tumor cells (malignant B cells = light chain restricted B cells – non-restricted B cells). T cells (T) refer to $CD3^+CD19^-$ cells, whereas Other refers to $CD19^-CD3^-$ double negative cells. Pearson's correlation coefficients (r) and the number of samples included (n) are given in the left top corner. C) Lymph node derived cells from 40 different patients were stained for viability, CD19, CD3, kappa light chain and lambda light chain. The proportion of healthy and malignant B cells were estimated based on the ratio of light chain restricted $CD19^+$ tumor cells and $CD19^+$ non-tumor cells (malignant B cells = light chain restricted B cells – non-restricted B cells). T cells (T) refers to $CD3^+CD19^-$ cells, whereas Other refers to $CD19^-CD3^-$ double negative cells. rLN: Reactive lymph node. MCL: Mantle cell lymphoma. FL: Follicular lymphoma. DLBCL: Diffuse large B cell lymphoma, CLL: Chronic lymphocytic leukemia.

### Supplementary Figure 4

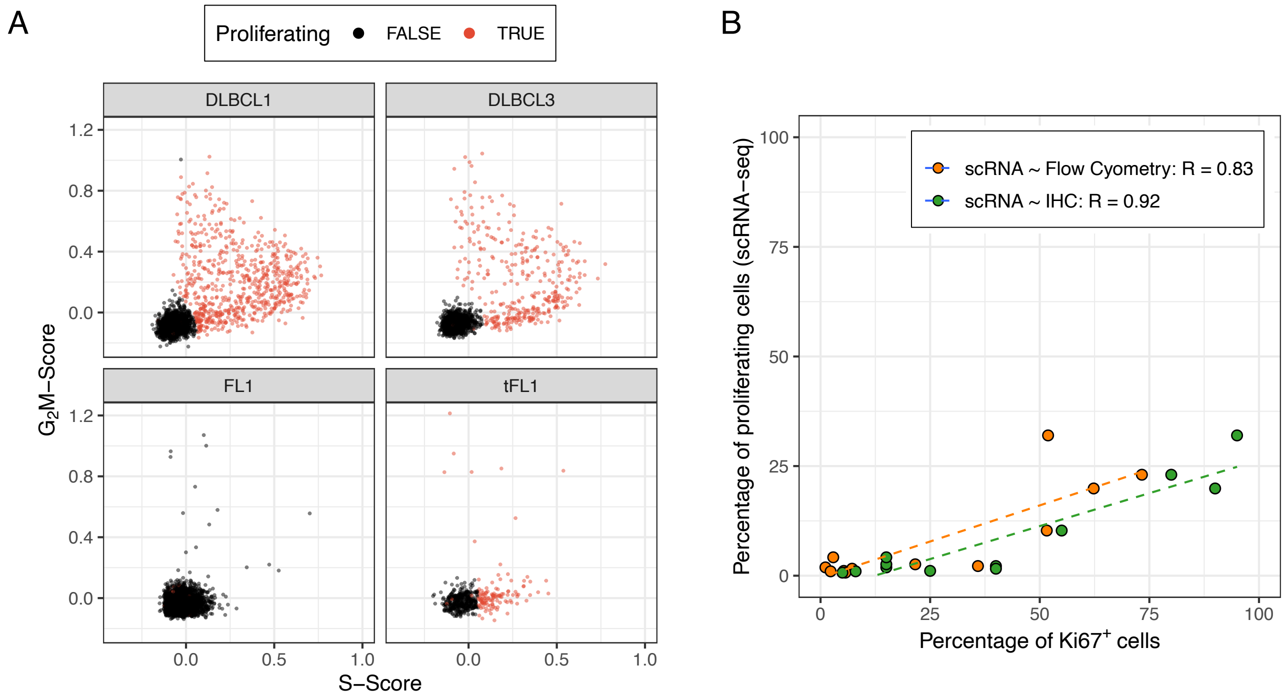


**Supplementary Figure 4.**

A) Dot plots show $G_2M$ and S score for the B cells of four representative samples. Cells with positive $G_2M$ or S score were marked as proliferating (please see method section for details). B) The proportion of proliferating cells based on scRNA-seq (panel A) was correlated with the percentage of $Ki67^+$ cells determined either by flow cytometry (orange) or immunohistochemistry (green). R values represent Pearson correlation coefficients.

### Supplementary Figure 5

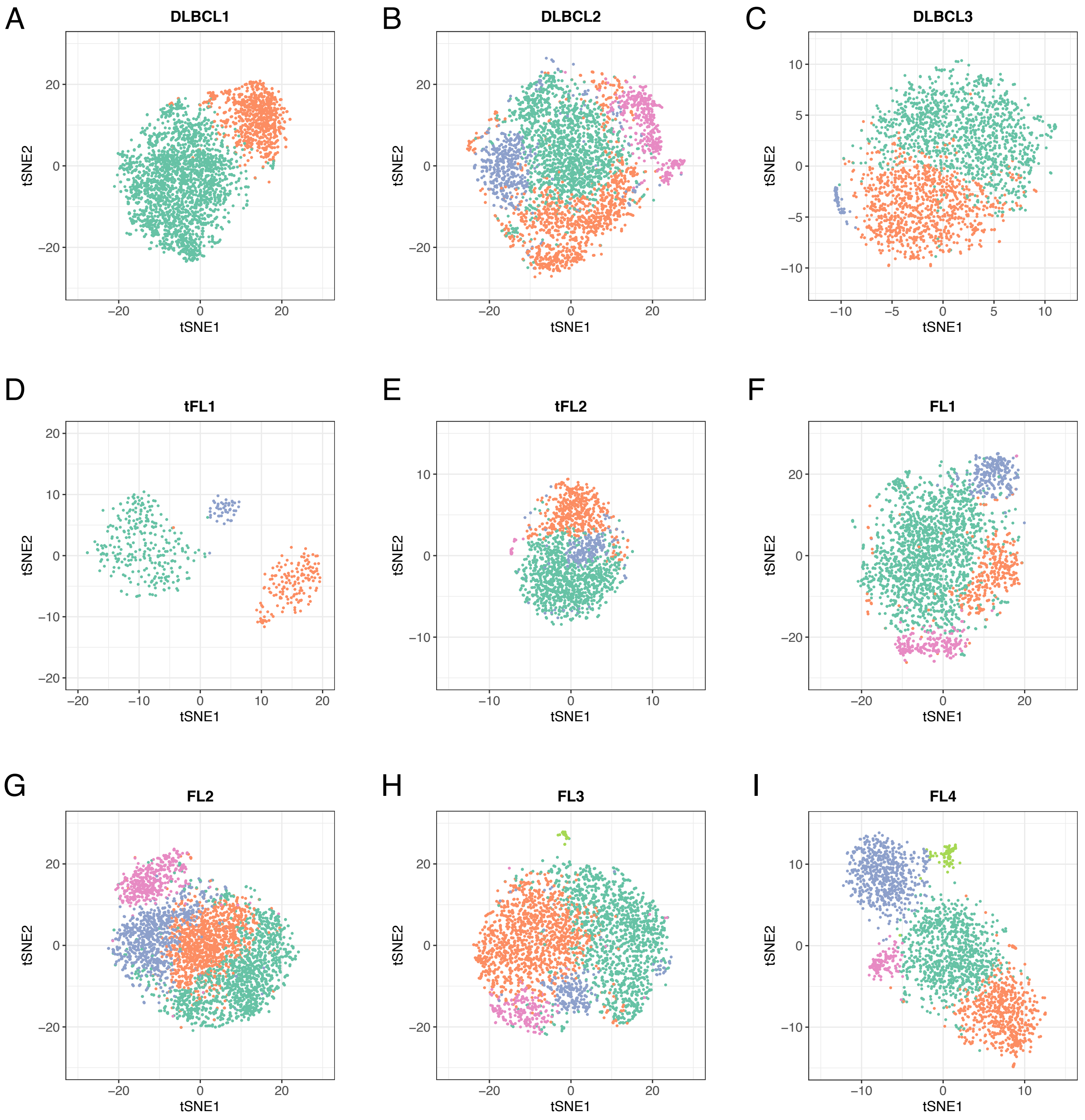


**Supplementary Figure 5.**

A-F) scRNA-seq data of malignant and non-malignant B cells only were subjected to SNN-based clustering and visualized by t-SNE. Each t-SNE represents one individual sample as indicated. Cells were colored by cluster. SNN: Shared-nearest-neighbor.

### Supplementary Figure 6

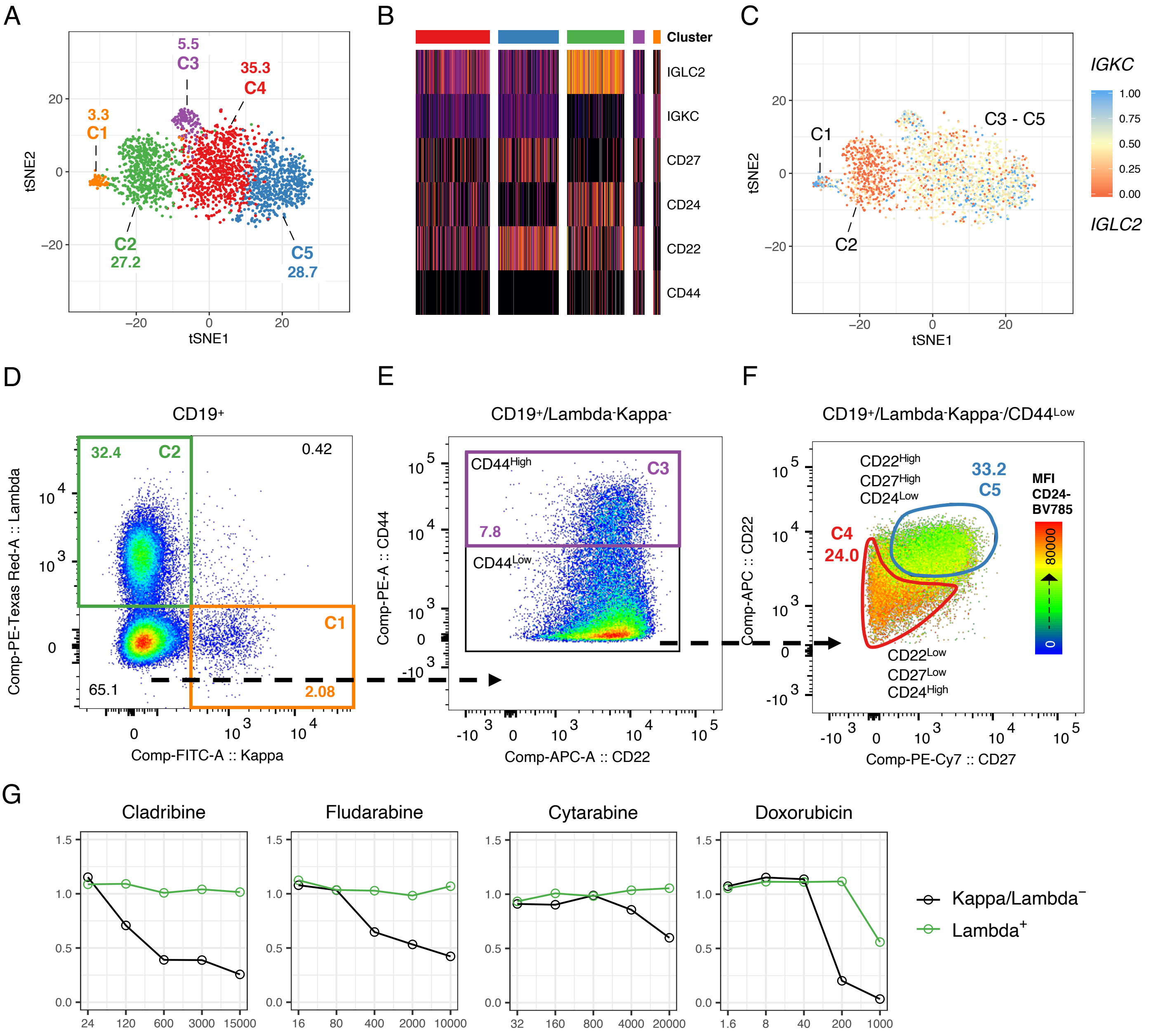


**Supplementary Figure 6.**

A) scRNA-seq data of B cells from the FL4 sample were subjected to SNN-based clustering and visualized by t-SNE. B) Heatmap shows a selection differentially expressed surface markers used for cluster differentiation. Gene expression values were scaled to the maximum of each row. C) T-SNE plot of panel A colored by the light chain kappa fraction *IGKC*/(*IGKC*+*IGLC2)* of each single cell. C1 contains cells either expressing *IGKC* or *IGLC2* predominantly (benign B cells), C2 contains only cells expressing predominantly *IGLC2*, whereas C3 to C5 hardly express both *IGKC* and *IGLC2.* D-F) Cells derived from sample FL4 were stained for viability, CD3, CD19, kappa, lambda, CD44, CD24, CD22 and CD27. Shown is the stepwise gating strategy to comprehend the scRNA-Seq-based clusters of panel A. G) Lymph node derived cells from the FL4 sample were incubated for 48 hours with 58 different drugs and 5 concentrations. Cells were stained for viability, CD3, CD19, kappa, lambda and CD27. Viability was normalized to DMSO controls for each subpopulation separately. Shows are only those drugs with significantly differential drug response between the two subpopulations. SNN: Shared-nearest-neighbor. DMSO: Dimethyl sulfoxide.

### Supplementary Figure 7

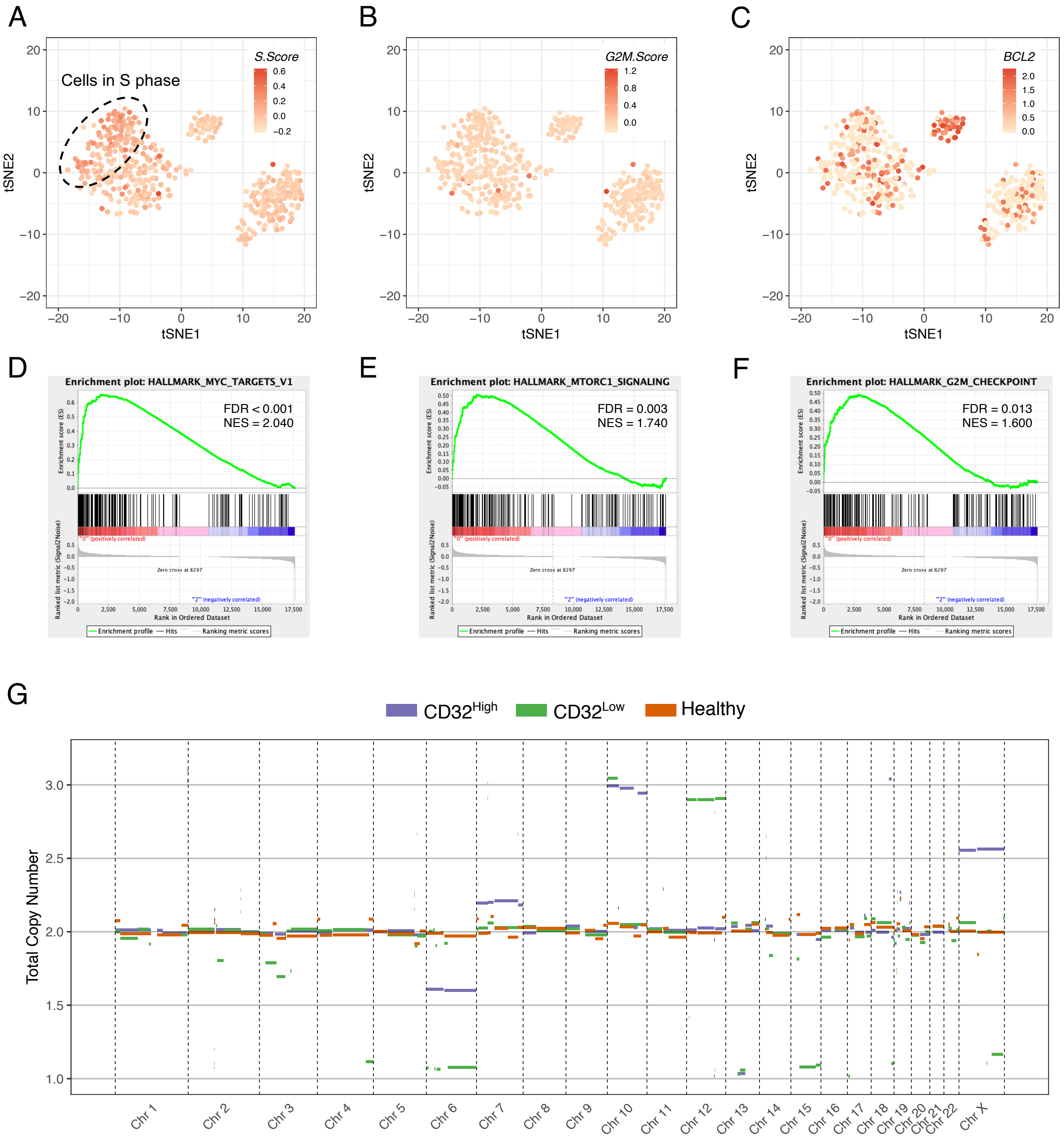


**Supplementary Figure 7.**

A-B) Single cell RNA expression profiles of B cells only (tFL1) were subjected to t-SNE and colored by S-Score (A, see Methods section for details), $G_2$M-Score (B, see Methods section for details) or *BCL2* expression (C). D-F) Gene set enrichment analysis was performed for each lymphoma sample versus all healthy B cells. Shown are enrichment plots for hallmark MYC targets (D), MTORC1 signaling E) and hallmark $G_2$M targets (F). Given are false-positive detection rate (FDR) and normalized enrichment score (NES). G) Line plot shows total cluster-specific copy number estimation for all chromosomes as inferred from whole exome sequencing.

### Supplementary Figure 8

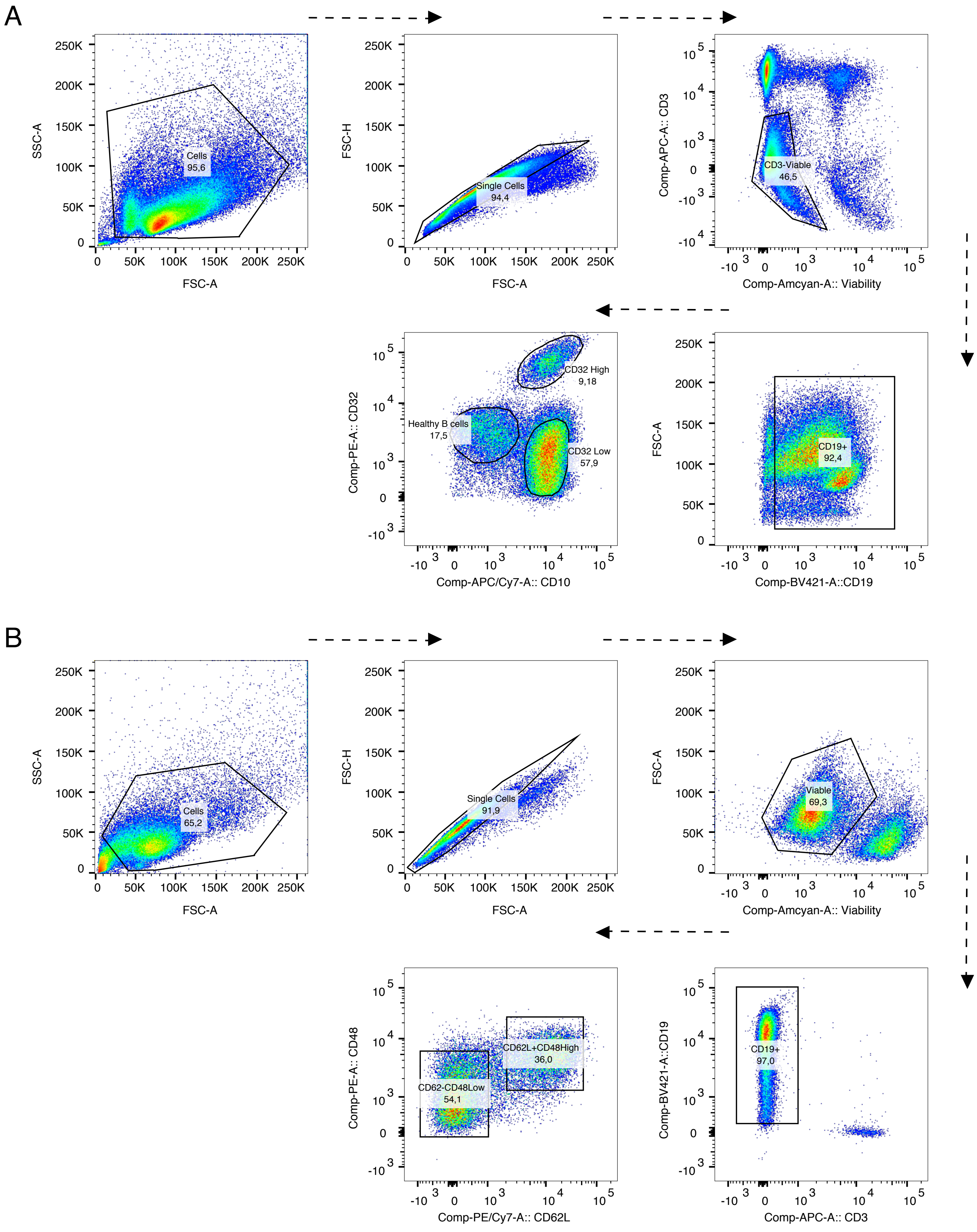


**Supplementary Figure 8.**

Pseudocolor dot plots illustrate the gating/sorting strategy for cluster-specific analysis, fluorescence-activated cell sorting and drug screening of tFL1 (A) and DLBCL1 (B) samples.

### Supplementary Figure 9

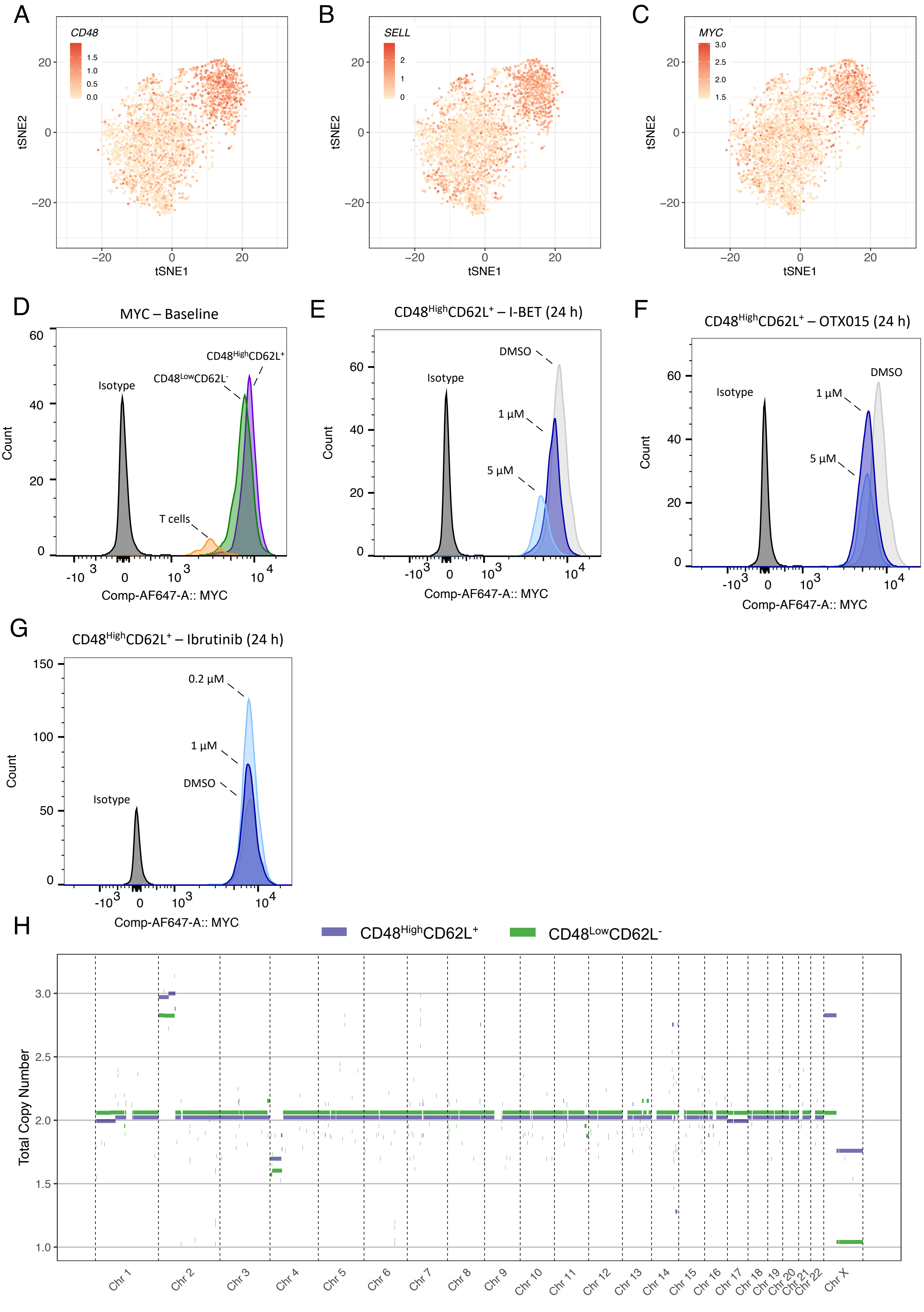

**Supplementary Figure 9.**

A-C) Single cell RNA expression profiles of B cells only (DLBCL1) were subjected to t-SNE and colored by *CD48* (A), *SELL* (B) and *MYC* (C) expression. D) DLBCL1 derived lymph node cells were stained for viability, CD19, CD3, CD48, CD62L and MYC or respective isotype control. Histograms show fluorescence intensity of MYC for isotype control, T cells, $CD48^{High}CD62L^{+}$ and $CD48^{Low}CD62L^{-}$ subclone. E-G) DLBCL1 derived lymph node cells were incubated with DMSO, I-BET-762 (E) at two concentrations (1 µM, 5µM), OTX015 (F) at two concentrations (1 µM, 5 µM), or ibrutinib (G) at two concentrations (0.2 µM, 1µM). After 24 hours, cells were harvested and stained as described in panel D. Histograms show fluorescence intensity for $CD48^{High}CD62L^{+}$ subclone. H) Line plot shows total cluster-specific copy number estimation of DLBCL1 sample for all chromosomes as indicated using whole genome sequencing. DMSO: Dimethyl sulfoxide.

### Supplementary Figure 10

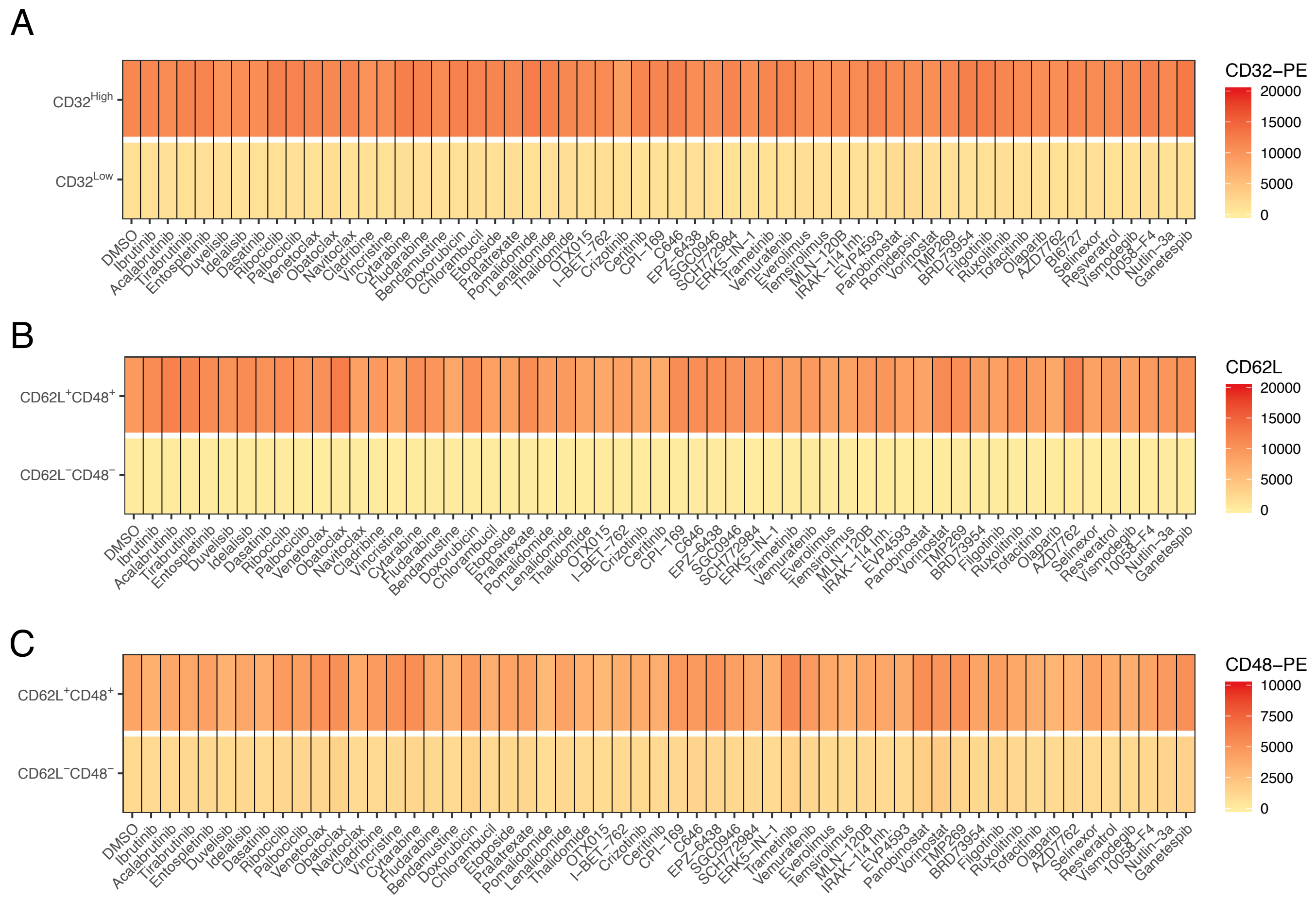


**Supplementary Figure 10.**

A-C) The heatmaps show the average median fluorescence intensities (MFI) of CD32-PE of tFL1 sample (A), CD62L-PE/Cy7 staining of sample DLBCL1 (B) and CD48-PE staining of sample DLBCL1 (C) of the investigated subpopulations upon drug treatment, as indicated.

### Supplementary Table Legends

**Supplementary Table 1.**

Patient characteristics showing sample name, diagnosis, subtype and clinical situation for all samples passed to scRNA-seq or flow cytometry analysis.

**Supplementary Table 2.**

Differentially expressed genes of T follicular helper cell population versus conventional T helper cells are shown. Genes were identified using the Wilcoxon rank-sum test. Shown are the mean expression levels for all cells of a specific cell population (pct.1) and all remaining cells (pct.2), adjusted p values (Wilcoxon test, Bonferroni correction) and average log fold changes. Only genes with an average log fold change greater than 0.5 and an adjusted p value lower than 0.05 are shown.

**Supplementary Table 3.**

Differentially expressed genes between both non-malignant B cells clusters. Genes were identified using the Wilcoxon rank-sum test. Shown are the differentially expressed genes, mean expression levels for all cells of a specific cell population (pct.1) and all remaining cells (pct.2), adjusted p values (Wilcoxon test, Bonferroni correction) and average log fold changes. Only genes with an average log fold change greater than 0.5 and an adjusted p value lower than 0.05 are shown.

**Supplementary Table 4.**

List of investigated ligand-receptor interactions. Given are gene names of protein A and B and the source.

**Supplementary Table 5.**

All drugs and concentrations (C1, C2, C3, C4, C5) used for subclone-specific drug screening. All numbers are given in nanomole per liter.

**Supplementary Table 6.**

All SNV that were detected in at least one of the sorted subclones of the tFL sample (healthy B cells, $CD32^{High}$ malignant cells, $CD32^{Low}$ malignant cells, whole tumor cells) using whole exome sequencing. Given are the chromosome (CHROM), position, coverage (COV), the allele count (AC) for each population, as well as the differential AC between both malignant populations. The two rightmost columns (private $CD32^{High}$, private $CD32^{Low}$) indicate whether the corresponding SNV was regarded as exclusive to one of the two malignant subclones. SNV: Single nucleotide variant.

**Supplementary Table 7.**

Exonic SNV that were detected in at least one of the sorted subclones of the DLBCL1 sample ($CD48^{High}$ malignant cells, $CD48^{Low}$ malignant cells, whole tumor cells) using whole genome sequencing. Given are the chromosome (CHROM), position, coverage (COV), the allele count (AC) for each population, as well as the differential AC between both malignant populations. The two rightmost columns (private $CD48^{High}$, private $CD48^{Low}$) indicate whether the corresponding SNV was regarded as exclusive for one of the two malignant subclones. SNV: Single nucleotide variant.

**Supplementary Table 8.**

Sequencing parameters determined using the Cell Ranger software (v2.1, 10x Genomics).
