## Supplementary File 1 for "Dissecting intratumor heterogeneity of nodal B cell lymphomas on the transcriptional, genetic, and drug response level"

scRNA-seq data analysis of lymph node derived B cell lymphoma samples


### scRNA-seq data analysis of lymph node derived B cell lymphoma samples

##### Differentially expressed genes between B cell subpopulations

###### 28 November 2019

### Contents

- 1 Load packages
- 2 Helper functions
  - 2.1 Extract data from Seurat objects
  - 2.2 Plot tSNE
- 3 Read data
- 4 DLBCL1
- 5 DLBCL2
- 6 DLBCL3
- 7 tFL1
- 8 tFL2
- 9 FL1
- 10 FL2
- 11 FL3
- 12 FL4
- 13 Merged B cells
- 14 Session Info

**LAST UPDATE AT**

```
   [1] "Thu Nov 28 22:01:06 2019"
```

This html file belongs to the **Manuscript**:

*“Dissecting intratumor heterogeneity of nodal B cell lymphoma on the transcriptional, genetic, and drug response level”* Roider *et al.*  
doi: …

It summarizes differentially expressed genes between different B cell subpopulations in malignant lymphoma samples. It enables the readers to browse differentially expressed genes for each B cell subpopulation interactively. Only genes with a minimal log fold change of 0.25 and an adjusted p value < 0.05 are shown.

**Sample abbreviations:**  
DLBCL: Diffuse large B cell lymphoma  
tFL: Transformed follicular lymphoma  
FL: Follicular lymphopma  
rLN: Reactive lymph node (non-malignant controls)

### 1 Load packages

```
library("Seurat")
library(dplyr)
library(DT)

filter <- dplyr::filter
```

### 2 Helper functions

#### 2.1 Extract data from Seurat objects

`get.data` extracts most important meta data and feature data, and puts them into a data frame.

```
get.data <- function(Sobj, genes) {
  
  # Extract Meta data
  df.meta <- data.frame(Cluster = Sobj@ident, Sobj@meta.data)

  # Extract tSNE coordinates if available
  if(!is.null(Sobj@dr$tsne))
    { 
    df.meta <- cbind(df.meta, Sobj@dr$tsne@cell.embeddings) 
    }
  
  # Extract umap coordinates if available
  if(!is.null(Sobj@dr$umap))
    { 
    df.meta <- cbind(df.meta, Sobj@dr$umap@cell.embeddings) 
    }

  # Extract gene expression data for expressed genes
  genes.red <- rownames(Sobj@data)[rownames(Sobj@data) %in% genes]
  dftotal <-  cbind(df.meta, FetchData(Sobj, genes.red))
  
  return(dftotal)

}
```

#### 2.2 Plot tSNE

`plot.TSNE` creates tSNE plots from Seurat objects in ggplot style.

```
plot.TSNE <- function(Sobj) {
  
get.data(Sobj, "") %>% 
  ggplot(aes(x=tSNE_1, y=tSNE_2, color=Cluster))+geom_point(size=1.25)+
  guides(color = guide_legend(override.aes = list(size = 3)))+
  theme_bw()

}
```

### 3 Read data

### 4 DLBCL1

```
plot.TSNE(sobj.B$DLBCL1)
```

```
Markers$DLBCL1 <- FindAllMarkers(sobj.B$DLBCL1, logfc.threshold = 0.25, only.pos = T)
Markers$DLBCL1[, 1:5] <- apply(Markers$DLBCL1[,1:5], 2, function(x) signif(x, digits = 3))

datatable(Markers$DLBCL1 %>% filter(p_val_adj < 0.05) %>% 
            select(7,6, 1:5), filter = "top", rownames=F)
```

### 5 DLBCL2

```
plot.TSNE(sobj.B$DLBCL2)
```

```
Markers$DLBCL2 <- FindAllMarkers(sobj.B$DLBCL2, logfc.threshold = 0.25, only.pos = T)
Markers$DLBCL2[, 1:5] <- apply(Markers$DLBCL2[,1:5], 2, function(x) signif(x, digits = 3))

datatable(Markers$DLBCL2 %>% filter(p_val_adj < 0.05) %>% 
            select(7,6, 1:5), filter = "top", rownames=F)
```

### 6 DLBCL3

```
plot.TSNE(sobj.B$DLBCL3)+ylim(-22,22)+xlim(-22,22)
```

```
Markers$DLBCL3 <- FindAllMarkers(sobj.B$DLBCL3, logfc.threshold = 0.25, only.pos = T)
Markers$DLBCL3[, 1:5] <- apply(Markers$DLBCL3[,1:5], 2, function(x) signif(x, digits = 3))

datatable(Markers$DLBCL3 %>% filter(p_val_adj < 0.05) %>% 
            select(7,6, 1:5), filter = "top", rownames=F)
```

### 7 tFL1

```
plot.TSNE(sobj.B$tFL1)+ylim(-25,25)+xlim(-25,25)
```

```
Markers$tFL1 <- FindAllMarkers(sobj.B$tFL1, logfc.threshold = 0.25, only.pos = T)
Markers$tFL1[, 1:5] <- apply(Markers$tFL1[,1:5], 2, function(x) signif(x, digits = 3))

datatable(Markers$tFL1 %>% filter(p_val_adj < 0.05) %>% 
            select(7,6, 1:5), filter = "top", rownames=F)
```

### 8 tFL2

```
plot.TSNE(sobj.B$tFL2)+ylim(-20,20)+xlim(-20,20)
```

```
Markers$tFL2 <- FindAllMarkers(sobj.B$tFL2, logfc.threshold = 0.25, only.pos = T)
Markers$tFL2[, 1:5] <- apply(Markers$tFL2[,1:5], 2, function(x) signif(x, digits = 3))

datatable(Markers$tFL2 %>% filter(p_val_adj < 0.05) %>% 
            select(7,6, 1:5), filter = "top", rownames=F)
```

# 9 FL1

```
plot.TSNE(sobj.B$FL1)+ylim(-30,30)+xlim(-30,30)
```

```
Markers$FL1 <- FindAllMarkers(sobj.B$FL1, logfc.threshold = 0.25, only.pos = T)
Markers$FL1[, 1:5] <- apply(Markers$FL1[,1:5], 2, function(x) signif(x, digits = 3))

datatable(Markers$FL1 %>% filter(p_val_adj < 0.05) %>% 
            select(7,6, 1:5), filter = "top", rownames=F)
```

# 10 FL2

```
plot.TSNE(sobj.B$FL2)+ylim(-30,30)+xlim(-30,30)
```

```
Markers$FL2 <- FindAllMarkers(sobj.B$FL2, logfc.threshold = 0.25, only.pos = T)
Markers$FL2[, 1:5] <- apply(Markers$FL2[,1:5], 2, function(x) signif(x, digits = 3))

datatable(Markers$FL2 %>% filter(p_val_adj < 0.05) %>% 
            select(7,6, 1:5), filter = "top", rownames=F)
```

# 11 FL3

```
plot.TSNE(sobj.B$FL3)+ylim(-30,30)+xlim(-30,30)+xlab("tSNE1")+ylab("tSNE2")+theme(axis.text = element_blank(), legend.position = "none", axis.ticks = element_blank())
```

```
Markers$FL3 <- FindAllMarkers(sobj.B$FL3, logfc.threshold = 0.25, only.pos = T)
Markers$FL3[, 1:5] <- apply(Markers$FL3[,1:5], 2, function(x) signif(x, digits = 3))

datatable(Markers$FL3 %>% filter(p_val_adj < 0.05) %>% 
            select(7,6, 1:5), filter = "top", rownames=F)
```

# 12 FL4

```
plot.TSNE(sobj.B$FL4)+ylim(-30,30)+xlim(-30,30)
```

```
Markers$FL4 <- FindAllMarkers(sobj.B$FL4, logfc.threshold = 0.25, only.pos = T)
Markers$FL4[, 1:5] <- apply(Markers$FL4[,1:5], 2, function(x) signif(x, digits = 3))

datatable(Markers$FL4 %>% filter(p_val_adj < 0.05) %>% 
            select(7,6, 1:5), filter = "top", rownames=F)
```

### 13 Merged B cells

```
get.data(TotalMerge_B, "") %>% 
  ggplot(aes(x=UMAP1, y=UMAP2, color=Cluster))+geom_point(size=0.75)+
  guides(color = guide_legend(override.aes = list(size = 3)))+
  theme_bw()
```

```
Markers$MergeB <- FindAllMarkers(TotalMerge_B, logfc.threshold = 0.25, only.pos = T)
```

```
Markers$MergeB[, 1:5] <- apply(Markers$MergeB[, 1:5], 2, function(x) {signif(x, digits = 3)})

datatable(Markers$MergeB %>% filter(p_val_adj < 0.05) %>% 
            select(7, 6, 1:5), filter = "top", rownames=F)
```

### 14 Session Info

```
sessionInfo()
```

```
   R version 3.6.1 (2019-07-05)
   Platform: x86_64-apple-darwin15.6.0 (64-bit)
   Running under: macOS Catalina 10.15.1
   
   Matrix products: default
   BLAS:   /Library/Frameworks/R.framework/Versions/3.6/Resources/lib/libRblas.0.dylib
   LAPACK: /Library/Frameworks/R.framework/Versions/3.6/Resources/lib/libRlapack.dylib
   
   locale:
   [1] en_US.UTF-8/en_US.UTF-8/en_US.UTF-8/C/en_US.UTF-8/en_US.UTF-8
   
   attached base packages:
   [1] stats     graphics  grDevices utils     datasets  methods   base     
   
   other attached packages:
   [1] DT_0.9           dplyr_0.8.3      Seurat_2.3.4     Matrix_1.2-17   
   [5] cowplot_1.0.0    ggplot2_3.2.1    knitr_1.25       BiocStyle_2.12.0
   
   loaded via a namespace (and not attached):
     [1] Rtsne_0.15          colorspace_1.4-1    class_7.3-15       
     [4] modeltools_0.2-22   ggridges_0.5.1      mclust_5.4.5       
     [7] htmlTable_1.13.2    base64enc_0.1-3     rstudioapi_0.10    
    [10] proxy_0.4-23        npsurv_0.4-0        flexmix_2.3-15     
    [13] bit64_0.9-7         codetools_0.2-16    splines_3.6.1      
    [16] R.methodsS3_1.7.1   lsei_1.2-0          robustbase_0.93-5  
    [19] zeallot_0.1.0       jsonlite_1.6        Formula_1.2-3      
    [22] ica_1.0-2           cluster_2.1.0       kernlab_0.9-27     
    [25] png_0.1-7           R.oo_1.23.0         shiny_1.4.0        
    [28] BiocManager_1.30.9  compiler_3.6.1      httr_1.4.1         
    [31] backports_1.1.5     fastmap_1.0.1       assertthat_0.2.1   
    [34] lazyeval_0.2.2      later_1.0.0         lars_1.2           
    [37] acepack_1.4.1       htmltools_0.4.0     tools_3.6.1        
    [40] igraph_1.2.4.1      gtable_0.3.0        glue_1.3.1         
    [43] RANN_2.6.1          reshape2_1.4.3      Rcpp_1.0.2         
    [46] vctrs_0.2.0         gdata_2.18.0        ape_5.3            
    [49] nlme_3.1-141        crosstalk_1.0.0     iterators_1.0.12   
    [52] fpc_2.2-3           gbRd_0.4-11         lmtest_0.9-37      
    [55] xfun_0.10           stringr_1.4.0       mime_0.7           
    [58] lifecycle_0.1.0     irlba_2.3.3         gtools_3.8.1       
    [61] DEoptimR_1.0-8      MASS_7.3-51.4       zoo_1.8-6          
    [64] scales_1.0.0        promises_1.1.0      doSNOW_1.0.18      
    [67] parallel_3.6.1      RColorBrewer_1.1-2  yaml_2.2.0         
    [70] reticulate_1.13     pbapply_1.4-2       gridExtra_2.3      
    [73] rpart_4.1-15        segmented_1.0-0     latticeExtra_0.6-28
    [76] stringi_1.4.3       foreach_1.4.7       checkmate_1.9.4    
    [79] caTools_1.17.1.2    bibtex_0.4.2        Rdpack_0.11-0      
    [82] SDMTools_1.1-221.1  rlang_0.4.1         pkgconfig_2.0.3    
    [85] dtw_1.21-3          prabclus_2.3-1      bitops_1.0-6       
    [88] evaluate_0.14       lattice_0.20-38     ROCR_1.0-7         
    [91] purrr_0.3.3         labeling_0.3        htmlwidgets_1.5.1  
    [94] bit_1.1-14          tidyselect_0.2.5    plyr_1.8.4         
    [97] magrittr_1.5        bookdown_0.14       R6_2.4.0           
   [100] snow_0.4-3          gplots_3.0.1.1      Hmisc_4.2-0        
   [103] pillar_1.4.2        foreign_0.8-72      withr_2.1.2        
   [106] fitdistrplus_1.0-14 mixtools_1.1.0      survival_2.44-1.1  
   [109] nnet_7.3-12         tsne_0.1-3          tibble_2.1.3       
   [112] crayon_1.3.4        hdf5r_1.3.0         KernSmooth_2.23-16 
   [115] rmarkdown_1.16      grid_3.6.1          data.table_1.12.6  
   [118] metap_1.1           digest_0.6.22       diptest_0.75-7     
   [121] xtable_1.8-4        httpuv_1.5.2        tidyr_1.0.0        
   [124] R.utils_2.9.0       stats4_3.6.1        munsell_0.5.0
```
