## Supplementary Table 1 for "Dissecting intratumor heterogeneity of nodal B cell lymphomas on the transcriptional, genetic, and drug response level"

| Running Number | Sample Name | Histological Diagnosis | Subtype/ Subclassification | Clinical Situation | scRNA-Seq |
|---|---|---|---|---|---|
| LN01 | rLN1 | Reactive Lymphdenitis | *Not applicable* | *Not applicable* | Yes |
| LN02 | rLN2 | Reactive Lymphdenitis | *Not applicable* | *Not applicable* | Yes |
| LN03 | rLN3 | Reactive Lymphdenitis | *Not applicable* | *Not applicable* | Yes |
| LN04 | LN4 | Reactive Lymphdenitis | *Not applicable* | *Not applicable* | No |
| LN05 | LN5 | Reactive Lymphdenitis | *Not applicable* | *Not applicable* | No |
| LN06 | LN6 | Reactive Lymphdenitis | *Not applicable* | *Not applicable* | No |
| LN07 | LN7 | Reactive Lymphdenitis | *Not applicable* | *Not applicable* | No |
| LN08 | LN8 | Reactive Lymphdenitis | *Not applicable* | *Not applicable* | No |
| LN09 | LN9 | Reactive Lymphdenitis | *Not applicable* | *Not applicable* | No |
| LN10 | LN10 | Mantle cell lymphoma | *Not applicable* | Initial diagnosis | No |
| LN11 | LN11 | Mantle cell lymphoma | *Not applicable* | Initial diagnosis | No |
| LN12 | LN12 | Mantle cell lymphoma | *Not applicable* | Relapse | No |
| LN13 | LN13 | Mantle cell lymphoma | *Not applicable* | Initial diagnosis | No |
| LN14 | FL1 | Follicular lymphoma | Grade 3a | Relapse | Yes |
| LN15 | FL2 | Follicular lymphoma | Grade 2 | Initial diagnosis | Yes |
| LN16 | FL3 | Follicular lymphoma | Grade 2 | Relapse | Yes |
| LN17 | FL4 | Follicular lymphoma | Grade 1 | Initial diagnosis | Yes |
| LN18 | LN18 | Follicular lymphoma | Grade 1 | Initial diagnosis | No |
| LN19 | LN19 | Follicular lymphoma | Grade 3a | Initial diagnosis | No |
| LN20 | LN20 | Follicular lymphoma | Grade 2 | Relapse | No |
| LN21 | LN21 | Follicular lymphoma | Grade 2 | Relapse | No |
| LN22 | LN22 | Follicular lymphoma | Grade 1 | Initial diagnosis | No |
| LN23 | LN23 | Follicular lymphoma | Grade 2 | Initial diagnosis | No |
| LN24 | LN24 | Follicular lymphoma | Grade 3a | Initial diagnosis | No |
| LN25 | LN25 | Follicular lymphoma | Grade 2 | Initial diagnosis | No |
| LN26 | tFL1 | Diffuse large B cell lymphoma | Germinal center B cell | Initial diagnosis | Yes |
| LN27 | tFL2 | Diffuse large B cell lymphoma | Germinal center B cell | Relapse | Yes |
| LN28 | DLBCL1 | Diffuse large B cell lymphoma | Germinal center B cell | Relapse | Yes |
| LN29 | DLBCL2 | Diffuse large B cell lymphoma | Germinal center B cell | Relapse | Yes |
| LN30 | DLBCL3 | Diffuse large B cell lymphoma | Non-germinal center B cell | Relapse | Yes |
| LN31 | LN31 | Diffuse large B cell lymphoma | Non-germinal center B cell | Initial diagnosis | No |
| LN32 | LN32 | Diffuse large B cell lymphoma | Germinal center B cell | Relapse | No |
| LN33 | LN33 | Diffuse large B cell lymphoma | Non-germinal center B cell | Initial diagnosis | No |
| LN34 | LN34 | Diffuse large B cell lymphoma | Non-germinal center B cell | Relapse | No |
| LN35 | LN35 | Chronic lymphocytic leukemia | *Not applicable* | Relapse | No |
| LN36 | LN36 | Chronic lymphocytic leukemia | *Not applicable* | Initial diagnosis | No |
| LN37 | LN37 | Chronic lymphocytic leukemia | *Not applicable* | Initial diagnosis | No |
| LN38 | LN38 | Chronic lymphocytic leukemia | *Not applicable* | Initial diagnosis | No |
| LN39 | LN39 | Chronic lymphocytic leukemia | *Not applicable* | Initial diagnosis | No |
| LN40 | LN40 | Chronic lymphocytic leukemia | *Not applicable* | Initial diagnosis | No |
| LN41 | LN41 | Chronic lymphocytic leukemia | *Not applicable* | Initial diagnosis | No |
