## Supplementary Table 2 for "Dissecting intratumor heterogeneity of nodal B cell lymphomas on the transcriptional, genetic, and drug response level"

| Gene | p_val | avg_logFC | pct.1 | pct.2 | p_val_adj |
|---|---|---|---|---|---|
| AAK1 | 4.84E-32 | -0.40 | 0.16 | 0.29 | 1.26E-27 |
| ABI3 | 6.45E-47 | -0.39 | 0.04 | 0.19 | 1.68E-42 |
| AC092580.4 | 1.08E-12 | -0.31 | 0.11 | 0.19 | 2.79E-08 |
| ACAP1 | 4.36E-26 | -0.38 | 0.52 | 0.59 | 1.13E-21 |
| ACTN1 | 7.68E-110 | 0.34 | 0.26 | 0.03 | 1.99E-105 |
| ACTR3 | 1.40E-26 | 0.21 | 0.76 | 0.54 | 3.64E-22 |
| ADD3 | 1.95E-84 | 0.53 | 0.45 | 0.18 | 5.07E-80 |
| ADGRE5 | 8.09E-22 | -0.46 | 0.23 | 0.33 | 2.10E-17 |
| ADI1 | 1.95E-71 | 0.25 | 0.34 | 0.11 | 5.05E-67 |
| AHI1 | 1.35E-47 | 0.22 | 0.24 | 0.08 | 3.52E-43 |
| ALDOA | 9.78E-35 | 0.26 | 0.82 | 0.61 | 2.54E-30 |
| ANP32B | 5.61E-43 | 0.21 | 0.62 | 0.36 | 1.46E-38 |
| ANXA1 | 1.23E-57 | -0.78 | 0.05 | 0.22 | 3.19E-53 |
| ANXA2 | 1.58E-17 | -0.37 | 0.17 | 0.26 | 4.11E-13 |
| ANXA2R | 4.71E-10 | -0.23 | 0.12 | 0.18 | 1.22E-05 |
| ANXA7 | 1.11E-43 | 0.23 | 0.55 | 0.30 | 2.88E-39 |
| AOAH | 1.80E-30 | -0.23 | 0.02 | 0.11 | 4.66E-26 |
| APMAP | 1.59E-15 | -0.33 | 0.23 | 0.31 | 4.14E-11 |
| APOBEC3G | 5.43E-51 | -0.60 | 0.19 | 0.37 | 1.41E-46 |
| ARL4C | 7.07E-76 | -0.64 | 0.15 | 0.38 | 1.84E-71 |
| ARL6IP1 | 1.43E-17 | 0.25 | 0.61 | 0.44 | 3.71E-13 |
| ARRB2 | 4.72E-16 | -0.21 | 0.07 | 0.14 | 1.23E-11 |
| ASCL2 | 5.69E-94 | 0.37 | 0.23 | 0.03 | 1.48E-89 |
| ATP5A1 | 1.50E-31 | 0.20 | 0.50 | 0.29 | 3.89E-27 |
| ATP5F1E | 6.78E-07 | -0.37 | 0.21 | 0.27 | 1.76E-02 |
| B2M | 1.44E-226 | -0.47 | 1.00 | 1.00 | 3.74E-222 |
| BATF | 2.43E-22 | 0.21 | 0.44 | 0.27 | 6.32E-18 |
| BCAT1 | 4.76E-68 | 0.22 | 0.19 | 0.03 | 1.24E-63 |
| BIN1 | 4.10E-07 | -0.28 | 0.32 | 0.34 | 1.07E-02 |
| BRK1 | 1.88E-39 | 0.21 | 0.72 | 0.46 | 4.87E-35 |
| BTG1 | 2.45E-86 | -0.59 | 0.96 | 0.97 | 6.37E-82 |
| BTG2 | 5.55E-11 | -0.41 | 0.45 | 0.47 | 1.44E-06 |
| BTLA | 9.76E-139 | 0.46 | 0.34 | 0.05 | 2.54E-134 |
| BTN3A1 | 4.46E-09 | -0.21 | 0.10 | 0.15 | 1.16E-04 |
| BTN3A2 | 6.17E-08 | -0.27 | 0.26 | 0.30 | 1.60E-03 |
| C10orf54 | 9.85E-34 | 0.24 | 0.25 | 0.10 | 2.56E-29 |
| C12orf75 | 2.08E-48 | -0.44 | 0.05 | 0.21 | 5.41E-44 |
| C16orf87 | 1.73E-61 | 0.28 | 0.38 | 0.15 | 4.48E-57 |
| C1orf228 | 5.98E-101 | 0.41 | 0.31 | 0.07 | 1.55E-96 |
| C5orf56 | 4.65E-12 | -0.25 | 0.10 | 0.17 | 1.21E-07 |
| C9orf16 | 8.90E-96 | 0.49 | 0.80 | 0.48 | 2.31E-91 |
| CADM1 | 1.01E-07 | -0.29 | 0.16 | 0.21 | 2.63E-03 |
| CALM1 | 2.37E-16 | -0.31 | 0.76 | 0.73 | 6.15E-12 |
| CAPN2 | 3.63E-43 | -0.36 | 0.06 | 0.20 | 9.44E-39 |
| CARD16 | 1.09E-09 | -0.28 | 0.22 | 0.28 | 2.83E-05 |
| CASP1 | 1.32E-12 | -0.23 | 0.11 | 0.18 | 3.43E-08 |
| CAV1 | 2.88E-84 | 0.99 | 0.40 | 0.15 | 7.48E-80 |
| CCDC50 | 2.95E-57 | 0.22 | 0.19 | 0.04 | 7.67E-53 |
| CCL3 | 5.81E-49 | -1.70 | 0.05 | 0.20 | 1.51E-44 |
| CCL3L3 | 6.40E-50 | -1.10 | 0.02 | 0.16 | 1.66E-45 |

| | | | | | |
|---|---|---|---|---|---|
| CCL4 | 5.06E-209 | -2.29 | 0.09 | 0.53 | 1.31E-204 |
| CCL4L2 | 1.51E-85 | -1.55 | 0.03 | 0.26 | 3.93E-81 |
| CCL5 | 0.00E+00 | -3.40 | 0.17 | 0.81 | 0.00E+00 |
| CCNI | 1.12E-45 | 0.28 | 0.85 | 0.61 | 2.90E-41 |
| CCR5 | 3.08E-21 | -0.23 | 0.03 | 0.11 | 7.99E-17 |
| CD200 | 1.39E-99 | 0.55 | 0.33 | 0.08 | 3.61E-95 |
| CD27 | 1.36E-21 | -0.38 | 0.81 | 0.79 | 3.54E-17 |
| CD38 | 3.89E-10 | -0.21 | 0.09 | 0.15 | 1.01E-05 |
| CD3E | 9.95E-27 | -0.36 | 0.80 | 0.78 | 2.58E-22 |
| CD4 | 4.73E-73 | 0.26 | 0.32 | 0.09 | 1.23E-68 |
| CD40LG | 4.80E-224 | 0.83 | 0.37 | 0.02 | 1.25E-219 |
| CD44 | 8.91E-19 | -0.38 | 0.58 | 0.60 | 2.31E-14 |
| CD48 | 1.43E-11 | -0.28 | 0.49 | 0.50 | 3.71E-07 |
| CD59 | 1.93E-112 | 0.46 | 0.41 | 0.11 | 5.02E-108 |
| CD69 | 1.31E-13 | -0.31 | 0.90 | 0.86 | 3.39E-09 |
| CD74 | 2.02E-99 | -0.55 | 0.93 | 0.95 | 5.25E-95 |
| CD79A | 1.59E-29 | 0.41 | 0.27 | 0.13 | 4.12E-25 |
| CD82 | 6.25E-41 | 0.21 | 0.43 | 0.22 | 1.62E-36 |
| CD8A | 3.57E-116 | -0.99 | 0.07 | 0.36 | 9.27E-112 |
| CD8B | 2.01E-22 | -0.38 | 0.21 | 0.33 | 5.21E-18 |
| CD96 | 3.43E-14 | -0.32 | 0.29 | 0.36 | 8.92E-10 |
| CD99 | 3.94E-23 | -0.38 | 0.60 | 0.62 | 1.02E-18 |
| CHST12 | 3.13E-34 | -0.42 | 0.12 | 0.26 | 8.14E-30 |
| CLDND1 | 9.99E-48 | -0.62 | 0.16 | 0.33 | 2.59E-43 |
| CLEC2B | 1.15E-34 | -0.50 | 0.28 | 0.41 | 3.00E-30 |
| CLIC1 | 1.40E-20 | -0.37 | 0.73 | 0.69 | 3.63E-16 |
| CMC1 | 1.27E-108 | -1.56 | 0.22 | 0.49 | 3.30E-104 |
| CNIH1 | 6.55E-75 | 0.33 | 0.55 | 0.24 | 1.70E-70 |
| CNN2 | 9.02E-15 | -0.33 | 0.28 | 0.35 | 2.34E-10 |
| CORO1A | 3.55E-89 | -0.62 | 0.83 | 0.86 | 9.22E-85 |
| CORO1B | 2.25E-154 | 0.76 | 0.79 | 0.38 | 5.85E-150 |
| COX6C | 5.14E-25 | 0.20 | 0.88 | 0.71 | 1.33E-20 |
| CPM | 9.64E-108 | 0.35 | 0.27 | 0.04 | 2.50E-103 |
| CST7 | 0.00E+00 | -1.79 | 0.24 | 0.85 | 0.00E+00 |
| CTLA4 | 3.97E-58 | 0.26 | 0.30 | 0.10 | 1.03E-53 |
| CTSB | 3.82E-49 | 0.25 | 0.46 | 0.23 | 9.91E-45 |
| CTSC | 6.17E-22 | -0.39 | 0.27 | 0.37 | 1.60E-17 |
| CTSW | 2.81E-158 | -1.11 | 0.03 | 0.38 | 7.29E-154 |
| CTTN | 2.50E-67 | 0.24 | 0.20 | 0.04 | 6.49E-63 |
| CXCL13 | 3.59E-128 | 2.36 | 0.51 | 0.20 | 9.32E-124 |
| CXCR3 | 1.10E-39 | -0.48 | 0.17 | 0.32 | 2.86E-35 |
| CXCR4 | 8.68E-52 | -0.82 | 0.82 | 0.82 | 2.26E-47 |
| CXCR5 | 1.34E-41 | 0.21 | 0.22 | 0.07 | 3.48E-37 |
| CXCR6 | 3.88E-45 | -0.40 | 0.01 | 0.13 | 1.01E-40 |
| CYBA | 1.94E-138 | -0.69 | 0.88 | 0.92 | 5.05E-134 |
| CYTOR | 4.99E-15 | -0.35 | 0.07 | 0.14 | 1.29E-10 |
| DHRS7 | 6.08E-24 | 0.22 | 0.45 | 0.28 | 1.58E-19 |
| DNAJB1 | 1.71E-49 | -0.65 | 0.46 | 0.61 | 4.43E-45 |
| DOK2 | 1.08E-14 | -0.35 | 0.19 | 0.27 | 2.80E-10 |
| DRAIC | 1.76E-21 | 0.21 | 0.12 | 0.05 | 4.58E-17 |
| DTHD1 | 4.01E-07 | -0.25 | 0.15 | 0.20 | 1.04E-02 |
| DUSP2 | 1.88E-101 | -1.01 | 0.60 | 0.75 | 4.88E-97 |

| | | | | | |
|---|---|---|---|---|---|
| DUSP4 | 1.18E-06 | -0.42 | 0.23 | 0.26 | 3.06E-02 |
| DUSP6 | 1.70E-38 | 0.26 | 0.20 | 0.07 | 4.43E-34 |
| EEF1A1 | 9.73E-114 | 0.35 | 1.00 | 1.00 | 2.53E-109 |
| EEF2 | 6.52E-32 | 0.22 | 0.92 | 0.77 | 1.69E-27 |
| EIF3L | 8.91E-40 | 0.23 | 0.66 | 0.40 | 2.31E-35 |
| EMP3 | 3.60E-72 | -0.69 | 0.14 | 0.37 | 9.35E-68 |
| ENO1 | 1.35E-29 | 0.34 | 0.80 | 0.60 | 3.51E-25 |
| EOMES | 5.09E-43 | -0.41 | 0.11 | 0.28 | 1.32E-38 |
| ERH | 7.83E-34 | 0.21 | 0.72 | 0.47 | 2.03E-29 |
| ESD | 1.77E-51 | 0.24 | 0.41 | 0.18 | 4.61E-47 |
| EVL | 7.02E-71 | -0.53 | 0.72 | 0.82 | 1.82E-66 |
| F2R | 4.86E-13 | -0.26 | 0.13 | 0.21 | 1.26E-08 |
| FABP5 | 8.23E-35 | 0.71 | 0.60 | 0.43 | 2.14E-30 |
| FAIM2 | 2.42E-55 | 0.23 | 0.14 | 0.02 | 6.29E-51 |
| FAM107B | 2.93E-31 | 0.21 | 0.56 | 0.34 | 7.60E-27 |
| FAM43A | 7.83E-83 | 0.37 | 0.39 | 0.13 | 2.03E-78 |
| FBLN7 | 2.54E-78 | 0.27 | 0.21 | 0.03 | 6.59E-74 |
| FCRL3 | 8.55E-18 | -0.23 | 0.05 | 0.12 | 2.22E-13 |
| FKBP1A | 6.72E-43 | 0.29 | 0.71 | 0.46 | 1.75E-38 |
| FKBP5 | 2.85E-252 | 0.90 | 0.59 | 0.12 | 7.40E-248 |
| FLNA | 6.40E-15 | -0.21 | 0.06 | 0.12 | 1.66E-10 |
| FOSB | 7.77E-16 | -0.47 | 0.53 | 0.53 | 2.02E-11 |
| FOXN3 | 1.41E-07 | -0.22 | 0.14 | 0.18 | 3.67E-03 |
| FTL | 1.15E-24 | -0.35 | 0.93 | 0.89 | 2.98E-20 |
| FXYD2 | 3.11E-48 | -0.54 | 0.01 | 0.13 | 8.07E-44 |
| FXYD5 | 1.18E-33 | 0.32 | 0.69 | 0.48 | 3.06E-29 |
| FYB | 3.67E-13 | 0.27 | 0.54 | 0.40 | 9.53E-09 |
| GADD45B | 7.08E-39 | -0.68 | 0.30 | 0.44 | 1.84E-34 |
| GADD45G | 2.59E-65 | 0.32 | 0.38 | 0.14 | 6.73E-61 |
| GAPDH | 1.85E-84 | 0.41 | 0.98 | 0.88 | 4.80E-80 |
| GIMAP1 | 5.98E-34 | -0.39 | 0.12 | 0.26 | 1.55E-29 |
| GIMAP4 | 5.64E-38 | -0.53 | 0.44 | 0.55 | 1.46E-33 |
| GIMAP7 | 1.39E-64 | -0.71 | 0.43 | 0.60 | 3.61E-60 |
| GLIPR2 | 8.39E-22 | -0.21 | 0.04 | 0.12 | 2.18E-17 |
| GNG4 | 2.01E-175 | 0.70 | 0.39 | 0.05 | 5.21E-171 |
| GPR171 | 2.95E-31 | -0.33 | 0.06 | 0.17 | 7.67E-27 |
| GPX1 | 7.05E-38 | 0.31 | 0.37 | 0.18 | 1.83E-33 |
| GSTP1 | 8.95E-23 | -0.40 | 0.33 | 0.42 | 2.33E-18 |
| GUK1 | 2.36E-07 | -0.26 | 0.61 | 0.57 | 6.14E-03 |
| GZMA | 0.00E+00 | -2.64 | 0.08 | 0.75 | 0.00E+00 |
| GZMB | 3.53E-19 | -0.89 | 0.04 | 0.11 | 9.18E-15 |
| GZMH | 9.63E-65 | -0.98 | 0.02 | 0.20 | 2.50E-60 |
| GZMK | 0.00E+00 | -2.35 | 0.35 | 0.97 | 0.00E+00 |
| GZMM | 3.65E-08 | -0.40 | 0.42 | 0.43 | 9.49E-04 |
| H2AFZ | 3.82E-129 | 0.95 | 0.85 | 0.55 | 9.91E-125 |
| H3F3A | 1.48E-53 | 0.32 | 0.96 | 0.81 | 3.85E-49 |
| HAVCR2 | 1.72E-49 | -0.49 | 0.03 | 0.18 | 4.46E-45 |
| HCST | 1.00E-143 | -0.84 | 0.69 | 0.83 | 2.60E-139 |
| HIF1A | 3.63E-70 | 0.34 | 0.55 | 0.26 | 9.44E-66 |
| HLA-C | 5.20E-42 | -0.26 | 1.00 | 1.00 | 1.35E-37 |
| HLA-DPA1 | 2.51E-34 | -0.41 | 0.58 | 0.65 | 6.51E-30 |
| HLA-DPB1 | 1.65E-82 | -0.69 | 0.51 | 0.69 | 4.29E-78 |

| | | | | | |
|---|---|---|---|---|---|
| HLA-DQA1 | 2.35E-12 | -0.31 | 0.17 | 0.24 | 6.09E-08 |
| HLA-DQB1 | 6.93E-22 | -0.34 | 0.29 | 0.40 | 1.80E-17 |
| HLA-DRB1 | 9.02E-49 | -0.45 | 0.49 | 0.65 | 2.34E-44 |
| HLA-DRB5 | 2.99E-10 | -0.30 | 0.28 | 0.35 | 7.77E-06 |
| HLA-F | 1.68E-11 | -0.34 | 0.54 | 0.54 | 4.36E-07 |
| HMGN1 | 6.65E-44 | 0.26 | 0.90 | 0.67 | 1.73E-39 |
| HNRNPA1 | 3.02E-36 | 0.24 | 0.97 | 0.91 | 7.85E-32 |
| HSP90AB1 | 3.86E-18 | 0.22 | 0.86 | 0.72 | 1.00E-13 |
| ICA1 | 0.00E+00 | 1.23 | 0.75 | 0.18 | 0.00E+00 |
| ICOS | 9.88E-89 | 0.42 | 0.43 | 0.15 | 2.57E-84 |
| ID2 | 6.37E-10 | -0.59 | 0.48 | 0.49 | 1.65E-05 |
| ID3 | 4.20E-33 | 0.30 | 0.21 | 0.09 | 1.09E-28 |
| IER2 | 3.15E-81 | -1.00 | 0.72 | 0.80 | 8.17E-77 |
| IFI27L2 | 5.47E-08 | -0.29 | 0.24 | 0.28 | 1.42E-03 |
| IFI6 | 1.71E-15 | -0.36 | 0.14 | 0.22 | 4.45E-11 |
| IFNG | 2.01E-40 | -0.91 | 0.25 | 0.40 | 5.23E-36 |
| IGFBP4 | 3.40E-70 | 0.27 | 0.26 | 0.06 | 8.83E-66 |
| IGHA1 | 2.02E-41 | 0.23 | 0.31 | 0.13 | 5.24E-37 |
| IGHG3 | 2.34E-10 | -0.92 | 0.11 | 0.05 | 6.08E-06 |
| IGHM | 1.78E-42 | 0.21 | 0.43 | 0.22 | 4.63E-38 |
| IL10RA | 9.02E-50 | -0.47 | 0.11 | 0.29 | 2.34E-45 |
| IL32 | 2.23E-86 | -0.52 | 0.89 | 0.92 | 5.79E-82 |
| IL6R | 2.89E-119 | 0.41 | 0.34 | 0.07 | 7.51E-115 |
| IL6ST | 1.83E-106 | 0.50 | 0.36 | 0.09 | 4.76E-102 |
| IRF1 | 8.31E-15 | -0.37 | 0.38 | 0.44 | 2.16E-10 |
| ISG15 | 3.80E-07 | -0.49 | 0.32 | 0.34 | 9.86E-03 |
| ISG20 | 1.08E-16 | 0.21 | 0.74 | 0.56 | 2.79E-12 |
| ITGA4 | 4.54E-15 | -0.26 | 0.10 | 0.17 | 1.18E-10 |
| ITGB2 | 5.80E-24 | -0.37 | 0.60 | 0.63 | 1.51E-19 |
| ITGB7 | 1.32E-36 | -0.31 | 0.04 | 0.16 | 3.42E-32 |
| ITM2A | 4.58E-160 | 0.99 | 0.94 | 0.69 | 1.19E-155 |
| ITM2B | 3.48E-11 | -0.21 | 0.89 | 0.84 | 9.05E-07 |
| ITM2C | 4.78E-28 | -0.51 | 0.07 | 0.18 | 1.24E-23 |
| JAKMIP1 | 1.35E-34 | -0.27 | 0.03 | 0.14 | 3.51E-30 |
| JAML | 8.29E-44 | -0.48 | 0.07 | 0.22 | 2.15E-39 |
| JUN | 1.02E-36 | -0.71 | 0.94 | 0.94 | 2.65E-32 |
| JUNB | 1.28E-26 | -0.43 | 0.96 | 0.96 | 3.32E-22 |
| KLF2 | 5.41E-28 | -0.50 | 0.12 | 0.23 | 1.41E-23 |
| KLRB1 | 6.79E-34 | 0.31 | 0.20 | 0.07 | 1.76E-29 |
| KLRG1 | 5.44E-64 | -0.65 | 0.05 | 0.23 | 1.41E-59 |
| KSR2 | 3.55E-65 | 0.21 | 0.20 | 0.04 | 9.22E-61 |
| LAG3 | 2.34E-44 | -0.95 | 0.24 | 0.40 | 6.09E-40 |
| LAPTM5 | 5.64E-17 | -0.27 | 0.73 | 0.72 | 1.47E-12 |
| LAT | 2.01E-28 | 0.28 | 0.81 | 0.61 | 5.21E-24 |
| LBH | 2.89E-35 | 0.27 | 0.66 | 0.43 | 7.50E-31 |
| LDHA | 1.87E-17 | 0.23 | 0.74 | 0.56 | 4.85E-13 |
| LGALS1 | 6.70E-19 | -0.46 | 0.05 | 0.13 | 1.74E-14 |
| LGMN | 3.22E-41 | 0.20 | 0.12 | 0.02 | 8.37E-37 |
| LIMS1 | 1.71E-87 | 0.43 | 0.69 | 0.36 | 4.43E-83 |
| LINC00152 | 4.51E-21 | -0.49 | 0.31 | 0.39 | 1.17E-16 |
| LINC00861 | 8.44E-42 | -0.46 | 0.10 | 0.25 | 2.19E-37 |
| LINC01480 | 7.80E-96 | 0.45 | 0.23 | 0.03 | 2.03E-91 |

| | | | | | |
|---|---|---|---|---|---|
| LITAF | 3.73E-28 | -0.46 | 0.26 | 0.38 | 9.68E-24 |
| LSP1 | 8.82E-10 | -0.28 | 0.54 | 0.54 | 2.29E-05 |
| LY6E | 3.22E-34 | -0.49 | 0.34 | 0.47 | 8.36E-30 |
| LY96 | 1.32E-64 | 0.28 | 0.32 | 0.11 | 3.42E-60 |
| LYAR | 2.12E-56 | -0.65 | 0.12 | 0.31 | 5.50E-52 |
| LYST | 7.18E-81 | -0.74 | 0.22 | 0.45 | 1.87E-76 |
| MAF | 3.16E-103 | 0.41 | 0.56 | 0.21 | 8.21E-99 |
| MAGEH1 | 1.61E-120 | 0.51 | 0.44 | 0.12 | 4.17E-116 |
| MALAT1 | 1.14E-132 | -0.57 | 1.00 | 1.00 | 2.97E-128 |
| MGAT4A | 8.44E-22 | -0.30 | 0.07 | 0.16 | 2.19E-17 |
| MIAT | 7.61E-25 | -0.31 | 0.06 | 0.16 | 1.98E-20 |
| MIF | 6.70E-28 | 0.32 | 0.81 | 0.65 | 1.74E-23 |
| MINOS1 | 3.29E-49 | 0.24 | 0.66 | 0.38 | 8.55E-45 |
| MT-ND2 | 6.71E-10 | -0.26 | 0.96 | 0.94 | 1.74E-05 |
| MT1F | 1.55E-27 | -0.26 | 0.04 | 0.14 | 4.02E-23 |
| MT2A | 1.26E-34 | -0.68 | 0.31 | 0.45 | 3.26E-30 |
| MTRNR2L1 | 3.12E-10 | -0.40 | 0.14 | 0.20 | 8.11E-06 |
| MX1 | 3.98E-14 | -0.22 | 0.05 | 0.11 | 1.03E-09 |
| MYADM | 4.99E-35 | -0.40 | 0.04 | 0.15 | 1.30E-30 |
| MYL12A | 2.57E-81 | -0.61 | 0.88 | 0.87 | 6.67E-77 |
| MYO1F | 1.14E-66 | -0.51 | 0.06 | 0.26 | 2.97E-62 |
| MYO1G | 8.96E-25 | -0.26 | 0.07 | 0.18 | 2.33E-20 |
| NAP1L1 | 1.00E-53 | 0.30 | 0.84 | 0.59 | 2.61E-49 |
| NAP1L4 | 9.72E-27 | 0.22 | 0.49 | 0.31 | 2.53E-22 |
| NDFIP1 | 4.86E-56 | 0.31 | 0.63 | 0.36 | 1.26E-51 |
| NDUFB2 | 4.42E-51 | 0.38 | 0.74 | 0.47 | 1.15E-46 |
| NDUFV2 | 3.56E-48 | 0.27 | 0.57 | 0.31 | 9.24E-44 |
| NEAT1 | 1.35E-20 | -0.45 | 0.69 | 0.70 | 3.50E-16 |
| NFATC1 | 1.94E-92 | 0.40 | 0.38 | 0.11 | 5.03E-88 |
| NFIA | 5.95E-66 | 0.21 | 0.14 | 0.01 | 1.55E-61 |
| NFKBIA | 7.65E-13 | 0.22 | 0.67 | 0.51 | 1.99E-08 |
| NKG7 | 0.00E+00 | -2.59 | 0.09 | 0.69 | 0.00E+00 |
| NMB | 7.66E-102 | 0.74 | 0.38 | 0.11 | 1.99E-97 |
| NPM1 | 6.17E-28 | 0.25 | 0.93 | 0.79 | 1.60E-23 |
| NR3C1 | 3.94E-226 | 0.85 | 0.66 | 0.18 | 1.02E-221 |
| OASL | 1.65E-15 | -0.23 | 0.05 | 0.12 | 4.30E-11 |
| PAG1 | 1.30E-14 | -0.26 | 0.09 | 0.17 | 3.38E-10 |
| PARP1 | 1.55E-54 | 0.31 | 0.62 | 0.35 | 4.02E-50 |
| PARP8 | 2.48E-21 | -0.27 | 0.08 | 0.17 | 6.43E-17 |
| PASK | 2.41E-152 | 0.70 | 0.58 | 0.18 | 6.25E-148 |
| PCAT29 | 4.93E-79 | 0.42 | 0.45 | 0.17 | 1.28E-74 |
| PDCD1 | 5.48E-128 | 0.64 | 0.65 | 0.27 | 1.42E-123 |
| PDLIM2 | 1.49E-19 | -0.29 | 0.10 | 0.19 | 3.88E-15 |
| PEBP1 | 1.06E-70 | 0.36 | 0.73 | 0.42 | 2.76E-66 |
| PECAM1 | 2.55E-44 | -0.34 | 0.03 | 0.17 | 6.61E-40 |
| PGAM1 | 5.64E-55 | 0.39 | 0.67 | 0.40 | 1.47E-50 |
| PGK1 | 1.66E-17 | 0.21 | 0.68 | 0.49 | 4.31E-13 |
| PHACTR2 | 1.80E-50 | 0.25 | 0.41 | 0.18 | 4.67E-46 |
| PIK3R1 | 5.82E-15 | -0.32 | 0.11 | 0.18 | 1.51E-10 |
| PIP4K2A | 5.84E-37 | -0.52 | 0.25 | 0.39 | 1.52E-32 |
| PKM | 1.37E-115 | 0.70 | 0.83 | 0.52 | 3.55E-111 |
| PLEK | 1.38E-43 | -0.46 | 0.12 | 0.28 | 3.58E-39 |

| | | | | | |
|---|---|---|---|---|---|
| PLEKHF1 | 1.02E-08 | -0.23 | 0.11 | 0.16 | 2.65E-04 |
| POU2AF1 | 6.49E-77 | 0.34 | 0.31 | 0.09 | 1.69E-72 |
| PPA1 | 1.38E-41 | 0.22 | 0.43 | 0.22 | 3.59E-37 |
| PPDPF | 1.35E-13 | -0.30 | 0.80 | 0.76 | 3.50E-09 |
| PPP1CC | 4.46E-171 | 0.81 | 0.80 | 0.41 | 1.16E-166 |
| PPP2R5C | 2.13E-08 | -0.26 | 0.68 | 0.65 | 5.53E-04 |
| PRDM1 | 3.45E-28 | -0.34 | 0.06 | 0.16 | 8.97E-24 |
| PRDX1 | 6.04E-32 | 0.26 | 0.59 | 0.36 | 1.57E-27 |
| PRDX2 | 1.82E-56 | 0.32 | 0.66 | 0.38 | 4.74E-52 |
| PRF1 | 2.69E-85 | -0.85 | 0.10 | 0.34 | 6.99E-81 |
| PRNP | 2.42E-44 | 0.26 | 0.28 | 0.11 | 6.28E-40 |
| PSIP1 | 7.81E-65 | 0.30 | 0.51 | 0.23 | 2.03E-60 |
| PSMB9 | 5.39E-18 | -0.31 | 0.70 | 0.69 | 1.40E-13 |
| PSME1 | 1.36E-08 | -0.20 | 0.79 | 0.75 | 3.53E-04 |
| PTMA | 6.00E-26 | 0.25 | 1.00 | 1.00 | 1.56E-21 |
| PTPN13 | 2.70E-58 | 0.25 | 0.22 | 0.06 | 7.02E-54 |
| PTPN2 | 2.39E-55 | 0.26 | 0.50 | 0.24 | 6.22E-51 |
| PVALB | 2.29E-114 | 0.72 | 0.19 | 0.01 | 5.94E-110 |
| PYCARD | 1.81E-12 | -0.26 | 0.13 | 0.19 | 4.71E-08 |
| RAB11FIP1 | 4.03E-53 | 0.27 | 0.32 | 0.12 | 1.05E-48 |
| RABAC1 | 2.63E-08 | -0.28 | 0.32 | 0.36 | 6.82E-04 |
| RAN | 2.90E-34 | 0.26 | 0.74 | 0.49 | 7.54E-30 |
| RAP1A | 3.62E-54 | 0.31 | 0.67 | 0.39 | 9.41E-50 |
| RARRES3 | 8.64E-50 | -0.49 | 0.48 | 0.61 | 2.24E-45 |
| RASAL3 | 1.34E-11 | -0.27 | 0.23 | 0.30 | 3.48E-07 |
| RASGRP1 | 1.79E-11 | -0.28 | 0.16 | 0.22 | 4.64E-07 |
| RCSD1 | 9.36E-09 | -0.25 | 0.22 | 0.27 | 2.43E-04 |
| RFTN1 | 6.15E-08 | -0.21 | 0.10 | 0.15 | 1.60E-03 |
| RGL4 | 1.75E-21 | -0.26 | 0.07 | 0.16 | 4.55E-17 |
| RGS1 | 2.82E-131 | -1.28 | 0.58 | 0.79 | 7.32E-127 |
| RILPL2 | 3.29E-52 | 0.29 | 0.47 | 0.23 | 8.55E-48 |
| RNASET2 | 4.79E-62 | 0.35 | 0.62 | 0.33 | 1.24E-57 |
| RNF213 | 2.38E-35 | -0.49 | 0.42 | 0.52 | 6.17E-31 |
| RP11-132N15.3 | 5.04E-66 | 0.24 | 0.12 | 0.01 | 1.31E-61 |
| RP11-455F5.5 | 2.36E-81 | 0.42 | 0.26 | 0.06 | 6.12E-77 |
| RP5-1028K7.2 | 3.55E-49 | 0.38 | 0.52 | 0.29 | 9.22E-45 |
| RPL15 | 1.67E-55 | 0.22 | 1.00 | 1.00 | 4.33E-51 |
| RPL23A | 1.38E-42 | -0.29 | 1.00 | 1.00 | 3.59E-38 |
| RPL27A | 3.53E-22 | -0.23 | 1.00 | 0.99 | 9.18E-18 |
| RPL36A | 3.94E-08 | -0.20 | 0.93 | 0.89 | 1.02E-03 |
| RPL38 | 9.59E-12 | -0.21 | 0.97 | 0.93 | 2.49E-07 |
| RPL7 | 1.29E-50 | 0.23 | 1.00 | 0.99 | 3.36E-46 |
| RPLP1 | 2.53E-107 | 0.33 | 1.00 | 1.00 | 6.58E-103 |
| RPLP2 | 1.98E-23 | -0.26 | 1.00 | 1.00 | 5.14E-19 |
| RPS12 | 1.67E-44 | -0.33 | 1.00 | 0.99 | 4.33E-40 |
| RPS26 | 6.84E-07 | -0.23 | 0.94 | 0.88 | 1.78E-02 |
| RPS27 | 1.48E-59 | -0.28 | 1.00 | 1.00 | 3.85E-55 |
| RPS29 | 4.38E-62 | -0.38 | 1.00 | 1.00 | 1.14E-57 |
| RPS8 | 7.37E-79 | 0.32 | 1.00 | 0.98 | 1.91E-74 |
| RUNX3 | 1.15E-35 | -0.37 | 0.06 | 0.19 | 2.97E-31 |
| S100A10 | 4.91E-23 | -0.40 | 0.21 | 0.32 | 1.28E-18 |
| S100A11 | 4.12E-18 | 0.21 | 0.31 | 0.18 | 1.07E-13 |

| | | | | | |
|---|---|---|---|---|---|
| S100A6 | 2.14E-67 | -0.77 | 0.47 | 0.62 | 5.56E-63 |
| SAMD3 | 4.52E-70 | -0.42 | 0.01 | 0.18 | 1.17E-65 |
| SAMSN1 | 8.47E-10 | -0.26 | 0.15 | 0.22 | 2.20E-05 |
| SCGB3A1 | 2.63E-62 | 0.30 | 0.14 | 0.01 | 6.84E-58 |
| SEC11A | 8.94E-43 | 0.23 | 0.58 | 0.33 | 2.32E-38 |
| SELT | 5.57E-64 | 0.38 | 0.60 | 0.32 | 1.45E-59 |
| SEPT1 | 6.08E-07 | -0.27 | 0.53 | 0.51 | 1.58E-02 |
| SEPT6 | 7.95E-46 | 0.28 | 0.63 | 0.37 | 2.06E-41 |
| SERPINE2 | 3.74E-61 | 0.48 | 0.15 | 0.02 | 9.72E-57 |
| SFXN1 | 4.39E-58 | 0.31 | 0.43 | 0.19 | 1.14E-53 |
| SH2D1A | 3.82E-24 | 0.27 | 0.69 | 0.50 | 9.93E-20 |
| SH3BGRL3 | 5.41E-11 | -0.24 | 0.93 | 0.88 | 1.41E-06 |
| SH3TC1 | 1.08E-50 | 0.24 | 0.24 | 0.08 | 2.80E-46 |
| SIAH2 | 5.67E-45 | 0.26 | 0.46 | 0.23 | 1.47E-40 |
| SIT1 | 2.02E-09 | -0.31 | 0.36 | 0.39 | 5.25E-05 |
| SKP1 | 1.49E-38 | 0.24 | 0.91 | 0.72 | 3.88E-34 |
| SLAMF7 | 3.10E-25 | -0.30 | 0.05 | 0.15 | 8.04E-21 |
| SLC20A1 | 1.56E-14 | -0.26 | 0.11 | 0.18 | 4.06E-10 |
| SLC25A5 | 1.73E-40 | 0.25 | 0.70 | 0.44 | 4.48E-36 |
| SLC2A3 | 3.97E-08 | -0.32 | 0.30 | 0.34 | 1.03E-03 |
| SLC9A9 | 9.60E-48 | 0.26 | 0.24 | 0.08 | 2.49E-43 |
| SLF1 | 3.05E-25 | -0.28 | 0.04 | 0.13 | 7.93E-21 |
| SLFN5 | 7.59E-14 | -0.28 | 0.15 | 0.23 | 1.97E-09 |
| SMCO4 | 5.18E-235 | 0.74 | 0.52 | 0.08 | 1.35E-230 |
| SNAP23 | 2.02E-45 | 0.21 | 0.40 | 0.18 | 5.24E-41 |
| SOD1 | 1.98E-39 | 0.39 | 0.81 | 0.61 | 5.13E-35 |
| SPCS2 | 9.36E-34 | 0.23 | 0.78 | 0.54 | 2.43E-29 |
| SPINT2 | 1.60E-44 | 0.25 | 0.20 | 0.06 | 4.16E-40 |
| SPOCK2 | 1.13E-35 | 0.21 | 0.52 | 0.30 | 2.94E-31 |
| SRGN | 1.24E-48 | 0.40 | 0.98 | 0.90 | 3.22E-44 |
| SRSF7 | 8.89E-60 | -0.76 | 0.78 | 0.81 | 2.31E-55 |
| STK17A | 1.13E-31 | -0.41 | 0.70 | 0.72 | 2.93E-27 |
| STK17B | 4.24E-17 | -0.33 | 0.51 | 0.57 | 1.10E-12 |
| SUB1 | 3.09E-08 | -0.27 | 0.81 | 0.73 | 8.04E-04 |
| SYPL1 | 4.01E-49 | 0.23 | 0.45 | 0.21 | 1.04E-44 |
| SYTL3 | 3.08E-26 | -0.27 | 0.03 | 0.11 | 7.99E-22 |
| TAPBP | 2.57E-07 | -0.28 | 0.42 | 0.42 | 6.68E-03 |
| TBC1D4 | 1.39E-167 | 0.62 | 0.52 | 0.13 | 3.61E-163 |
| TCF7 | 7.26E-52 | 0.27 | 0.53 | 0.27 | 1.89E-47 |
| THADA | 1.49E-166 | 0.64 | 0.44 | 0.09 | 3.86E-162 |
| TIGIT | 3.07E-84 | 0.43 | 0.81 | 0.48 | 7.97E-80 |
| TLK1 | 1.13E-36 | 0.20 | 0.43 | 0.22 | 2.95E-32 |
| TMEM123 | 4.08E-129 | 0.57 | 0.64 | 0.25 | 1.06E-124 |
| TMEM70 | 3.56E-38 | 0.22 | 0.31 | 0.14 | 9.25E-34 |
| TMSB10 | 1.27E-77 | -0.49 | 0.99 | 0.98 | 3.29E-73 |
| TMSB4X | 1.30E-46 | -0.20 | 1.00 | 1.00 | 3.38E-42 |
| TNFAIP3 | 2.38E-18 | -0.46 | 0.22 | 0.31 | 6.18E-14 |
| TNFAIP8 | 7.57E-59 | 0.45 | 0.60 | 0.34 | 1.97E-54 |
| TNFRSF18 | 2.66E-80 | 0.47 | 0.36 | 0.12 | 6.91E-76 |
| TNFRSF25 | 1.23E-51 | 0.24 | 0.23 | 0.07 | 3.21E-47 |
| TNFRSF4 | 1.96E-169 | 0.80 | 0.56 | 0.15 | 5.09E-165 |
| TNFRSF9 | 4.35E-18 | -0.33 | 0.08 | 0.17 | 1.13E-13 |

| | | | | | |
|---|---|---|---|---|---|
| TOX2 | 7.11E-180 | 0.75 | 0.54 | 0.14 | 1.85E-175 |
| TPD52L1 | 3.19E-70 | 0.46 | 0.13 | 0.01 | 8.28E-66 |
| TPI1 | 3.54E-58 | 0.50 | 0.78 | 0.54 | 9.20E-54 |
| TPT1 | 1.46E-26 | -0.26 | 0.95 | 0.92 | 3.78E-22 |
| TRGC2 | 2.63E-34 | -0.68 | 0.09 | 0.23 | 6.82E-30 |
| TSEN54 | 4.12E-09 | -0.21 | 0.10 | 0.16 | 1.07E-04 |
| TSHZ2 | 6.88E-79 | 0.33 | 0.37 | 0.12 | 1.79E-74 |
| TUBA4A | 6.99E-08 | -0.35 | 0.42 | 0.42 | 1.82E-03 |
| TXNIP | 6.18E-13 | -0.37 | 0.34 | 0.40 | 1.60E-08 |
| UBB | 2.22E-28 | -0.31 | 0.96 | 0.94 | 5.76E-24 |
| UBC | 2.23E-12 | -0.21 | 0.99 | 0.98 | 5.80E-08 |
| UCP2 | 2.98E-33 | 0.33 | 0.64 | 0.43 | 7.73E-29 |
| UQCRC2 | 8.90E-68 | 0.37 | 0.56 | 0.27 | 2.31E-63 |
| VDAC1 | 1.64E-64 | 0.35 | 0.56 | 0.29 | 4.25E-60 |
| VIM | 3.63E-77 | -0.68 | 0.16 | 0.40 | 9.42E-73 |
| VOPP1 | 5.58E-57 | 0.30 | 0.46 | 0.22 | 1.45E-52 |
| XCL2 | 2.16E-15 | -0.24 | 0.05 | 0.12 | 5.60E-11 |
| XIST | 6.42E-26 | -0.32 | 0.06 | 0.16 | 1.67E-21 |
| YBX1 | 1.38E-25 | 0.22 | 0.88 | 0.71 | 3.59E-21 |
| YBX3 | 1.06E-12 | -0.22 | 0.06 | 0.12 | 2.75E-08 |
| YWHAQ | 1.75E-68 | 0.44 | 0.70 | 0.41 | 4.54E-64 |
| ZFP36 | 1.03E-40 | -0.65 | 0.63 | 0.70 | 2.67E-36 |
| ZFP36L2 | 2.01E-25 | -0.52 | 0.68 | 0.71 | 5.22E-21 |
| ZNF331 | 7.05E-09 | -0.36 | 0.41 | 0.43 | 1.83E-04 |
