## Supplementary Table 3 for "Dissecting intratumor heterogeneity of nodal B cell lymphomas on the transcriptional, genetic, and drug response level"

| Gene | p_val | avg_logFC | pct.1 | pct.2 | p_val_adj |
|---|---|---|---|---|---|
| IGHM | 4.65E-147 | 2.54 | 0.98 | 0.60 | 1.22E-142 |
| TCL1A | 4.57E-179 | 2.33 | 0.82 | 0.09 | 1.20E-174 |
| IGHD | 1.29E-167 | 1.88 | 0.87 | 0.18 | 3.39E-163 |
| FCER2 | 1.26E-69 | 1.20 | 0.65 | 0.25 | 3.31E-65 |
| IL4R | 7.03E-58 | 1.16 | 0.37 | 0.04 | 1.85E-53 |
| PLPP5 | 2.81E-40 | 1.05 | 0.40 | 0.12 | 7.39E-36 |
| CD72 | 1.75E-46 | 0.93 | 0.56 | 0.24 | 4.59E-42 |
| MEF2C | 6.49E-47 | 0.87 | 0.83 | 0.66 | 1.71E-42 |
| CD69 | 1.45E-40 | 0.86 | 0.93 | 0.84 | 3.82E-36 |
| HVCN1 | 8.31E-38 | 0.83 | 0.66 | 0.41 | 2.19E-33 |
| APLP2 | 5.92E-30 | 0.70 | 0.29 | 0.07 | 1.56E-25 |
| PHACTR1 | 9.39E-25 | 0.70 | 0.50 | 0.27 | 2.47E-20 |
| CD79B | 2.71E-54 | 0.69 | 0.98 | 0.93 | 7.12E-50 |
| SELENOH | 1.13E-23 | 0.68 | 0.44 | 0.22 | 2.96E-19 |
| TMSB10 | 1.67E-72 | 0.68 | 1.00 | 1.00 | 4.38E-68 |
| HIST1H1C | 4.92E-10 | 0.68 | 0.31 | 0.19 | 1.29E-05 |
| CXCR4 | 1.88E-24 | 0.67 | 0.92 | 0.86 | 4.96E-20 |
| LINC00926 | 5.13E-19 | 0.67 | 0.59 | 0.45 | 1.35E-14 |
| CHI3L2 | 5.59E-14 | 0.63 | 0.34 | 0.20 | 1.47E-09 |
| FOS | 2.46E-29 | 0.63 | 0.95 | 0.92 | 6.46E-25 |
| DBI | 2.61E-21 | 0.63 | 0.56 | 0.38 | 6.86E-17 |
| ADK | 6.76E-17 | 0.62 | 0.33 | 0.16 | 1.78E-12 |
| FCRL1 | 6.25E-19 | 0.61 | 0.33 | 0.16 | 1.64E-14 |
| CLEC2B | 1.17E-25 | 0.61 | 0.33 | 0.12 | 3.09E-21 |
| ISG20 | 2.36E-16 | 0.59 | 0.64 | 0.53 | 6.20E-12 |
| BTG1 | 1.62E-26 | 0.59 | 0.97 | 0.98 | 4.25E-22 |
| SNX29 | 1.73E-17 | 0.58 | 0.30 | 0.14 | 4.55E-13 |
| C1orf162 | 8.37E-22 | 0.57 | 0.43 | 0.21 | 2.20E-17 |
| TSPAN13 | 5.33E-18 | 0.56 | 0.47 | 0.29 | 1.40E-13 |
| NCF1 | 5.77E-29 | 0.55 | 0.79 | 0.63 | 1.52E-24 |
| YBX3 | 1.84E-22 | 0.54 | 0.22 | 0.05 | 4.84E-18 |
| AL139020.1 | 4.75E-22 | 0.54 | 0.15 | 0.01 | 1.25E-17 |
| CD79A | 1.26E-32 | 0.54 | 0.96 | 0.92 | 3.32E-28 |
| CD83 | 6.15E-11 | 0.54 | 0.47 | 0.33 | 1.62E-06 |
| FCMR | 9.34E-13 | 0.54 | 0.49 | 0.38 | 2.46E-08 |
| RHOH | 8.41E-16 | 0.54 | 0.57 | 0.41 | 2.21E-11 |
| H3F3A | 4.99E-32 | 0.53 | 0.93 | 0.87 | 1.31E-27 |
| CLEC2D | 9.59E-19 | 0.52 | 0.66 | 0.51 | 2.52E-14 |
| ABRACL | 3.03E-15 | 0.52 | 0.38 | 0.23 | 7.96E-11 |
| FAM129C | 8.80E-18 | 0.52 | 0.31 | 0.14 | 2.31E-13 |
| HLA-DMA | 7.15E-24 | 0.51 | 0.88 | 0.84 | 1.88E-19 |
| CDCA7L | 5.79E-19 | 0.51 | 0.26 | 0.10 | 1.52E-14 |
| BLOC1S2 | 1.92E-13 | 0.50 | 0.45 | 0.30 | 5.06E-09 |
| RNASE6 | 4.46E-15 | 0.50 | 0.39 | 0.22 | 1.17E-10 |
| NBEAL1 | 6.71E-16 | -0.52 | 0.59 | 0.74 | 1.76E-11 |
| SRGN | 5.41E-13 | -0.52 | 0.25 | 0.43 | 1.42E-08 |
| GLTSCR2 | 6.25E-15 | -0.52 | 0.26 | 0.45 | 1.64E-10 |
| MARCKS | 1.64E-26 | -0.53 | 0.06 | 0.25 | 4.31E-22 |
| VIM | 1.77E-28 | -0.53 | 0.30 | 0.60 | 4.65E-24 |
| LGALS1 | 1.01E-27 | -0.57 | 0.06 | 0.26 | 2.66E-23 |

| | | | | | |
|---|---|---|---|---|---|
| IGHGP | 2.57E-16 | -0.60 | 0.05 | 0.18 | 6.77E-12 |
| GZMK | 9.91E-09 | -0.64 | 0.10 | 0.20 | 2.61E-04 |
| CLECL1 | 1.32E-38 | -0.65 | 0.10 | 0.38 | 3.47E-34 |
| ITGB1 | 4.57E-22 | -0.65 | 0.09 | 0.27 | 1.20E-17 |
| AIM2 | 5.01E-49 | -0.68 | 0.04 | 0.32 | 1.32E-44 |
| CRIP1 | 4.72E-34 | -0.68 | 0.15 | 0.42 | 1.24E-29 |
| AC090498.1 | 1.23E-19 | -0.73 | 0.30 | 0.51 | 3.24E-15 |
| IL32 | 1.43E-08 | -0.75 | 0.12 | 0.22 | 3.77E-04 |
| IGHG3 | 7.40E-23 | -0.76 | 0.26 | 0.49 | 1.94E-18 |
| TNFRSF13B | 2.70E-54 | -0.82 | 0.10 | 0.45 | 7.10E-50 |
| GPR183 | 1.79E-45 | -0.97 | 0.23 | 0.57 | 4.70E-41 |
| LINC01781 | 6.31E-35 | -1.08 | 0.05 | 0.28 | 1.66E-30 |
| CD27 | 6.06E-102 | -1.17 | 0.11 | 0.63 | 1.59E-97 |
| IGHG1 | 2.25E-60 | -1.30 | 0.14 | 0.51 | 5.91E-56 |
| RP5-887A10.1 | 7.33E-62 | -1.32 | 0.01 | 0.32 | 1.93E-57 |
| AL928768.3 | 1.32E-28 | -1.41 | 0.01 | 0.17 | 3.47E-24 |
| IGKC | 6.25E-07 | -1.43 | 0.89 | 0.87 | 1.64E-02 |
| IGHA2 | 1.53E-44 | -3.78 | 0.02 | 0.26 | 4.01E-40 |
| IGHA1 | 1.96E-67 | -5.09 | 0.24 | 0.63 | 5.14E-63 |
