## Supplementary Table 4 for "Dissecting intratumor heterogeneity of nodal B cell lymphomas on the transcriptional, genetic, and drug response level"

| Protein_A | Protein_B | annotation_strategy | Source |
|---|---|---|---|
| A4 | FPR2 | CellPhoneDB, guidetopharmacology.org | |
| AA1R | ENTP1 | CellPhoneDB, curated | uniprot |
| AA2AR | ENTP1 | CellPhoneDB, curated | uniprot |
| AA2BR | ENTP1 | CellPhoneDB, curated | uniprot |
| AA3R | ENTP1 | CellPhoneDB, curated | uniprot |
| ACE2 | GHRL | CellPhoneDB, I2D | |
| ACHA7 | SLUR1 | CellPhoneDB, IMEx,InnateDB-All,UniProt | |
| ACKR1 | CCL17 | CellPhoneDB, I2D | |
| ACKR2 | CCL2 | CellPhoneDB, curated | PMID: 24218476 |
| ACKR2 | CCL14 | CellPhoneDB, curated | PMID: 24218476 |
| ACKR2 | CCL13 | CellPhoneDB, curated | PMID: 24218476 |
| ACKR2 | CL3L1 | CellPhoneDB, I2D | |
| ACKR2 | CCL11 | CellPhoneDB, curated | PMID: 24218476 |
| ACKR2 | CCL28 | CellPhoneDB, I2D | |
| ACKR2 | CCL7 | CellPhoneDB, curated | PMID: 24218476 |
| ACKR2 | CCL5 | CellPhoneDB, curated | PMID: 24218476 |
| ACKR2 | CCL4 | CellPhoneDB, curated | PMID: 24218476 |
| ACKR2 | CCL8 | CellPhoneDB, curated | PMID: 24218476 |
| ACKR2 | CCL27 | CellPhoneDB, I2D | |
| ACKR3 | SDF1 | CellPhoneDB, guidetopharmacology.org | |
| ADIPO | MO2R1 | CellPhoneDB, IMEx,IntAct | |
| ADIPO | PKR2 | CellPhoneDB, IMEx,IntAct | |
| ADIPO | CLC2D | CellPhoneDB, IMEx,IntAct | |
| ADIPO | GP152 | CellPhoneDB, IMEx,IntAct | |
| ADML | MRGX2 | CellPhoneDB, guidetopharmacology.org | |
| AGRG5 | FAM3C | CellPhoneDB, InnateDB-All | |
| AGRL1 | NRG1 | CellPhoneDB, InnateDB-All | |
| AGRP | MC3R | CellPhoneDB, guidetopharmacology.org | |
| AGRP | MC4R | CellPhoneDB, guidetopharmacology.org | |
| AGRP | MC5R | CellPhoneDB, guidetopharmacology.org | |
| ANF | ANPRB | CellPhoneDB, I2D,InnateDB-All | |
| ANF | ANPRA | CellPhoneDB, guidetopharmacology.org | |
| ANF | ANPRC | CellPhoneDB, I2D,InnateDB-All | |
| ANFB | ANPRC | CellPhoneDB, I2D,InnateDB-All | |
| ANFB | DPP4 | CellPhoneDB, InnateDB-All,MINT | |
| ANFB | ANPRB | CellPhoneDB, I2D,InnateDB-All | |
| ANGP2 | TIE2 | CellPhoneDB, curated | uniprot |
| ANGT | AGTR1 | CellPhoneDB, guidetopharmacology.org | |
| ANGT | AGTR2 | CellPhoneDB, guidetopharmacology.org | |
| ANPRA | ANFB | CellPhoneDB, guidetopharmacology.org | |
| ANPRA | ANFC | CellPhoneDB, InnateDB-All | |
| ANPRB | ANFC | CellPhoneDB, guidetopharmacology.org | |
| ANPRC | OSTN | CellPhoneDB, guidetopharmacology.org | |
| ANPRC | ANFC | CellPhoneDB, I2D,InnateDB-All | |
| APJ | APEL | CellPhoneDB, guidetopharmacology.org | |
| ASGR2 | ADIPO | CellPhoneDB, IMEx,IntAct | |
| AVR2A | INHBE | CellPhoneDB, curated | PMID: 22710174 |
| AVR2A | INHBC | CellPhoneDB, curated | PMID: 22710174 |
| BDNF | JAM1 | CellPhoneDB, IMEx,InnateDB-All,IntAct | |
| BDNF | GP152 | CellPhoneDB, IMEx,IntAct | |
| BDNF | SORT | CellPhoneDB, I2D,IMEx,InnateDB-All,IntAct | |

| | | | |
|---|---|---|---|
| BDNF | NTRK2 | CellPhoneDB, curated | uniprot |
| BMP10 | KI3L1 | CellPhoneDB, IMEx,IntAct | |
| BMP10 | KI2L3 | CellPhoneDB, IMEx,IntAct | |
| BMP10 | GI24 | CellPhoneDB, IMEx,IntAct | |
| BMP10 | KI3S1 | CellPhoneDB, IMEx,IntAct | |
| BMP10 | GP152 | CellPhoneDB, IMEx,IntAct | |
| BMP2 | SMO | CellPhoneDB, I2D | |
| BMP8B | UPAR | CellPhoneDB, InnateDB-All | |
| BTLA | TNR14 | CellPhoneDB, guidetopharmacology.org | |
| BY55 | TNR14 | CellPhoneDB, curated | PMID: 21959263 |
| C5AR1 | RS19 | CellPhoneDB, guidetopharmacology.org | |
| CADH1 | KLRG1 | CellPhoneDB, curated | PMC: 3030123 |
| CADM3 | PVRL3 | CellPhoneDB, curated | uniprot |
| CADM3 | CADM4 | CellPhoneDB, curated | uniprot |
| CADM3 | E41L1 | CellPhoneDB, curated | uniprot |
| CALC | CALCR | CellPhoneDB, guidetopharmacology.org | |
| CALCA | CALCR | CellPhoneDB, guidetopharmacology.org | |
| CALCB | CALCR | CellPhoneDB, guidetopharmacology.org | |
| CALCR | ADM2 | CellPhoneDB, guidetopharmacology.org | |
| CALCR | ADML | CellPhoneDB, guidetopharmacology.org | |
| CC4L | GP151 | CellPhoneDB, IMEx,IntAct | |
| CC4L | GI24 | CellPhoneDB, IMEx,IntAct | |
| CC4L | GP152 | CellPhoneDB, IMEx,IntAct | |
| CC4L | GP101 | CellPhoneDB, IMEx,IntAct | |
| CCKN | GASR | CellPhoneDB, guidetopharmacology.org | |
| CCKN | CCKAR | CellPhoneDB, guidetopharmacology.org | |
| CCL1 | CCR8 | CellPhoneDB, curated | PMID: 24218476 |
| CCL11 | ACKR4 | CellPhoneDB, I2D | |
| CCL11 | CCR3 | CellPhoneDB, curated | PMID: 24218476 |
| CCL16 | CCR1 | CellPhoneDB, curated | PMID: 24218476 |
| CCL16 | CCR8 | CellPhoneDB, I2D | |
| CCL16 | HRH4 | CellPhoneDB, guidetopharmacology.org | |
| CCL16 | CCR2 | CellPhoneDB, curated | PMID: 24218476 |
| CCL19 | ACKR4 | CellPhoneDB, curated | PMID: 24218476 |
| CCL2 | CCR10 | CellPhoneDB, I2D | |
| CCL2 | CCR2 | CellPhoneDB, curated | PMID: 24218476 |
| CCL2 | ACKR1 | CellPhoneDB, curated | PMID: 24218476 |
| CCL21 | CCR7 | CellPhoneDB, curated | PMID: 24218476 |
| CCL21 | ACKR4 | CellPhoneDB, curated | PMID: 24218476 |
| CCL22 | CCR4 | CellPhoneDB, curated | PMID: 24218476 |
| CCL22 | DPP4 | CellPhoneDB, curated | PMID: 24218476 |
| CCL24 | CCR3 | CellPhoneDB, curated | PMID: 24218476 |
| CCL24 | CCR2 | CellPhoneDB, guidetopharmacology.org | |
| CCL25 | CCR9 | CellPhoneDB, curated | PMID: 24218476 |
| CCL25 | ACKR4 | CellPhoneDB, curated | PMID: 24218476 |
| CCL4 | CCR5 | CellPhoneDB, curated | uniprot |
| CCL4 | CCR8 | CellPhoneDB, I2D | |
| CCL4 | CNR2 | CellPhoneDB, IMEx,IntAct | |
| CCL4 | GPC5D | CellPhoneDB, IMEx,IntAct | |
| CCL4 | GP152 | CellPhoneDB, IMEx,IntAct | |
| CCL4 | CTR1 | CellPhoneDB, IMEx,IntAct | |
| CCL5 | CCR4 | CellPhoneDB, curated | PMID: 24218476 |

| | | | |
|---|---|---|---|
| CCL5 | CCR5 | CellPhoneDB, curated | PMID: 24218476 |
| CCL5 | ACKR1 | CellPhoneDB, curated | PMID: 24218476 |
| CCL5 | CCR1 | CellPhoneDB, curated | PMID: 24218476 |
| CCL5 | ACKR4 | CellPhoneDB, I2D | |
| CCL5 | CCR3 | CellPhoneDB, curated | PMID: 24218476 |
| CCL7 | ACKR1 | CellPhoneDB, I2D | |
| CCL8 | ACKR1 | CellPhoneDB, I2D | |
| CCR1 | CCL18 | CellPhoneDB, guidetopharmacology.org | |
| CCR1 | CCL26 | CellPhoneDB, curated | PMID: 24218476 |
| CCR1 | CCL14 | CellPhoneDB, curated | PMID: 24218476 |
| CCR1 | CCL7 | CellPhoneDB, curated | PMID: 24218476 |
| CCR1 | CCL23 | CellPhoneDB, curated | PMID: 24218476 |
| CCR1 | CCL13 | CellPhoneDB, guidetopharmacology.org | |
| CCR1 | CCL15 | CellPhoneDB, curated | uniprot |
| CCR1 | CCL8 | CellPhoneDB, curated | PMID: 24218476 |
| CCR10 | CCL7 | CellPhoneDB, I2D | |
| CCR10 | CCL28 | CellPhoneDB, guidetopharmacology.org | |
| CCR10 | CCL27 | CellPhoneDB, curated | PMID: 24218476 |
| CCR2 | CCL7 | CellPhoneDB, curated | PMID: 24218476 |
| CCR2 | CCL26 | CellPhoneDB, curated | PMID: 24218476 |
| CCR2 | CCL8 | CellPhoneDB, curated | PMID: 24218476 |
| CCR2 | CCL13 | CellPhoneDB, curated | PMID: 24218476 |
| CCR2 | CCL11 | CellPhoneDB, guidetopharmacology.org | |
| CCR3 | CCL15 | CellPhoneDB, curated | uniprot |
| CCR3 | CCL28 | CellPhoneDB, guidetopharmacology.org | |
| CCR3 | CCL26 | CellPhoneDB, curated | PMID: 24218476 |
| CCR3 | CCL7 | CellPhoneDB, curated | PMID: 24218476 |
| CCR3 | CCL8 | CellPhoneDB, curated | PMID: 24218476 |
| CCR3 | CCL13 | CellPhoneDB, curated | PMID: 24218476 |
| CCR3 | CCL14 | CellPhoneDB, curated | PMID: 24218476 |
| CCR4 | CCL17 | CellPhoneDB, curated | PMID: 24218476 |
| CCR5 | CCL7 | CellPhoneDB, curated | PMID: 24218476 |
| CCR5 | CCL8 | CellPhoneDB, curated | PMID: 24218476 |
| CCR6 | CCL20 | CellPhoneDB, curated | PMID: 24218476 |
| CCR7 | CCL19 | CellPhoneDB, curated | PMID: 24218476 |
| CCR8 | CCL18 | CellPhoneDB, curated | PMID: 24218476 |
| CCRL2 | CCL19 | CellPhoneDB, guidetopharmacology.org | |
| CD1D | LIRB2 | CellPhoneDB, curated | PMID: 19124746 |
| CD2 | LFA3 | CellPhoneDB, curated | PMID: 23602662 |
| CD226 | PVRL2 | CellPhoneDB, curated | PMID: 15607800, PMID: 2444014[illegible] |
| CD27 | CD70 | CellPhoneDB, curated | PMID: 26697006 |
| CD28 | CD80 | CellPhoneDB, curated | PMID: 23954143 |
| CD28 | CD86 | CellPhoneDB, curated | PMID: 23954143 |
| CD40LG | CD40 | curated | uniprot |
| CD47 | SIRPG | CellPhoneDB, curated | PMID: 15294972 |
| CD48 | CD244 | CellPhoneDB, curated | uniprot & PMID: 26697006 |
| CD52 | SIG10 | CellPhoneDB, curated | PMID: 23685786 |
| CD6 | CD166 | CellPhoneDB, curated | PMID: 23602662 |
| CD70 | GPC5B | CellPhoneDB, InnateDB-All | |
| CD70 | TNR17 | CellPhoneDB, InnateDB-All | |
| CD72 | SEM4D | CellPhoneDB, curated | PMID: 22325954 |
| CD80 | CTLA4 | CellPhoneDB, curated | PMID: 23954143 |

| | | | |
|---|---|---|---|
| CD80 | PD1L1 | CellPhoneDB, curated | PMID: 17629517,PMID: 18585785 |
| CD86 | CTLA4 | CellPhoneDB, curated | PMID: 23954143 |
| CD8A | CEAM5 | CellPhoneDB, curated | PMID: 24104458 |
| CEAM1 | CD209 | CellPhoneDB, curated | PMID: 16282604,PMID: 16246332 |
| CEAM1 | CEAM6 | CellPhoneDB, curated | PMID: 21982860 |
| CEAM1 | LYAM2 | CellPhoneDB, curated | PMID: 1378450 |
| CEAM1 | CEAM8 | CellPhoneDB, curated | PMID: 24743304 |
| CEAM5 | CEAM6 | CellPhoneDB, curated | uniprot |
| CEAM5 | CD1D | CellPhoneDB, curated | PMID: 24104458 |
| CEAM5 | CEAM1 | CellPhoneDB, curated | PMID:24987108 |
| CEAM6 | CEAM6 | CellPhoneDB, curated | PMID: 11590190 |
| CEAM8 | CEAM6 | CellPhoneDB, curated | PMID: 11590190 |
| CER1 | MRC2 | CellPhoneDB, InnateDB-All | |
| CL3L1 | DPP4 | CellPhoneDB, I2D | |
| CL3L1 | CCR1 | CellPhoneDB, curated | PMID: 24218476 |
| CL3L1 | CCR3 | CellPhoneDB, curated | PMID: 24218476 |
| CLC2B | KLRF1 | CellPhoneDB, curated | PMID: 24223577 |
| CML1 | RARR2 | CellPhoneDB, guidetopharmacology.org | |
| CNR2 | ADIPO | CellPhoneDB, IMEx,IntAct | |
| CO3 | C3AR | CellPhoneDB, guidetopharmacology.org | |
| CO5 | C5AR1 | CellPhoneDB, guidetopharmacology.org | |
| CO5 | C5AR2 | CellPhoneDB, guidetopharmacology.org | |
| COLI | MSHR | CellPhoneDB, guidetopharmacology.org | |
| COLI | MC4R | CellPhoneDB, guidetopharmacology.org | |
| COLI | MC3R | CellPhoneDB, guidetopharmacology.org | |
| COLI | MC5R | CellPhoneDB, guidetopharmacology.org | |
| COLI | ACTHR | CellPhoneDB, I2D | |
| CORT | GHSR | CellPhoneDB, I2D | |
| CORT | MRGX2 | CellPhoneDB, I2D | |
| CRF | CRFR2 | CellPhoneDB, I2D,InnateDB-All | |
| CRFR2 | UCN3 | CellPhoneDB, guidetopharmacology.org | |
| CRFR2 | UCN2 | CellPhoneDB, guidetopharmacology.org | |
| CSF3 | CSF3R | CellPhoneDB, curated | uniprot |
| CX3C1 | X3CL1 | CellPhoneDB, curated | PMID: 24218476 |
| CXAR | FAM3C | CellPhoneDB, IMEx,IntAct | |
| CXCL13 | CXCR5 | curated | KEGG |
| CXCL13 | CXCR7 | curated | KEGG |
| CXCL2 | CXCR1 | CellPhoneDB, I2D | |
| CXCL2 | DPP4 | CellPhoneDB, curated | PMID: 24218476 |
| CXCL2 | CXCR2 | CellPhoneDB, curated | PMID: 24218476 |
| CXCL3 | CXCR2 | CellPhoneDB, curated | PMID: 24218476 |
| CXCL3 | CXCR1 | CellPhoneDB, I2D | |
| CXCL5 | ACKR1 | CellPhoneDB, I2D | |
| CXCL7 | CXCR1 | CellPhoneDB, curated | PMID: 24218476 |
| CXCL7 | CXCR2 | CellPhoneDB, curated | PMID: 24218476 |
| CXCR1 | CXCL6 | CellPhoneDB, curated | PMID: 24218476 |
| CXCR1 | CXCL5 | CellPhoneDB, curated | PMID: 24218476 |
| CXCR1 | SYYC | CellPhoneDB, guidetopharmacology.org | |
| CXCR2 | CXCL5 | CellPhoneDB, curated | PMID: 24218476 |
| CXCR2 | CXCL6 | CellPhoneDB, curated | PMID: 24218476 |
| CXCR3 | CCL19 | CellPhoneDB, guidetopharmacology.org | |
| CXCR3 | CXCL9 | CellPhoneDB, curated | PMID: 24218476 |

| | | | |
|---|---|---|---|
| CXCR3 | CCL20 | CellPhoneDB, guidetopharmacology.org | |
| CXCR6 | CXL16 | CellPhoneDB, guidetopharmacology.org | |
| CXL10 | CXCR3 | CellPhoneDB, curated | PMID: 24218476 |
| CXL10 | DPP4 | CellPhoneDB, curated | PMID: 24218476 |
| CXL11 | ACKR3 | CellPhoneDB, guidetopharmacology.org | |
| CXL11 | CXCR3 | CellPhoneDB, curated | PMID: 24218476 |
| CXL11 | DPP4 | CellPhoneDB, curated | PMID: 24218476 |
| CXL13 | ACKR4 | CellPhoneDB, curated | PMID: 24218476 |
| CXL13 | CXCR5 | CellPhoneDB, curated | PMID: 24218476 |
| DAF | CD97 | CellPhoneDB, curated | PMID: 11297558 |
| DLK1 | NOTC3 | CellPhoneDB, curated | PMID: 22353464 |
| DLK1 | NOTC4 | CellPhoneDB, curated | PMID: 22353464 |
| DLK1 | NOTC2 | CellPhoneDB, curated | PMID: 22353464 |
| DLL1 | NOTC2 | CellPhoneDB, curated | PMID: 22353464 |
| DLL1 | NOTC3 | CellPhoneDB, curated | PMID: 22353464 |
| DLL1 | NOTC1 | CellPhoneDB, curated | PMID: 22353464 |
| DLL1 | NOTC4 | CellPhoneDB, curated | PMID: 22353464 |
| DLL3 | NOTC3 | CellPhoneDB, curated | PMID: 22353464 |
| DLL4 | NOTC3 | CellPhoneDB, curated | PMID: 22353464 |
| DPP4 | SDF1 | CellPhoneDB, curated | PMID: 24218476 |
| DPP4 | CXCL9 | CellPhoneDB, curated | PMID: 24218476 |
| DPP4 | CCL11 | CellPhoneDB, curated | PMID: 24218476 |
| DSC1 | DSG2 | CellPhoneDB, curated | PMID: 27298358 |
| DSC2 | DSG2 | CellPhoneDB, curated | PMID: 27298358 |
| DSG1 | DSC2 | CellPhoneDB, curated | PMID: 27298358 |
| DSG1 | DSC1 | CellPhoneDB, curated | PMID: 27298358 |
| DSG1 | DSC3 | CellPhoneDB, curated | PMID: 27298358 |
| DSG2 | DSC3 | CellPhoneDB, curated | PMID: 27298358 |
| EDA | TNR27 | CellPhoneDB, guidetopharmacology.org | |
| EDA | EDAR | CellPhoneDB, guidetopharmacology.org | |
| EDN2 | EDNRA | CellPhoneDB, guidetopharmacology.org | |
| EFNA2 | EPHA7 | CellPhoneDB, curated | PMID: 15114347 |
| EFNA2 | EPHA8 | CellPhoneDB, curated | PMID: 15114347 |
| EFNA2 | EPHA2 | CellPhoneDB, curated | PMID: 15114347 |
| EFNA2 | EPHA3 | CellPhoneDB, curated | PMID: 15114347 |
| EFNA2 | EPHA5 | CellPhoneDB, curated | PMID: 15114347 |
| EGF | NRG1 | CellPhoneDB, I2D | |
| EGFR | TGFB1 | CellPhoneDB, IMEx,InnateDB-All,MINT | |
| EGFR | MIF | CellPhoneDB, IMEx,InnateDB-All | |
| EGFR | EPGN | CellPhoneDB, curated | uniprot |
| EGFR | BTC | CellPhoneDB, curated | uniprot |
| EGFR | NRG1 | CellPhoneDB, I2D,IMEx,InnateDB-All,IntAct | |
| EGFR | CNTF | CellPhoneDB, InnateDB-All | |
| EGFR | TGFA | CellPhoneDB, curated | uniprot |
| EGFR | EGF | CellPhoneDB, curated | uniprot |
| ELA | APJ | CellPhoneDB, guidetopharmacology.org | |
| EPHB3 | EFNB3 | CellPhoneDB, curated | PMID: 15114347 |
| EPO | EPOR | CellPhoneDB, curated | uniprot |
| EPOR | SCF | CellPhoneDB, I2D | |
| ERBB3 | BTC | CellPhoneDB, curated | uniprot |
| ERBB3 | NRG1 | CellPhoneDB, curated | uniprot |
| EREG | EGFR | CellPhoneDB, curated | uniprot |

| | | | |
|---|---|---|---|
| ETBR2 | SAP | CellPhoneDB, guidetopharmacology.org | |
| FAM3C | CLC2D | CellPhoneDB, IMEx,InnateDB-All,IntAct | |
| FASLG | FAS | uniprot | |
| FCG2A | CXCL9 | CellPhoneDB, IMEx,IntAct | |
| FFAR2 | CC4L | CellPhoneDB, IMEx,IntAct | |
| FFAR2 | FAM3C | CellPhoneDB, IMEx,IntAct | |
| FFAR2 | TNFA | CellPhoneDB, IMEx,IntAct | |
| FFAR2 | BMP10 | CellPhoneDB, IMEx,IntAct | |
| FGF1 | FGFR3 | CellPhoneDB, I2D,IMEx,InnateDB-All,IntAct | |
| FGF1 | TGBR3 | CellPhoneDB, InnateDB-All | |
| FGF1 | FGFR4 | CellPhoneDB, I2D,IMEx,InnateDB-All,IntAct | |
| FGF17 | FGFR4 | CellPhoneDB, I2D | |
| FGF17 | FGFR3 | CellPhoneDB, I2D | |
| FGF19 | FGFR4 | CellPhoneDB, I2D,IMEx,InnateDB-All | |
| FGF3 | FGFR4 | CellPhoneDB, I2D | |
| FGF3 | FGFR3 | CellPhoneDB, I2D | |
| FGF4 | FGFR3 | CellPhoneDB, I2D | |
| FGF4 | FGFR4 | CellPhoneDB, I2D | |
| FGF5 | FGFR4 | CellPhoneDB, I2D,IMEx,InnateDB-All,IntAct | |
| FGF5 | FGFR3 | CellPhoneDB, I2D | |
| FGF6 | FGFR3 | CellPhoneDB, I2D | |
| FGF6 | FGFR4 | CellPhoneDB, I2D | |
| FGFR3 | PVRL1 | CellPhoneDB, InnateDB-All | |
| FGFR3 | FGF9 | CellPhoneDB, I2D,InnateDB-All | |
| FGFR3 | FGF23 | CellPhoneDB, I2D | |
| FGFR3 | FGF8 | CellPhoneDB, I2D,IMEx,InnateDB-All | |
| FGFR4 | SG3A1 | CellPhoneDB, IMEx,InnateDB-All,MINT | |
| FGFR4 | FGF8 | CellPhoneDB, I2D | |
| FGFR4 | FGF9 | CellPhoneDB, I2D | |
| FGFR4 | PTPRR | CellPhoneDB, InnateDB-All | |
| FLT3 | FLT3L | CellPhoneDB, curated | uniprot |
| FPR2 | CAMP | CellPhoneDB, guidetopharmacology.org | |
| FPR3 | HEBP1 | CellPhoneDB, guidetopharmacology.org | |
| FSHB | FSHR | CellPhoneDB, I2D,IMEx,InnateDB-All,IntAct | |
| FZD6 | WNT5A | CellPhoneDB, curated | PMID: 24032637 |
| FZD6 | WNT4 | CellPhoneDB, curated | PMID: 24032637 |
| FZD7 | WNT3 | CellPhoneDB, curated | PMID: 24032637 |
| FZD9 | WNT1 | CellPhoneDB, I2D | |
| FZD9 | WNT2 | CellPhoneDB, curated | PMID: 24032637 |
| GALA | GP151 | CellPhoneDB, I2D | |
| GALA | GALR1 | CellPhoneDB, guidetopharmacology.org | |
| GALR1 | GALP | CellPhoneDB, guidetopharmacology.org | |
| GALR2 | GALP | CellPhoneDB, guidetopharmacology.org | |
| GALR2 | GALA | CellPhoneDB, guidetopharmacology.org | |
| GALR3 | GALA | CellPhoneDB, guidetopharmacology.org | |
| GALR3 | GALP | CellPhoneDB, guidetopharmacology.org | |
| GAST | KI2L3 | CellPhoneDB, IMEx,IntAct | |
| GAST | GASR | CellPhoneDB, guidetopharmacology.org | |
| GAST | GP152 | CellPhoneDB, IMEx,IntAct | |
| GDF1 | 5HT4R | CellPhoneDB, IMEx,MINT | |
| GDF11 | ANTR1 | CellPhoneDB, InnateDB-All | |
| GFRAL | GDF15 | CellPhoneDB, guidetopharmacology.org | |

| | | | |
|---|---|---|---|
| GHRHR | GHRL | CellPhoneDB, I2D | |
| GHSR | GHRL | CellPhoneDB, guidetopharmacology.org | |
| GHSR | LEAP2 | CellPhoneDB, guidetopharmacology.org | |
| GIP | DPP4 | CellPhoneDB, InnateDB-All,MINT | |
| GLHA | LSHR | CellPhoneDB, I2D,InnateDB-All | |
| GLHA | FSHR | CellPhoneDB, I2D,InnateDB-All | |
| GLP2R | GLUC | CellPhoneDB, guidetopharmacology.org | |
| GLRA2 | FAM3C | CellPhoneDB, InnateDB-All | |
| GLUC | DPP4 | CellPhoneDB, InnateDB-All,MINT | |
| GLUC | GLR | CellPhoneDB, I2D | |
| GON1 | GNRHR | CellPhoneDB, I2D | |
| GON2 | GNRHR | CellPhoneDB, guidetopharmacology.org | |
| GPHA2 | EPHA6 | CellPhoneDB, InnateDB-All | |
| GPR1 | RARR2 | CellPhoneDB, guidetopharmacology.org | |
| GPR19 | ENHO | CellPhoneDB, guidetopharmacology.org | |
| GPR25 | RETN | CellPhoneDB, IMEx,IntAct | |
| GPR37 | SAP | CellPhoneDB, guidetopharmacology.org | |
| GPR42 | CC4L | CellPhoneDB, IMEx,IntAct | |
| GPR42 | FAM3C | CellPhoneDB, IMEx,IntAct | |
| GPR42 | ADIPO | CellPhoneDB, IMEx,IntAct | |
| GPR75 | CCL5 | CellPhoneDB, guidetopharmacology.org | |
| GPR98 | GPHA2 | CellPhoneDB, InnateDB-All | |
| GROA | ACKR1 | CellPhoneDB, curated | PMID: 24218476 |
| GROA | CXCR2 | CellPhoneDB, curated | PMID: 24218476 |
| GROA | CXCR1 | CellPhoneDB, curated | PMID: 24218476 |
| GRP | GRPR | CellPhoneDB, guidetopharmacology.org | |
| GUC2C | GUC2B | CellPhoneDB, guidetopharmacology.org | |
| GUC2C | GUC2A | CellPhoneDB, guidetopharmacology.org | |
| HG2A | A4 | CellPhoneDB, curated | PMID: 19849849 |
| HG2A | MIF | CellPhoneDB, I2D,InnateDB-All | |
| HGFL | RON | CellPhoneDB, curated | PMID: 23792360 |
| HLAA | KI3L1 | CellPhoneDB, curated | uniprot |
| HLAB | KI3L2 | CellPhoneDB, curated | uniprot |
| HLAC | FAM3C | CellPhoneDB, InnateDB-All | |
| HLAC | KI2L3 | CellPhoneDB, curated | PMID: 28484462 |
| HLAC | KI2L1 | CellPhoneDB, curated | PMID: 28484462 |
| HLAC | TSHB | CellPhoneDB, InnateDB-All | |
| HLADPA1 | TNFL9 | CellPhoneDB, InnateDB-All | |
| HLADPA1 | GALA | CellPhoneDB, InnateDB-All | |
| HLADPB1 | TN13B | CellPhoneDB, InnateDB-All | |
| HLADPB1 | NRG1 | CellPhoneDB, InnateDB-All | |
| HLADQB2 | PRL | CellPhoneDB, InnateDB-All | |
| HLADRB1 | MIME | CellPhoneDB, InnateDB-All | |
| HLAE | NKG2D | CellPhoneDB, curated | PMID: 24223577 |
| HLAE | NKG2A | CellPhoneDB, curated | Induces NK Cell Inhibition. Reviewe |
| HLAE | NKG2C | CellPhoneDB, curated | uniprot & PMID: 26697006 |
| HLAF | LIRB1 | CellPhoneDB, curated | PMID: 11169396 |
| HLAF | KI3S1 | CellPhoneDB, curated | PMID: 27455421 |
| HLAF | KI3L1 | CellPhoneDB, curated | PMID: 27455421 |
| HLAF | LIRB2 | CellPhoneDB, curated | PMID: 11169396 |
| HLAF | KI3L2 | CellPhoneDB, curated | PMID: 27455421 |
| HLAG | LIRB1 | CellPhoneDB, curated | PMID:24987108 |

| | | | |
|---|---|---|---|
| HLAG | LIRB2 | CellPhoneDB, curated | uniprot |
| IAPP | CALCR | CellPhoneDB, guidetopharmacology.org | |
| ICAM3 | CLC4M | CellPhoneDB, curated | PMID: 15795245 |
| ICAM3 | CD209 | CellPhoneDB, curated | PMID: 15795245 |
| ICOSL | ICOS | CellPhoneDB, curated | PMID: 23954143 |
| IFNA8 | GP152 | CellPhoneDB, IMEx,IntAct | |
| IFNE | GPR98 | CellPhoneDB, InnateDB-All | |
| IGF1 | IGF1R | CellPhoneDB, curated | uniprot |
| IGF2 | GP152 | CellPhoneDB, IMEx,IntAct | |
| IGF2 | MPRI | CellPhoneDB, curated | uniprot |
| IGF2 | IGF1R | CellPhoneDB, curated | PMID: 27102148 |
| IGFL1 | IGFR1 | CellPhoneDB, IMEx,InnateDB-All,UniProt | |
| IGFL2 | IGFR1 | CellPhoneDB, IMEx,InnateDB-All,UniProt | |
| IGFL3 | IGFR1 | CellPhoneDB, IMEx,InnateDB-All,UniProt | |
| IL10 | IL10RA | CellPhoneDB, uniprot | |
| IL10 | IL10RB | uniprot | |
| IL11 | IL11RA | curated | uniprot |
| IL13 | IL13RA1 | curated | uniprot |
| IL13 | TM219 | CellPhoneDB, IMEx,IntAct | |
| IL13 | I13R2 | CellPhoneDB, curated | PMID: 26471366 |
| IL15 | IL15RA | curated | uniprot |
| IL15 | IL15RB | curated | uniprot |
| IL15 | IL2RG | curated | uniprot |
| IL15 | I15RA | CellPhoneDB, guidetopharmacology.org | |
| IL16 | NMDE3 | CellPhoneDB, I2D | |
| IL1B | IL1R1 | curated | uniprot |
| IL1B | IL1R2 | curated | uniprot |
| IL2 | IL2RA | curated | uniprot |
| IL2 | IL2RB | curated | uniprot |
| IL2 | IL2RG | curated | uniprot |
| IL20 | IL20RA | curated | uniprot |
| IL21 | IL21R | CellPhoneDB, uniprot | |
| IL3 | IL3RA | curated | uniprot |
| IL36A | IL1RL2 | curated | uniprot |
| IL37 | IRPL1 | CellPhoneDB, InnateDB | |
| IL4 | IL13RA1 | CellPhoneDB, uniprot | |
| IL4 | IL4R | CellPhoneDB, uniprot | |
| IL4 | I13R2 | CellPhoneDB, I2D | |
| IL5 | IL5R | curated | uniprot |
| IL6 | IL6R | CellPhoneDB, uniprot | |
| IL6 | IL6ST | CellPhoneDB, uniprot | |
| IL6 | HRH1 | CellPhoneDB, IMEx,InnateDB-All,IntAct | |
| IL8 | CXCR2 | CellPhoneDB, curated | PMID: 24218476 |
| IL8 | CXCR1 | CellPhoneDB, curated | PMID: 24218476 |
| IL8 | ACKR1 | CellPhoneDB, curated | PMID: 24218476 |
| INHA | TGBR3 | CellPhoneDB, I2D | |
| INHBC | AVR2B | CellPhoneDB, curated | PMID: 22710174 |
| INHBE | AVR2B | CellPhoneDB, curated | PMID: 22710174 |
| INS | LIRB2 | CellPhoneDB, IMEx,IntAct | |
| INS | INSR | CellPhoneDB, curated | uniprot |
| INS | LIRB1 | CellPhoneDB, IMEx,IntAct | |
| INSL3 | RXFP2 | CellPhoneDB, guidetopharmacology.org | |

| | | | |
|---|---|---|---|
| INSL3 | RXFP1 | CellPhoneDB, guidetopharmacology.org | |
| JAG1 | NOTC2 | CellPhoneDB, curated | PMID: 22353464 |
| JAG1 | NOTC4 | CellPhoneDB, curated | PMID: 22353464 |
| JAG1 | NOTC3 | CellPhoneDB, curated | PMID: 22353464 |
| KI2L3 | FAM3C | CellPhoneDB, IMEx,IntAct | |
| KI2L3 | CXCL9 | CellPhoneDB, IMEx,IntAct | |
| KISS1 | KISSR | CellPhoneDB, guidetopharmacology.org | |
| KIT | SCF | CellPhoneDB, curated | uniprot |
| KLRB1 | CLC2D | CellPhoneDB, curated | PMID: 24223577 |
| KLRF2 | CLC2A | CellPhoneDB, curated | PMID: 24223577 |
| KLRG2 | WNT5B | CellPhoneDB, InnateDB-All | |
| KLRG2 | WNT11 | CellPhoneDB, InnateDB-All | |
| KLRG2 | TNFL9 | CellPhoneDB, InnateDB-All | |
| KNG1 | BKRB1 | CellPhoneDB, guidetopharmacology.org | |
| L1CAM | L1CAM | CellPhoneDB, curated | PMID: 18701456 |
| LAIR1 | LIRB4 | CellPhoneDB, curated | PMID: 19283782/ |
| LAMP1 | FAM3C | CellPhoneDB, InnateDB-All | |
| LAMP1 | VSTM1 | CellPhoneDB, IMEx,IntAct | |
| LEG9 | AAAT | CellPhoneDB, I2D,InnateDB-All | |
| LEG9 | CD47 | CellPhoneDB, InnateDB-All | |
| LEG9 | LRP1 | CellPhoneDB, InnateDB-All | |
| LEG9 | MRC2 | CellPhoneDB, InnateDB-All | |
| LEG9 | COL12 | CellPhoneDB, InnateDB-All | |
| LEG9 | SORL | CellPhoneDB, InnateDB-All | |
| LEG9 | MET | CellPhoneDB, InnateDB-All | |
| LEG9 | DAG1 | CellPhoneDB, InnateDB-All | |
| LEG9 | HAVR2 | CellPhoneDB, curated | PMID: 27192565 |
| LEUK | SN | CellPhoneDB, curated | PMID: 11238599 |
| LFTY1 | TDGF1 | CellPhoneDB, curated | PMID: 22710174 |
| LFTY2 | TDGF1 | CellPhoneDB, curated | PMID: 22710174 |
| LGR4 | RSPO3 | CellPhoneDB, guidetopharmacology.org | |
| LGR5 | RSPO4 | CellPhoneDB, guidetopharmacology.org | |
| LGR5 | RSPO3 | CellPhoneDB, guidetopharmacology.org | |
| LGR5 | RSPO1 | CellPhoneDB, guidetopharmacology.org | |
| LGR5 | RSPO2 | CellPhoneDB, guidetopharmacology.org | |
| LIRA4 | BST2 | CellPhoneDB, curated | PMID: 19564354 |
| LOX5 | AL5AP | CellPhoneDB, guidetopharmacology.org | |
| LRP1 | ERFE | CellPhoneDB, InnateDB-All | |
| LRP5 | FAM3B | CellPhoneDB, InnateDB-All | |
| LSHB | LSHR | CellPhoneDB, I2D | |
| LYAM1 | SELPL | CellPhoneDB, curated | PMC: 2431087, PMID: 8892633, P[illegible] |
| LYAM1 | CD34 | CellPhoneDB, curated | PMID: 7692600, PMID: 8977216 |
| LYAM2 | GSLG1 | CellPhoneDB, curated | PMID: 11404363 |
| LYAM2 | SELPL | CellPhoneDB, curated | PMC: 4571854 |
| LYAM3 | CD34 | CellPhoneDB, curated | PMID: 18606703 |
| LYAM3 | SELPL | CellPhoneDB, curated | PMID: 19118202, PMID: 9829984 |
| LYAM3 | CD24 | CellPhoneDB, curated | PMID: 9129046 |
| MCH | MCHR2 | CellPhoneDB, I2D | |
| MCH | MCHR1 | CellPhoneDB, I2D | |
| MCP | JAG1 | CellPhoneDB, curated | PMID: 23086448 |
| MERTK | GAS6 | CellPhoneDB, guidetopharmacology.org | |
| MET | HGF | CellPhoneDB, curated | uniprot |

| | | | |
|---|---|---|---|
| MIF | TNR14 | CellPhoneDB, I2D,IntAct | |
| MIF | TR10D | CellPhoneDB, I2D,IntAct | |
| MK | LRP1 | CellPhoneDB, I2D,InnateDB-All | |
| MK | ALK | CellPhoneDB, curated | uniprot |
| MK | PTPRZ | CellPhoneDB, curated | PMID: 10212223 |
| MK | SORL | CellPhoneDB, InnateDB-All | |
| MTLR | MOTI | CellPhoneDB, I2D | |
| MTLR | GHRL | CellPhoneDB, I2D,InnateDB-All | |
| NCAM1 | BDNF | CellPhoneDB, I2D | |
| NCAM1 | GDNF | CellPhoneDB, I2D | |
| NCTR3 | BAG6 | CellPhoneDB, curated | uniprot |
| NCTR3 | NR3L1 | CellPhoneDB, curated | uniprot |
| NEU1 | OXYR | CellPhoneDB, I2D | |
| NEU2 | V1AR | CellPhoneDB, I2D,IMEx,IntAct | |
| NEU2 | V1BR | CellPhoneDB, I2D | |
| NEU2 | OXYR | CellPhoneDB, I2D | |
| NEU2 | V2R | CellPhoneDB, I2D,InnateDB-All | |
| NGF | TNR16 | CellPhoneDB, curated | uniprot |
| NMDE1 | IL16 | CellPhoneDB, I2D | |
| NMDE2 | IL16 | CellPhoneDB, I2D | |
| NMDE4 | IL16 | CellPhoneDB, I2D | |
| NMS | NMUR1 | CellPhoneDB, guidetopharmacology.org | |
| NMS | NMUR2 | CellPhoneDB, guidetopharmacology.org | |
| NMU | NMUR1 | CellPhoneDB, guidetopharmacology.org | |
| NMU | NMUR2 | CellPhoneDB, guidetopharmacology.org | |
| NOTC1 | DLL4 | CellPhoneDB, curated | PMID: 22353464 |
| NOTC1 | DLL3 | CellPhoneDB, curated | PMID: 22353464 |
| NOTC1 | JAG2 | CellPhoneDB, curated | PMID: 22353464 |
| NOTC1 | DLK1 | CellPhoneDB, curated | PMID: 22353464 |
| NOTC1 | WNT4 | CellPhoneDB, I2D | |
| NOTC1 | JAG1 | CellPhoneDB, curated | PMID: 22353464 |
| NOTC1 | NOV | CellPhoneDB, I2D,InnateDB-All | |
| NOTC2 | DLL3 | CellPhoneDB, curated | PMID: 22353464 |
| NOTC2 | JAG2 | CellPhoneDB, curated | PMID: 22353464 |
| NOTC2 | DLL4 | CellPhoneDB, curated | PMID: 22353464 |
| NOTC2 | IL24 | CellPhoneDB, IMEx,InnateDB-All,MINT | |
| NOTC3 | JAG2 | CellPhoneDB, curated | PMID: 22353464 |
| NOTC4 | DLL4 | CellPhoneDB, curated | PMID: 22353464 |
| NOTC4 | JAG2 | CellPhoneDB, curated | PMID: 22353464 |
| NOTC4 | DLL3 | CellPhoneDB, curated | PMID: 22353464 |
| NPBW1 | NPW | CellPhoneDB, guidetopharmacology.org | |
| NPBW1 | NPB | CellPhoneDB, guidetopharmacology.org | |
| NPBW2 | NPB | CellPhoneDB, guidetopharmacology.org | |
| NPBW2 | NPW | CellPhoneDB, guidetopharmacology.org | |
| NPFF1 | NPVF | CellPhoneDB, guidetopharmacology.org | |
| NPS | NPSR1 | CellPhoneDB, guidetopharmacology.org | |
| NRG1 | LGR4 | CellPhoneDB, InnateDB-All | |
| NRG1 | LSR | CellPhoneDB, InnateDB-All | |
| NRG1 | M4A4A | CellPhoneDB, IMEx,IntAct | |
| NRG1 | NETO2 | CellPhoneDB, InnateDB-All | |
| NRG2 | ERBB3 | CellPhoneDB, curated | uniprot |
| NTF4 | NTRK2 | CellPhoneDB, I2D,IMEx,InnateDB-All,IntAct | |

| | | | |
|---|---|---|---|
| NTR1 | NEUT | CellPhoneDB, guidetopharmacology.org | |
| OREX | OX2R | CellPhoneDB, guidetopharmacology.org | |
| OREX | OX1R | CellPhoneDB, guidetopharmacology.org | |
| OSTP | CCR8 | CellPhoneDB, IMEx,MINT | |
| OSTP | PE2R4 | CellPhoneDB, IMEx,MINT | |
| OX26 | QRFPR | CellPhoneDB, guidetopharmacology.org | |
| OX2G | MO2R1 | CellPhoneDB, curated | PMID: 22020332 |
| PACA | SCTR | CellPhoneDB, I2D,InnateDB-All | |
| PACA | PACR | CellPhoneDB, I2D,IMEx | |
| PACA | VIPR2 | CellPhoneDB, I2D | |
| PACA | VIPR1 | CellPhoneDB, I2D | |
| PACA | DPP4 | CellPhoneDB, InnateDB-All,MINT | |
| PAHO | NPY4R | CellPhoneDB, guidetopharmacology.org | |
| PAHO | NPY5R | CellPhoneDB, guidetopharmacology.org | |
| PDCD1 | PD1L1 | CellPhoneDB, curated | PMID: 23954143 |
| PDCD1 | FAM3C | CellPhoneDB, InnateDB-All | |
| PDCD1 | PD1L2 | CellPhoneDB, curated | PMID: 23954143 |
| PDGFA | PGFRA | CellPhoneDB, curated | uniprot |
| PDGFB | LRP1 | CellPhoneDB, I2D,InnateDB-All | |
| PDGFB | PGFRB | CellPhoneDB, curated | uniprot |
| PDGFB | GPR98 | CellPhoneDB, InnateDB-All | |
| PDGFB | PGFRA | CellPhoneDB, curated | uniprot |
| PE2R3 | GHRL | CellPhoneDB, InnateDB-All | |
| PECA1 | CD177 | CellPhoneDB, curated | PMID: 17580308 |
| PECA1 | CD38 | CellPhoneDB, curated | PMID: 9551996, PMID: 26407101 |
| PENK | MRGX1 | CellPhoneDB, guidetopharmacology.org | |
| PGFRA | PDGFC | CellPhoneDB, curated | uniprot |
| PGFRB | PDGFD | CellPhoneDB, curated | uniprot |
| PGRC2 | CC4L | CellPhoneDB, IMEx,IntAct | |
| PI2R | GHRL | CellPhoneDB, InnateDB-All | |
| PKR1 | PROK2 | CellPhoneDB, guidetopharmacology.org | |
| PKR2 | PROK2 | CellPhoneDB, guidetopharmacology.org | |
| PLF4 | CXCR3 | CellPhoneDB, curated | PMID: 24218476 |
| PLF4 | ACKR1 | CellPhoneDB, I2D,InnateDB | |
| PLXB1 | SEM4D | CellPhoneDB, curated | PMID: 22325954 |
| PLXB2 | SEM4D | CellPhoneDB, curated | PMID: 22325954 |
| PLXB2 | SEM4C | CellPhoneDB, curated | uniprot |
| PLXB2 | PTN | CellPhoneDB, I2D,IMEx,InnateDB-All,IntAct | |
| PLXB2 | SEM4G | CellPhoneDB, curated | uniprot |
| PLXC1 | SEM7A | CellPhoneDB, curated | PMID: 22325954 |
| PODXL | LYAM1 | CellPhoneDB, curated | PMID: 22814396 |
| PRL | PRLR | CellPhoneDB, I2D,IMEx,InnateDB,IntAct | |
| PRLHR | PRRP | CellPhoneDB, guidetopharmacology.org | |
| PRLR | CSHL | CellPhoneDB, guidetopharmacology.org | |
| PROK1 | PKR2 | CellPhoneDB, guidetopharmacology.org | |
| PROK1 | PKR1 | CellPhoneDB, guidetopharmacology.org | |
| PROS | TYRO3 | CellPhoneDB, guidetopharmacology.org | |
| PROS | UFO | CellPhoneDB, guidetopharmacology.org | |
| PRPRC | CD22 | CellPhoneDB, curated | PMID: 12115612 |
| PTHR | PTH2R | CellPhoneDB, guidetopharmacology.org | |
| PTHR | PTH1R | CellPhoneDB, guidetopharmacology.org | |
| PTHR | PRLHR | CellPhoneDB, guidetopharmacology.org | |

| | | | |
|---|---|---|---|
| PTHY | PTH2R | CellPhoneDB, I2D,InnateDB-All | |
| PTHY | PTH1R | CellPhoneDB, I2D,IMEx,InnateDB-All,IntAct | |
| PTN | PTPRS | CellPhoneDB, I2D,IMEx,InnateDB-All,IntAct | |
| PTN | PTPRZ | CellPhoneDB, curated | PMID: 25644401 |
| PTN | ALK | CellPhoneDB, curated | uniprot |
| PVR | TIGIT | CellPhoneDB, curated | PMID: 19815499, PMID:24987108 |
| PVR | TNFL9 | CellPhoneDB, InnateDB-All | |
| PVR | CD226 | CellPhoneDB, curated | PMID: 15607800, PMID: 24440149 |
| PVR | TACT | CellPhoneDB, curated | uniprot |
| PVR | PVRL3 | CellPhoneDB, curated | PMID: 19011627, PMID: 12740392 |
| PVRL1 | CADM3 | CellPhoneDB, curated | uniprot |
| PVRL1 | PVRL4 | CellPhoneDB, curated | PMID: 22902367 |
| PVRL1 | PVRL3 | CellPhoneDB, curated | PMID: 12740392 |
| PVRL2 | PVRL3 | CellPhoneDB, curated | PMID: 19011627,PMID: 12740392 |
| PYY | NPY5R | CellPhoneDB, I2D | |
| PYY | NPY4R | CellPhoneDB, guidetopharmacology.org | |
| PYY | NPY1R | CellPhoneDB, guidetopharmacology.org | |
| PYY | DPP4 | CellPhoneDB, InnateDB-All,MINT | |
| PYY | NPY2R | CellPhoneDB, guidetopharmacology.org | |
| REL1 | RXFP2 | CellPhoneDB, guidetopharmacology.org | |
| REL1 | RXFP1 | CellPhoneDB, guidetopharmacology.org | |
| REL2 | RXFP2 | CellPhoneDB, guidetopharmacology.org | |
| REL2 | RXFP1 | CellPhoneDB, guidetopharmacology.org | |
| REL2 | RL3R1 | CellPhoneDB, guidetopharmacology.org | |
| REL3 | RL3R1 | CellPhoneDB, guidetopharmacology.org | |
| REL3 | RXFP1 | CellPhoneDB, guidetopharmacology.org | |
| RL3R1 | INSL5 | CellPhoneDB, guidetopharmacology.org | |
| RL3R2 | INSL5 | CellPhoneDB, guidetopharmacology.org | |
| RL3R2 | REL3 | CellPhoneDB, guidetopharmacology.org | |
| RSPO1 | LGR4 | CellPhoneDB, guidetopharmacology.org | |
| RSPO1 | LGR6 | CellPhoneDB, guidetopharmacology.org | |
| RSPO2 | LGR6 | CellPhoneDB, guidetopharmacology.org | |
| RSPO2 | LGR4 | CellPhoneDB, guidetopharmacology.org | |
| RSPO3 | LGR6 | CellPhoneDB, guidetopharmacology.org | |
| RSPO4 | LGR4 | CellPhoneDB, guidetopharmacology.org | |
| RSPO4 | LGR6 | CellPhoneDB, guidetopharmacology.org | |
| RXFP2 | REL3 | CellPhoneDB, guidetopharmacology.org | |
| RYK | WNT3A | CellPhoneDB, I2D | |
| SAA1 | FPR2 | CellPhoneDB, guidetopharmacology.org | |
| SDF1 | CXCR4 | CellPhoneDB, curated | PMID: 24218476 |
| SDF1 | CXCR3 | CellPhoneDB, guidetopharmacology.org | |
| SECR | SCTR | CellPhoneDB, guidetopharmacology.org | |
| SECR | VIPR1 | CellPhoneDB, I2D,InnateDB-All | |
| SEM3E | PLXD1 | CellPhoneDB, curated | PMID: 22325954 |
| SEM4A | PLXD1 | CellPhoneDB, curated | PMID: 22325954 |
| SEM5A | PLXB3 | CellPhoneDB, curated | uniprot |
| SG3A1 | MARCO | CellPhoneDB, I2D,InnateDB-All | |
| SG3A1 | NOTC3 | CellPhoneDB, InnateDB-All | |
| SHPS1 | CD47 | CellPhoneDB, curated | PMID: 23602662 |
| SLIB | VIPR1 | CellPhoneDB, guidetopharmacology.org | |
| SLIB | GHRHR | CellPhoneDB, guidetopharmacology.org | |
| SOM2 | GHR | CellPhoneDB, guidetopharmacology.org | |

| | | | |
|---|---|---|---|
| SOMA | PRLR | CellPhoneDB, I2D,InnateDB-All,IntAct | |
| SOMA | GHR | CellPhoneDB, guidetopharmacology.org | |
| TA2R | GHRL | CellPhoneDB, InnateDB-All | |
| TACT | PVRL1 | CellPhoneDB, curated | PMID: 17971293 |
| TFF1 | FCRL4 | CellPhoneDB, IMEx,InnateDB-All,IntAct | |
| TFR1 | TN13B | CellPhoneDB, IMEx,IntAct | |
| TGFB1 | TGFBR1 | curated | |
| TGFB1 | TGFBR2 | curated | |
| TGFB1 | TGBR3 | CellPhoneDB, curated | uniprot |
| TGFB2 | TGBR3 | CellPhoneDB, curated | uniprot |
| TGFB3 | TGBR3 | CellPhoneDB, curated | uniprot |
| THYG | ASGR1 | CellPhoneDB, I2D | |
| TIE2 | ANGP4 | CellPhoneDB, curated | uniprot |
| TIE2 | ANGP1 | CellPhoneDB, curated | uniprot |
| TIGIT | PVRL3 | CellPhoneDB, curated | PMID: 1313846 |
| TIGIT | PVRL2 | CellPhoneDB, curated | PMID: 1313846, PMID:24987108 |
| TNF10 | TR10D | CellPhoneDB, curated | uniprot |
| TNF10 | RIPK1 | CellPhoneDB, InnateDB-All | |
| TNF11 | TNR11 | CellPhoneDB, curated | uniprot |
| TNF12 | TNR12 | CellPhoneDB, curated | uniprot |
| TNF12 | TNR25 | CellPhoneDB, I2D | |
| TNF13 | TNR6 | CellPhoneDB, I2D,InnateDB-All | |
| TNF13 | TNR17 | CellPhoneDB, curated | uniprot |
| TNF13 | TNR14 | CellPhoneDB, I2D,InnateDB-All | |
| TNF13 | TNR1A | CellPhoneDB, I2D,InnateDB-All | |
| TNF14 | TNR14 | CellPhoneDB, curated | uniprot |
| TNF14 | TNR3 | CellPhoneDB, curated | uniprot |
| TNF14 | TNF6B | CellPhoneDB, curated | uniprot |
| TNF15 | TNF6B | CellPhoneDB, curated | uniprot |
| TNF15 | TNR25 | CellPhoneDB, guidetopharmacology.org | |
| TNF18 | TNR18 | CellPhoneDB, curated | uniprot |
| TNF6B | TNFL6 | CellPhoneDB, curated | uniprot |
| TNFA | NOTC1 | CellPhoneDB, I2D | |
| TNFA | CELR2 | CellPhoneDB, InnateDB-All | |
| TNFA | ICOS | CellPhoneDB, InnateDB-All | |
| TNFA | GI24 | CellPhoneDB, IMEx,IntAct | |
| TNFA | RIPK1 | CellPhoneDB, I2D,IMEx,InnateDB,InnateDB-All,IntAct,MINT | |
| TNFA | TNR1A | CellPhoneDB, curated | uniprot |
| TNFA | TNR6 | CellPhoneDB, InnateDB-All | |
| TNFA | VGFR3 | CellPhoneDB, InnateDB-All | |
| TNFA | SEM4C | CellPhoneDB, InnateDB-All | |
| TNFA | TNR1B | CellPhoneDB, curated | uniprot |
| TNFA | DAG1 | CellPhoneDB, InnateDB-All | |
| TNFA | PTPRS | CellPhoneDB, InnateDB-All | |
| TNFB | TNR14 | CellPhoneDB, curated | uniprot |
| TNFB | TNR3 | CellPhoneDB, I2D | |
| TNFB | TNR1B | CellPhoneDB, curated | uniprot |
| TNFB | TNR1A | CellPhoneDB, curated | uniprot |
| TNFB | RIPK1 | CellPhoneDB, IMEx,InnateDB-All,MINT | |
| TNFL4 | TNR4 | CellPhoneDB, curated | uniprot |
| TNFL9 | I13R2 | CellPhoneDB, InnateDB-All | |
| TNFL9 | TNR9 | CellPhoneDB, curated | uniprot |

| | | | |
|---|---|---|---|
| TNFL9 | AGRG5 | CellPhoneDB, InnateDB-All | |
| TNFRSF13B | TNFSF13B | curated | uniprot |
| TNFRSF13C | TNFSF13B | curated | uniprot |
| TNFRSF17 | TNFSF13B | curated | uniprot |
| TNFSF13 | TNFRSF13B | curated | |
| TNR16 | IL2 | CellPhoneDB, I2D | |
| TNR16 | BDNF | CellPhoneDB, curated | uniprot |
| TNR16 | NTF4 | CellPhoneDB, curated | uniprot |
| TNR17 | TN13B | CellPhoneDB, curated | uniprot |
| TNR1A | TNFL6 | CellPhoneDB, I2D,InnateDB-All | |
| TNR3 | TNFC | CellPhoneDB, curated | uniprot |
| TNR5 | INSL3 | CellPhoneDB, I2D | |
| TNR5 | CD40L | CellPhoneDB, curated | PMID: 23602662 |
| TNR5 | TN13B | CellPhoneDB, I2D,InnateDB | |
| TNR6 | TNFL6 | CellPhoneDB, curated | uniprot |
| TNR8 | TNFL8 | CellPhoneDB, curated | uniprot |
| TPO | TPOR | CellPhoneDB, curated | uniprot |
| TR10A | TNF10 | CellPhoneDB, curated | uniprot |
| TR10A | FGFR4 | CellPhoneDB, InnateDB-All | |
| TR10B | TNFL6 | CellPhoneDB, I2D,InnateDB-All | |
| TR10B | TNF10 | CellPhoneDB, curated | uniprot |
| TR10C | TNF10 | CellPhoneDB, curated | uniprot |
| TR11B | TNF11 | CellPhoneDB, curated | uniprot |
| TR11B | TNF10 | CellPhoneDB, curated | uniprot |
| TR13B | TN13B | CellPhoneDB, curated | uniprot |
| TR13B | CD70 | CellPhoneDB, InnateDB-All | |
| TR13B | TNF13 | CellPhoneDB, curated | uniprot |
| TR13C | TN13B | CellPhoneDB, guidetopharmacology.org | |
| TRH | TRFR | CellPhoneDB, I2D | |
| TSHB | TSHR | CellPhoneDB, I2D | |
| TTHY | DDR1 | CellPhoneDB, I2D,IMEx,InnateDB-All,IntAct | |
| TTHY | RAGE | CellPhoneDB, I2D,IMEx,InnateDB-All,MINT | |
| TTHY | TNR16 | CellPhoneDB, IMEx,InnateDB-All,MINT | |
| TYRO3 | GAS6 | CellPhoneDB, guidetopharmacology.org | |
| UCN1 | CRFR2 | CellPhoneDB, guidetopharmacology.org | |
| UFO | GAS6 | CellPhoneDB, guidetopharmacology.org | |
| UFO | I15RA | CellPhoneDB, I2D | |
| UTS2 | UR2R | CellPhoneDB, guidetopharmacology.org | |
| UTS2B | UR2R | CellPhoneDB, I2D,InnateDB-All | |
| VCC1 | GPR35 | CellPhoneDB, guidetopharmacology.org | |
| VEGFD | VGFR3 | CellPhoneDB, curated | uniprot |
| VEGFD | VGFR2 | CellPhoneDB, curated | uniprot |
| VGFR1 | PLGF | CellPhoneDB, curated | uniprot |
| VGFR2 | VEGFC | CellPhoneDB, curated | uniprot |
| VGFR3 | PDGFC | CellPhoneDB, guidetopharmacology.org | |
| VGFR3 | VEGFC | CellPhoneDB, curated | uniprot |
| VIP | VIPR2 | CellPhoneDB, I2D | |
| VIP | V1AR | CellPhoneDB, IMEx,IntAct | |
| VIP | VIPR1 | CellPhoneDB, guidetopharmacology.org | |
| VIP | DPP4 | CellPhoneDB, InnateDB-All,MINT | |
| VSTM1 | AGRG3 | CellPhoneDB, IMEx,IntAct | |
| WISP3 | SORL | CellPhoneDB, InnateDB-All | |

| | | | |
|---|---|---|---|
| WNT1 | ROR2 | CellPhoneDB, InnateDB-All | |
| WNT1 | FZD3 | CellPhoneDB, I2D | |
| WNT1 | FZD8 | CellPhoneDB, I2D,InnateDB-All | |
| WNT1 | RYK | CellPhoneDB, I2D | |
| WNT1 | FZD1 | CellPhoneDB, curated | PMID: 24032637 |
| WNT1 | CD36 | CellPhoneDB, InnateDB | |
| WNT2 | FZD2 | CellPhoneDB, curated | PMID: 24032637 |
| WNT2 | FZD4 | CellPhoneDB, curated | PMID: 24032637 |
| WNT2 | FZD1 | CellPhoneDB, I2D | |
| WNT2 | FZD3 | CellPhoneDB, curated | PMID: 24032637 |
| WNT2B | FZD4 | CellPhoneDB, curated | PMID: 24032637 |
| WNT3 | FZD1 | CellPhoneDB, I2D | |
| WNT3A | FZD2 | CellPhoneDB, InnateDB-All | |
| WNT3A | FZD8 | CellPhoneDB, IMEx,IntAct | |
| WNT3A | FZD1 | CellPhoneDB, I2D,InnateDB-All | |
| WNT3A | LRP1 | CellPhoneDB, I2D | |
| WNT4 | SMO | CellPhoneDB, I2D | |
| WNT4 | FZD1 | CellPhoneDB, I2D | |
| WNT4 | FZD8 | CellPhoneDB, I2D | |
| WNT5A | FZD3 | CellPhoneDB, curated | PMID: 24032637 |
| WNT5A | FZD2 | CellPhoneDB, curated | PMID: 24032637 |
| WNT5A | ROR1 | CellPhoneDB, IMEx | |
| WNT5A | FZD1 | CellPhoneDB, curated | PMID: 24032637 |
| WNT5A | ANTR1 | CellPhoneDB, InnateDB-All | |
| WNT5A | EPHA7 | CellPhoneDB, InnateDB-All | |
| WNT5A | FZD5 | CellPhoneDB, curated | PMID: 24032637 |
| WNT5A | ROR2 | CellPhoneDB, I2D | |
| WNT7B | FZD10 | CellPhoneDB, curated | PMID: 24032637 |
| WNT7B | FZD4 | CellPhoneDB, curated | PMID: 24032637 |
| WNT7B | FZD1 | CellPhoneDB, curated | PMID: 24032637 |
| XCL1 | GPR98 | CellPhoneDB, InnateDB-All | |
| XCR1 | XCL2 | CellPhoneDB, curated | uniprot |
| XCR1 | XCL1 | CellPhoneDB, curated | PMID: 24218476 |

;, PMC: 3168865

!

ed in PMID: 24223577

MID: 8896607

PMID: 12393589

3

9

2

2/
