## Supplementary Table 5 for "Dissecting intratumor heterogeneity of nodal B cell lymphomas on the transcriptional, genetic, and drug response level"

| Drug | C1 | C2 | C3 | C4 | C5 |
|---|---|---|---|---|---|
| 10058-F4 | 15000 | 3000 | 600 | 120 | 24 |
| Acalabrutinib | 5000 | 1000 | 200 | 40 | 8 |
| AZD7762 | 15000 | 3000 | 600 | 120 | 24 |
| Bendamustine | 20000 | 4000 | 800 | 160 | 32 |
| BI6727 | 15000 | 3000 | 600 | 120 | 24 |
| BRD73954 | 15000 | 3000 | 600 | 120 | 24 |
| C646 | 15000 | 3000 | 600 | 120 | 24 |
| Ceritinib | 15000 | 3000 | 600 | 120 | 24 |
| Chlorambucil | 20000 | 4000 | 800 | 160 | 32 |
| Cladribine | 15000 | 3000 | 600 | 120 | 24 |
| CPI-169 | 15000 | 3000 | 600 | 120 | 24 |
| Crizotinib | 15000 | 3000 | 600 | 120 | 24 |
| Cytarabine | 20000 | 4000 | 800 | 160 | 32 |
| Dasatinib | 1000 | 200 | 40 | 8 | 1.6 |
| Doxorubicin | 1000 | 200 | 40 | 8 | 1.6 |
| Duvelisib | 1000 | 200 | 40 | 8 | 1.6 |
| Entospletinib | 15000 | 3000 | 600 | 120 | 24 |
| EPZ-6438 | 15000 | 3000 | 600 | 120 | 24 |
| ERK5-IN-1 | 15000 | 3000 | 600 | 120 | 24 |
| Etoposide | 20000 | 4000 | 800 | 160 | 32 |
| Everolimus | 2000 | 400 | 80 | 16 | 3.2 |
| EVP4593 | 5000 | 1000 | 200 | 40 | 8 |
| Filgotinib | 15000 | 3000 | 600 | 120 | 24 |
| Fludarabine | 10000 | 2000 | 400 | 80 | 16 |
| Ganetespib | 2000 | 400 | 80 | 16 | 3.2 |
| I-BET-762 | 5000 | 1000 | 200 | 40 | 8 |
| Ibrutinib | 1000 | 200 | 40 | 8 | 1.6 |
| Idelalisib | 2000 | 400 | 80 | 16 | 3.2 |
| IRAK-1\|4 Inh. | 10000 | 2000 | 400 | 80 | 16 |
| Lenalidomide | 5000 | 1000 | 200 | 40 | 8 |
| MLN-120B | 15000 | 3000 | 600 | 120 | 24 |
| Navitoclax | 2000 | 400 | 80 | 16 | 3.2 |
| Nutlin-3a | 20000 | 4000 | 800 | 160 | 32 |
| Obatoclax | 15000 | 3000 | 600 | 120 | 24 |
| Olaparib | 15000 | 3000 | 600 | 120 | 24 |
| OTX015 | 10000 | 2000 | 400 | 80 | 16 |
| Palbociclib | 15000 | 3000 | 600 | 120 | 24 |
| Panobinostat | 400 | 80 | 1.6 | 0.32 | 0.064 |
| Pomalidomide | 2000 | 400 | 80 | 16 | 3.2 |
| Pralatrexate | 15000 | 3000 | 600 | 120 | 24 |
| Resveratrol | 15000 | 3000 | 600 | 120 | 24 |
| Ribociclib | 15000 | 3000 | 600 | 120 | 24 |
| Romidepsin | 200 | 40 | 8 | 0.16 | 0.032 |
| Ruxolitinib | 15000 | 3000 | 600 | 120 | 24 |
| SCH772984 | 5000 | 1000 | 200 | 40 | 8 |
| Selinexor | 5000 | 1000 | 200 | 40 | 8 |
| SGC0946 | 15000 | 3000 | 600 | 120 | 24 |
| Temsirolimus | 20000 | 4000 | 800 | 160 | 32 |
| Thalidomide | 20000 | 4000 | 800 | 160 | 32 |
| Tirabrutinib | 1000 | 200 | 40 | 8 | 1.6 |

| | | | | | |
|---|---|---|---|---|---|
| TMP269 | 15000 | 3000 | 600 | 120 | 24 |
| Tofacitinib | 15000 | 3000 | 600 | 120 | 24 |
| Trametinib | 500 | 100 | 20 | 4 | 0.8 |
| Vemurafenib | 15000 | 3000 | 600 | 120 | 24 |
| Venetoclax | 400 | 80 | 1.6 | 0.32 | 0.064 |
| Vincristine | 10000 | 2000 | 400 | 80 | 16 |
| Vismodegib | 15000 | 3000 | 600 | 120 | 24 |
| Vorinostat | 10000 | 2000 | 400 | 80 | 16 |
