## Supplementary Table 6 for "Dissecting intratumor heterogeneity of nodal B cell lymphomas on the transcriptional, genetic, and drug response level"

| CHROM | POSITION | CD32High COV | CD32High AC | CD32Low COV | CD32Low AC | HealthyB COV | HealthyB AC | Tumor COV | Tumor AC | GENE | Differential AC | Private CD32Low | Private CD32High |
|---|---|---|---|---|---|---|---|---|---|---|---|---|---|
| 1 | 232601083 | 174 | 0.67 | 109 | 0.02 | 146 | 0.00 | 225 | 0.04 | SIPA1L2 | 0.65 | FALSE | TRUE |
| 1 | 34158599 | 219 | 0.94 | 189 | 1.06 | 297 | 0.01 | 282 | 0.71 | CSMD2 | -0.12 | FALSE | FALSE |
| 1 | 55525194 | 135 | 0.00 | 130 | 0.88 | 185 | 0.00 | 155 | 0.59 | PCSK9 | -0.88 | TRUE | FALSE |
| 1 | 74549888 | 211 | 0.92 | 139 | 0.95 | 132 | 0.01 | 314 | 0.68 | LRRIQ3 | -0.03 | FALSE | FALSE |
| 1 | 82402497 | 47 | 0.77 | 33 | 0.67 | 25 | 0.24 | 57 | 1.00 | LPHN2 | 0.10 | FALSE | FALSE |
| 2 | 116535384 | 163 | 0.78 | 83 | 0.80 | 98 | 0.00 | 196 | 0.75 | DPP10 | -0.02 | FALSE | FALSE |
| 2 | 120383231 | 216 | 0.02 | 108 | 0.62 | 175 | 0.00 | 241 | 0.35 | PCDP1 | -0.60 | TRUE | FALSE |
| 2 | 121747745 | 257 | 0.91 | 287 | 0.92 | 360 | 0.01 | 394 | 0.78 | GLI2 | -0.01 | FALSE | FALSE |
| 2 | 166908259 | 32 | 0.81 | 8 | 1.00 | 34 | 0.00 | 58 | 0.42 | SCN1A | -0.19 | FALSE | FALSE |
| 2 | 178257571 | 53 | 0.94 | 65 | 0.86 | 79 | 0.08 | 61 | 0.80 | AGPS | 0.08 | FALSE | FALSE |
| 2 | 187531911 | 55 | 0.00 | 26 | 0.99 | 31 | 0.00 | 72 | 0.39 | ITGAV | -0.99 | TRUE | FALSE |
| 2 | 220159854 | 89 | 0.85 | 107 | 0.83 | 153 | 0.06 | 128 | 0.78 | PTPRN | 0.01 | FALSE | FALSE |
| 2 | 231112641 | 110 | 0.99 | 80 | 0.84 | 88 | 0.02 | 151 | 0.79 | SP140 | 0.15 | FALSE | FALSE |
| 2 | 89160093 | 193 | 0.87 | 117 | 0.12 | 84 | 0.00 | 260 | 0.19 | IGKJ5 | 0.75 | FALSE | TRUE |
| 2 | 96810581 | 117 | 1.01 | 183 | 0.97 | 189 | 0.01 | 181 | 0.83 | DUSP2 | 0.04 | FALSE | FALSE |
| 3 | 146312828 | 15 | 0.40 | 8 | 1.00 | 12 | 0.00 | 16 | 0.75 | PLSCR5 | -0.60 | FALSE | FALSE |
| 3 | 25805760 | 150 | 0.00 | 131 | 0.68 | 114 | 0.00 | 191 | 0.39 | NGLY1 | -0.68 | TRUE | FALSE |
| 3 | 30713277 | 170 | 1.78 | 141 | 1.78 | 240 | 0.07 | 233 | 1.63 | TGFBR2 | 0.00 | FALSE | FALSE |
| 3 | 67705007 | 66 | 0.71 | 52 | 0.00 | 89 | 0.00 | 90 | 0.00 | SUCLG2 | 0.71 | FALSE | TRUE |
| 4 | 126336270 | 212 | 1.05 | 135 | 0.92 | 144 | 0.04 | 264 | 0.85 | FAT4 | 0.13 | FALSE | FALSE |
| 4 | 156723467 | 90 | 0.76 | 77 | 0.96 | 84 | 0.02 | 155 | 0.84 | GUCY1B3 | -0.21 | FALSE | FALSE |
| 4 | 183696222 | 124 | 0.46 | 61 | 0.05 | 158 | 0.03 | 148 | 0.19 | TENM3 | 0.41 | FALSE | TRUE |
| 4 | 25864328 | 13 | 1.08 | 18 | 1.11 | 19 | 0.10 | 21 | 0.77 | SEL1L3 | -0.03 | FALSE | FALSE |
| 4 | 31144280 | 214 | 0.94 | 201 | 0.84 | 235 | 0.03 | 234 | 0.82 | PCDH7 | 0.10 | FALSE | FALSE |
| 4 | 38134776 | 162 | 0.10 | 110 | 0.04 | 124 | 0.00 | 239 | 0.03 | TBC1D1 | 0.06 | FALSE | FALSE |
| 4 | 57384859 | 105 | 1.07 | 92 | 0.00 | 77 | 0.00 | 138 | 0.18 | ARL9 | 1.07 | FALSE | TRUE |
| 5 | 1294772 | 141 | 1.02 | 193 | 1.04 | 270 | 0.04 | 264 | 0.80 | TERT | -0.03 | FALSE | FALSE |
| 5 | 175906252 | 280 | 0.68 | 175 | 0.03 | 180 | 0.00 | 356 | 0.11 | FAF2 | 0.65 | FALSE | TRUE |
| 5 | 176882923 | 185 | 1.01 | 227 | 0.89 | 251 | 0.05 | 275 | 0.62 | PRR7 | 0.12 | FALSE | FALSE |
| 5 | 178418890 | 286 | 0.88 | 290 | 0.83 | 352 | 0.03 | 360 | 0.68 | GRM6 | 0.05 | FALSE | FALSE |
| 5 | 179393977 | 118 | 1.00 | 110 | 1.14 | 133 | 0.06 | 138 | 0.83 | RNF130 | -0.14 | FALSE | FALSE |
| 6 | 129799844 | 138 | 0.71 | 42 | 0.05 | 114 | 0.00 | 189 | 0.19 | LAMA2 | 0.66 | FALSE | TRUE |
| 6 | 26032147 | 242 | 1.23 | 360 | 0.98 | 444 | 0.05 | 430 | 0.75 | HIST1H3B | 0.25 | FALSE | FALSE |
| 6 | 26104436 | 82 | 0.00 | 90 | 1.00 | 113 | 0.00 | 147 | 0.63 | HIST1H4C | -1.00 | TRUE | FALSE |
| 6 | 26234877 | 86 | 0.65 | 112 | 1.11 | 125 | 0.05 | 134 | 0.69 | HIST1H1D | -0.46 | FALSE | FALSE |
| 6 | 32159568 | 49 | 1.01 | 57 | 0.03 | 67 | 0.03 | 67 | 0.12 | GPSM3 | 0.97 | FALSE | TRUE |
| 6 | 93955155 | 56 | 0.10 | 24 | 0.00 | 41 | 0.00 | 101 | 0.07 | EPHA7 | 0.10 | FALSE | FALSE |
| 7 | 14622705 | 271 | 0.85 | 192 | 1.05 | 179 | 0.02 | 359 | 0.76 | DGKB | -0.20 | FALSE | FALSE |
| 7 | 150164098 | 135 | 1.05 | 127 | 0.86 | 124 | 0.00 | 165 | 0.94 | GIMAP8 | 0.19 | FALSE | FALSE |
| 7 | 154379756 | 317 | 1.02 | 308 | 0.92 | 384 | 0.02 | 360 | 0.55 | DPP6 | 0.10 | FALSE | FALSE |
| 7 | 39379551 | 353 | 1.00 | 307 | 0.93 | 428 | 0.03 | 459 | 0.67 | POU6F2 | 0.08 | FALSE | FALSE |
| 7 | 45104209 | 81 | 1.04 | 61 | 1.08 | 101 | 0.06 | 97 | 0.76 | CCM2 | -0.04 | FALSE | FALSE |
| 8 | 128748842 | 135 | 0.01 | 175 | 0.58 | 147 | 0.01 | 225 | 0.49 | MYC | -0.57 | TRUE | FALSE |
| 8 | 17206550 | 99 | 1.08 | 69 | 0.94 | 95 | 0.04 | 144 | 0.48 | MTMR7 | 0.15 | FALSE | FALSE |
| 8 | 24197030 | 164 | 0.80 | 112 | 0.05 | 107 | 0.00 | 203 | 0.10 | ADAM28 | 0.75 | FALSE | TRUE |
| 8 | 93074820 | 388 | 0.93 | 373 | 0.93 | 407 | 0.02 | 466 | 0.87 | RUNX1T1 | 0.00 | FALSE | FALSE |
| 9 | 101983318 | 149 | 1.02 | 101 | 0.12 | 105 | 0.00 | 181 | 0.09 | ALG2 | 0.90 | FALSE | TRUE |
| 9 | 109690781 | 492 | 1.01 | 536 | 0.99 | 571 | 0.04 | 564 | 0.78 | ZNF462 | 0.01 | FALSE | FALSE |
| 9 | 116823823 | 90 | 0.02 | 72 | 0.90 | 81 | 0.00 | 114 | 0.81 | AMBP | -0.88 | TRUE | FALSE |
| 9 | 138011411 | 325 | 1.03 | 359 | 0.95 | 456 | 0.02 | 397 | 0.86 | OLFM1 | 0.08 | FALSE | FALSE |
| 9 | 140351870 | 316 | 0.93 | 305 | 0.94 | 399 | 0.02 | 415 | 0.75 | NSMF | -0.01 | FALSE | FALSE |
| 9 | 79324693 | 152 | 0.96 | 109 | 0.76 | 135 | 0.04 | 199 | 0.70 | PRUNE2 | 0.20 | FALSE | FALSE |
| 9 | 8499743 | 101 | 1.00 | 84 | 0.91 | 111 | 0.02 | 164 | 0.94 | PTPRD | 0.09 | FALSE | FALSE |
| 10 | 37506676 | 97 | 0.00 | 25 | 0.69 | 34 | 0.00 | 67 | 0.78 | ANKRD30A | -0.69 | TRUE | FALSE |
| 10 | 56138604 | 119 | 0.77 | 56 | 0.95 | 58 | 0.14 | 116 | 0.62 | PCDH15 | -0.18 | FALSE | FALSE |
| 10 | 77160060 | 330 | 0.47 | 262 | 0.09 | 356 | 0.00 | 332 | 0.10 | ZNF503 | 0.38 | FALSE | TRUE |
| 11 | 123480970 | 190 | 0.80 | 158 | 0.05 | 184 | 0.01 | 206 | 0.08 | GRAMD1B | 0.74 | FALSE | TRUE |
| 11 | 124266695 | 173 | 0.18 | 155 | 0.00 | 145 | 0.00 | 246 | 0.03 | OR8B3 | 0.18 | FALSE | FALSE |
| 11 | 55406604 | 245 | 1.24 | 159 | 1.18 | 160 | 0.03 | 301 | 1.09 | OR4P4 | 0.06 | FALSE | FALSE |
| 11 | 65999698 | 68 | 0.94 | 64 | 0.59 | 89 | 0.05 | 106 | 0.64 | PACS1 | 0.35 | FALSE | FALSE |
| 11 | 66235677 | 137 | 0.93 | 162 | 0.85 | 195 | 0.01 | 193 | 0.85 | PELI3 | 0.08 | FALSE | FALSE |
| 11 | 67177064 | 170 | 0.87 | 165 | 0.77 | 240 | 0.03 | 260 | 0.74 | TBC1D10C | 0.09 | FALSE | FALSE |
| 11 | 76892456 | 46 | 0.00 | 47 | 0.80 | 57 | 0.00 | 45 | 0.98 | MYO7A | -0.80 | TRUE | FALSE |
| 11 | 7723323 | 50 | 1.00 | 41 | 1.08 | 40 | 0.10 | 64 | 0.45 | OVCH2 | -0.07 | FALSE | FALSE |
| 11 | 9595744 | 24 | 1.17 | 35 | 0.92 | 41 | 0.10 | 43 | 0.93 | WEE1 | 0.25 | FALSE | FALSE |
| 12 | 108634227 | 299 | 0.08 | 451 | 1.73 | 371 | 0.01 | 453 | 1.09 | WSCD2 | -1.66 | TRUE | FALSE |
| 12 | 112572563 | 174 | 0.00 | 162 | 1.45 | 127 | 0.00 | 281 | 1.13 | TRAFD1 | -1.45 | TRUE | FALSE |
| 12 | 116418529 | 203 | 1.28 | 217 | 1.86 | 177 | 0.02 | 314 | 1.53 | MED13L | -0.57 | FALSE | FALSE |
| 12 | 25261587 | 327 | 1.20 | 352 | 0.08 | 203 | 0.09 | 492 | 0.22 | CASC1 | 1.12 | FALSE | TRUE |
| 12 | 49420573 | 205 | 1.56 | 384 | 1.78 | 300 | 0.05 | 316 | 1.27 | KMT2D | -0.21 | FALSE | FALSE |
| 12 | 49431346 | 207 | 1.42 | 338 | 0.97 | 275 | 0.04 | 304 | 0.81 | KMT2D | 0.45 | FALSE | FALSE |
| 13 | 109540825 | 65 | 0.00 | 52 | 0.20 | 51 | 0.00 | 75 | 0.22 | MYO16 | -0.20 | FALSE | FALSE |
| 13 | 112722458 | 27 | 0.00 | 31 | 1.06 | 49 | 0.00 | 38 | 0.11 | SOX1 | -1.06 | TRUE | FALSE |
| 13 | 41240349 | 27 | 0.98 | 22 | 1.02 | 69 | 0.06 | 41 | 0.47 | FOXO1 | -0.04 | FALSE | FALSE |
| 13 | 51522157 | 16 | 0.00 | 7 | 0.00 | 23 | 0.17 | 37 | 0.00 | RNASEH2B | 0.00 | FALSE | FALSE |
| 13 | 72204754 | 109 | 1.01 | 68 | 1.00 | 101 | 0.06 | 143 | 1.00 | DACH1 | 0.01 | FALSE | FALSE |
| 14 | 106067778 | 235 | 0.85 | 225 | 0.82 | 390 | 0.03 | 378 | 0.59 | IGHE | 0.03 | FALSE | FALSE |
| 14 | 106209306 | 20 | 0.69 | 12 | 0.00 | 48 | 0.00 | 36 | 0.07 | IGHG1 | 0.69 | FALSE | TRUE |
| 14 | 106329456 | 34 | 1.22 | 32 | 1.18 | 87 | 0.00 | 118 | 0.27 | IGHJ6 | 0.05 | FALSE | FALSE |
| 14 | 106329461 | 32 | 1.36 | 34 | 1.34 | 86 | 0.00 | 121 | 0.30 | IGHJ6 | 0.02 | FALSE | FALSE |
| 14 | 106329465 | 32 | 1.42 | 34 | 1.34 | 91 | 0.00 | 120 | 0.33 | IGHJ6 | 0.08 | FALSE | FALSE |
| 14 | 106330050 | 15 | 0.40 | 17 | 0.12 | 86 | 0.00 | 90 | 0.00 | IGHJ5 | 0.28 | FALSE | FALSE |
| 14 | 106330067 | 12 | 1.81 | 15 | 0.40 | 66 | 0.00 | 71 | 0.04 | IGHJ5 | 1.42 | FALSE | FALSE |
| 14 | 107169956 | 104 | 0.00 | 363 | 0.75 | 398 | 0.03 | 428 | 0.37 | IGHV1-69 | -0.75 | TRUE | FALSE |

| | | | | | | | | | | | | | |
|---|---|---|---|---|---|---|---|---|---|---|---|---|---|
| 14 | 107178827 | 21 | 2.12 | 44 | 1.06 | 29 | 0.00 | 34 | 1.53 | IGHV2-70 | 1.06 | FALSE | FALSE |
| 14 | 107178836 | 25 | 2.13 | 56 | 0.04 | 38 | 0.00 | 48 | 0.08 | IGHV2-70 | 2.09 | FALSE | TRUE |
| 14 | 107179009 | 82 | 2.17 | 197 | 1.05 | 241 | 0.07 | 236 | 0.70 | IGHV2-70 | 1.12 | FALSE | FALSE |
| 14 | 107179023 | 86 | 2.17 | 184 | 0.05 | 233 | 0.01 | 217 | 0.20 | IGHV2-70 | 2.12 | FALSE | TRUE |
| 14 | 21560770 | 26 | 0.16 | 38 | 0.11 | 29 | 0.35 | 46 | 0.64 | ZNF219 | 0.05 | FALSE | FALSE |
| 14 | 26917441 | 312 | 0.97 | 249 | 0.98 | 271 | 0.04 | 415 | 0.88 | NOVA1 | 0.00 | FALSE | FALSE |
| 14 | 29237390 | 244 | 0.01 | 264 | 0.92 | 281 | 0.00 | 304 | 0.54 | FOXG1 | -0.91 | TRUE | FALSE |
| 14 | 55817426 | 255 | 0.99 | 187 | 0.98 | 204 | 0.03 | 364 | 0.88 | FBXO34 | 0.00 | FALSE | FALSE |
| 14 | 96180284 | 249 | 0.72 | 282 | 0.03 | 353 | 0.00 | 349 | 0.14 | TCL1A | 0.69 | FALSE | TRUE |
| 14 | 96707131 | 206 | 0.82 | 246 | 1.09 | 313 | 0.01 | 297 | 0.93 | BDKRB2 | -0.27 | FALSE | FALSE |
| 15 | 30010879 | 91 | 0.51 | 72 | 0.95 | 165 | 0.04 | 120 | 1.05 | TJP1 | -0.45 | FALSE | FALSE |
| 15 | 41043818 | 86 | 0.50 | 47 | 0.07 | 99 | 0.00 | 116 | 0.15 | RMDN3 | 0.43 | FALSE | TRUE |
| 15 | 93563500 | 245 | 0.51 | 121 | 0.06 | 224 | 0.04 | 288 | 0.21 | CHD2 | 0.45 | FALSE | TRUE |
| 16 | 11348880 | 136 | 0.01 | 166 | 0.86 | 253 | 0.01 | 174 | 0.32 | SOCS1 | -0.84 | TRUE | FALSE |
| 16 | 11349099 | 168 | 0.02 | 165 | 0.89 | 243 | 0.00 | 204 | 0.53 | SOCS1 | -0.86 | TRUE | FALSE |
| 16 | 11349287 | 97 | 0.18 | 117 | 0.02 | 139 | 0.00 | 134 | 0.00 | SOCS1 | 0.16 | FALSE | FALSE |
| 16 | 30735448 | 390 | 0.92 | 352 | 0.81 | 461 | 0.06 | 452 | 0.80 | SRCAP | 0.11 | FALSE | FALSE |
| 16 | 46952662 | 184 | 0.78 | 209 | 0.88 | 284 | 0.03 | 257 | 0.64 | GPT2 | -0.11 | FALSE | FALSE |
| 16 | 7703931 | 63 | 0.96 | 66 | 0.89 | 95 | 0.11 | 111 | 0.76 | RBFOX1 | 0.07 | FALSE | FALSE |
| 16 | 84778799 | 149 | 0.97 | 140 | 0.04 | 164 | 0.00 | 211 | 0.10 | USP10 | 0.93 | FALSE | TRUE |
| 16 | 88061206 | 74 | 1.19 | 101 | 1.09 | 108 | 0.04 | 101 | 0.84 | BANP | 0.10 | FALSE | FALSE |
| 17 | 34149650 | 70 | 0.00 | 58 | 0.88 | 43 | 0.00 | 88 | 0.68 | TAF15 | -0.88 | TRUE | FALSE |
| 17 | 37944556 | 89 | 0.95 | 51 | 0.81 | 61 | 0.03 | 89 | 0.90 | IKZF3 | 0.14 | FALSE | FALSE |
| 17 | 39535403 | 165 | 0.21 | 187 | 0.00 | 196 | 0.00 | 239 | 0.05 | KRT34 | 0.21 | FALSE | FALSE |
| 17 | 48594699 | 135 | 0.81 | 107 | 0.99 | 118 | 0.03 | 187 | 0.75 | MYCBPAP | -0.18 | FALSE | FALSE |
| 18 | 44773299 | 148 | 0.00 | 188 | 0.95 | 203 | 0.00 | 218 | 0.60 | SKOR2 | -0.95 | TRUE | FALSE |
| 18 | 60985412 | 560 | 0.58 | 474 | 0.04 | 470 | 0.00 | 558 | 0.16 | BCL2 | 0.54 | FALSE | TRUE |
| 18 | 60985513 | 529 | 0.64 | 443 | 0.88 | 470 | 0.01 | 557 | 0.81 | BCL2 | -0.24 | FALSE | FALSE |
| 18 | 60985549 | 406 | 0.64 | 350 | 0.10 | 336 | 0.02 | 409 | 0.27 | BCL2 | 0.54 | FALSE | TRUE |
| 18 | 60985754 | 241 | 0.47 | 220 | 0.86 | 240 | 0.01 | 240 | 0.70 | BCL2 | -0.39 | FALSE | FALSE |
| 18 | 60985760 | 246 | 0.46 | 223 | 0.85 | 263 | 0.02 | 255 | 0.70 | BCL2 | -0.39 | FALSE | FALSE |
| 18 | 60985761 | 254 | 0.00 | 222 | 0.78 | 264 | 0.00 | 257 | 0.48 | BCL2 | -0.78 | TRUE | FALSE |
| 18 | 60985834 | 571 | 0.40 | 483 | 0.03 | 554 | 0.02 | 594 | 0.15 | BCL2 | 0.36 | FALSE | FALSE |
| 18 | 60985846 | 587 | 0.52 | 484 | 0.84 | 576 | 0.02 | 610 | 0.62 | BCL2 | -0.32 | FALSE | FALSE |
| 18 | 6244582 | 148 | 1.07 | 82 | 0.81 | 117 | 0.04 | 174 | 0.74 | L3MBTL4 | 0.26 | FALSE | FALSE |
| 18 | 74587572 | 66 | 1.09 | 43 | 1.04 | 82 | 0.02 | 91 | 1.24 | ZNF236 | 0.05 | FALSE | FALSE |
| 19 | 17186222 | 269 | 1.09 | 281 | 0.90 | 385 | 0.04 | 370 | 0.86 | HAUS8 | 0.19 | FALSE | FALSE |
| 19 | 17450365 | 122 | 0.68 | 149 | 0.04 | 183 | 0.01 | 163 | 0.10 | GTPBP3 | 0.64 | FALSE | TRUE |
| 19 | 18392267 | 9 | 0.00 | 11 | 0.36 | 22 | 0.00 | 13 | 0.92 | JUND | -0.36 | FALSE | FALSE |
| 19 | 19260084 | 125 | 1.21 | 159 | 0.80 | 215 | 0.01 | 170 | 0.73 | B,MEF2BNB-M | 0.41 | FALSE | FALSE |
| 19 | 3198836 | 226 | 1.02 | 191 | 0.07 | 256 | 0.05 | 257 | 0.20 | NCLN | 0.95 | FALSE | TRUE |
| 19 | 40947569 | 87 | 1.00 | 82 | 0.71 | 98 | 0.00 | 117 | 0.65 | SERTAD3 | 0.29 | FALSE | FALSE |
| 19 | 4292750 | 107 | 0.00 | 96 | 0.86 | 124 | 0.00 | 95 | 0.55 | TMIGD2 | -0.86 | TRUE | FALSE |
| 19 | 535920 | 402 | 0.00 | 459 | 0.82 | 580 | 0.00 | 548 | 0.60 | CDC34 | -0.81 | TRUE | FALSE |
| 19 | 54401359 | 152 | 0.82 | 137 | 0.06 | 158 | 0.00 | 199 | 0.14 | PRKCG | 0.76 | FALSE | TRUE |
| 20 | 31421613 | 33 | 0.06 | 20 | 0.86 | 25 | 0.00 | 45 | 0.61 | MAPRE1 | -0.80 | TRUE | FALSE |
| 20 | 42815250 | 146 | 0.00 | 187 | 0.72 | 202 | 0.00 | 222 | 0.57 | JPH2 | -0.72 | TRUE | FALSE |
| 20 | 56137765 | 120 | 1.08 | 122 | 0.83 | 153 | 0.01 | 145 | 0.95 | PCK1 | 0.25 | FALSE | FALSE |
| 21 | 46309301 | 253 | 0.89 | 259 | 0.90 | 320 | 0.04 | 326 | 0.84 | ITGB2 | -0.01 | FALSE | FALSE |
| 22 | 23063496 | 120 | 0.92 | 112 | 1.08 | 99 | 0.04 | 155 | 0.82 | IGLV3-19 | -0.16 | FALSE | FALSE |
| 22 | 23223296 | 129 | 0.86 | 115 | 0.96 | 159 | 0.03 | 253 | 0.64 | IGLV3-1 | -0.10 | FALSE | FALSE |
| 22 | 23223297 | 129 | 1.12 | 116 | 1.08 | 161 | 0.06 | 259 | 0.72 | IGLV3-1 | 0.05 | FALSE | FALSE |
| 22 | 23223344 | 137 | 1.10 | 132 | 1.08 | 171 | 0.07 | 249 | 0.72 | IGLV3-1 | 0.02 | FALSE | FALSE |
| 22 | 23223348 | 140 | 0.91 | 135 | 0.98 | 177 | 0.01 | 241 | 0.64 | IGLV3-1 | -0.08 | FALSE | FALSE |
| 22 | 23223385 | 139 | 2.00 | 147 | 2.01 | 164 | 0.09 | 223 | 1.41 | IGLV3-1 | -0.01 | FALSE | FALSE |
| 22 | 23223389 | 136 | 0.86 | 150 | 0.97 | 169 | 0.04 | 227 | 0.63 | IGLV3-1 | -0.11 | FALSE | FALSE |
| 22 | 23223432 | 169 | 1.05 | 195 | 0.95 | 171 | 0.05 | 248 | 0.88 | IGLV3-1 | 0.10 | FALSE | FALSE |
| 22 | 23223439 | 185 | 0.91 | 210 | 1.06 | 174 | 0.11 | 250 | 0.76 | IGLV3-1 | -0.15 | FALSE | FALSE |
| 22 | 23223563 | 216 | 0.91 | 279 | 1.01 | 243 | 0.02 | 327 | 0.67 | IGLV3-1 | -0.10 | FALSE | FALSE |
| 22 | 23230315 | 102 | 0.71 | 105 | 0.04 | 111 | 0.00 | 130 | 0.18 | IGLL5 | 0.67 | FALSE | TRUE |
| 22 | 23230347 | 82 | 0.00 | 97 | 0.10 | 100 | 0.00 | 115 | 0.00 | IGLL5 | -0.10 | FALSE | FALSE |
| 22 | 23230409 | 62 | 0.00 | 65 | 0.22 | 57 | 0.00 | 91 | 0.35 | IGLL5 | -0.22 | FALSE | FALSE |
| 22 | 23230410 | 59 | 0.78 | 62 | 0.81 | 57 | 0.00 | 87 | 0.91 | IGLL5 | -0.03 | FALSE | FALSE |
| 22 | 45741448 | 116 | 0.03 | 67 | 0.78 | 113 | 0.00 | 143 | 0.72 | SMC1B | -0.74 | TRUE | FALSE |
| 22 | 50470419 | 230 | 0.92 | 309 | 1.03 | 373 | 0.04 | 345 | 0.74 | TTLL8 | -0.11 | FALSE | FALSE |
| X | 100385012 | 88 | 0.00 | 32 | 1.00 | 62 | 0.00 | 72 | 0.46 | CENPI | -1.00 | TRUE | FALSE |
| X | 101581426 | 773 | 0.38 | 584 | 0.46 | 634 | 0.00 | 809 | 0.34 | NXF2 | -0.08 | FALSE | FALSE |
| X | 103349933 | 115 | 0.83 | 80 | 0.85 | 102 | 0.00 | 127 | 0.62 | SLC25A53 | -0.02 | FALSE | FALSE |
| X | 114426201 | 459 | 0.08 | 233 | 0.00 | 471 | 0.00 | 378 | 0.02 | RBMXL3 | 0.07 | FALSE | FALSE |
| X | 123517938 | 190 | 0.39 | 70 | 0.03 | 141 | 0.00 | 158 | 0.23 | TENM1 | 0.36 | FALSE | FALSE |
| X | 31089536 | 230 | 0.78 | 171 | 0.94 | 233 | 0.03 | 242 | 0.76 | FTHL17 | -0.16 | FALSE | FALSE |
| X | 36402949 | 273 | 1.18 | 142 | 0.85 | 135 | 0.03 | 287 | 0.89 | CXorf30 | 0.33 | FALSE | FALSE |
| X | 86887316 | 144 | 0.51 | 87 | 0.02 | 90 | 0.00 | 179 | 0.25 | KLHL4 | 0.49 | FALSE | TRUE |
