## Supplementary Table 7 for "Dissecting intratumor heterogeneity of nodal B cell lymphomas on the transcriptional, genetic, and drug response level"

| CHROM | POSITION | CD48High COV | CD48High AC | CD48Low COV | CD48Low AC | Tumor COV | Tumor AC | GENE | Differential AC | Private CD48Low | Private CD48High |
|---|---|---|---|---|---|---|---|---|---|---|---|
| 1 | 16259123 | 68 | 2.04 | 63 | 2.04 | 56 | 2.00 | SPEN | 0.00 | FALSE | FALSE |
| 1 | 36205026 | 86 | 2.04 | 82 | 2.04 | 71 | 1.95 | CLSPN | 0.00 | FALSE | FALSE |
| 1 | 39322636 | 61 | 2.04 | 77 | 2.04 | 81 | 2.04 | RRAGC | 0.00 | FALSE | FALSE |
| 1 | 75199055 | 74 | 1.19 | 69 | 0.80 | 53 | 0.92 | TYW3 | 0.39 | FALSE | FALSE |
| 1 | 155020319 | 71 | 0.78 | 76 | 1.08 | 71 | 0.83 | DCST1 | -0.30 | FALSE | FALSE |
| 1 | 155239338 | 71 | 0.89 | 88 | 1.02 | 66 | 1.08 | CLK2 | -0.13 | FALSE | FALSE |
| 1 | 157772385 | 86 | 0.81 | 76 | 1.02 | 73 | 0.98 | FCRL1 | -0.21 | FALSE | FALSE |
| 1 | 161641412 | 77 | 1.01 | 81 | 0.86 | 60 | 1.19 | FCGR2B | 0.15 | FALSE | FALSE |
| 1 | 196448308 | 71 | 1.09 | 87 | 0.92 | 86 | 0.90 | KCNT2 | 0.18 | FALSE | FALSE |
| 1 | 203186123 | 73 | 0.98 | 90 | 1.07 | 75 | 1.14 | CHIT1 | -0.09 | FALSE | FALSE |
| 1 | 203274817 | 84 | 0.95 | 94 | 0.81 | 43 | 1.04 | BTG2 | 0.14 | FALSE | FALSE |
| 2 | 29451828 | 54 | 0.36 | 59 | 0.14 | 38 | 0.00 | ALK | 0.22 | FALSE | FALSE |
| 2 | 46770213 | 54 | 0.62 | 41 | 1.03 | 34 | 0.94 | RHOQ | -0.40 | FALSE | FALSE |
| 2 | 46770214 | 52 | 0.65 | 37 | 1.22 | 29 | 1.10 | RHOQ | -0.57 | FALSE | FALSE |
| 2 | 46770914 | 113 | 0.75 | 128 | 1.03 | 59 | 0.88 | RHOQ | -0.29 | FALSE | FALSE |
| 2 | 86948104 | 69 | 1.16 | 78 | 1.05 | 43 | 0.95 | CHMP3 | 0.11 | FALSE | FALSE |
| 2 | 98351783 | 81 | 1.06 | 87 | 1.03 | 58 | 1.02 | ZAP70 | 0.03 | FALSE | FALSE |
| 2 | 110053377 | 77 | 0.93 | 89 | 1.06 | 51 | 0.92 | SH3RF3 | -0.13 | FALSE | FALSE |
| 2 | 131130738 | 77 | 0.96 | 67 | 0.85 | 52 | 0.75 | PTPN18 | 0.10 | FALSE | FALSE |
| 2 | 131130741 | 81 | 0.93 | 65 | 0.88 | 55 | 0.71 | PTPN18 | 0.05 | FALSE | FALSE |
| 2 | 189916114 | 91 | 1.10 | 77 | 0.98 | 57 | 0.82 | COL5A2 | 0.12 | FALSE | FALSE |
| 2 | 192700711 | 73 | 0.87 | 63 | 0.97 | 41 | 0.75 | SDPR | -0.11 | FALSE | FALSE |
| 2 | 210569238 | 76 | 0.54 | 75 | 0.00 | 81 | 0.13 | MAP2 | 0.54 | FALSE | TRUE |
| 2 | 216248845 | 80 | 1.10 | 79 | 0.78 | 70 | 0.93 | FN1 | 0.32 | FALSE | FALSE |
| 3 | 32022389 | 94 | 2.05 | 95 | 2.05 | 45 | 1.91 | OSBPL10 | 0.00 | FALSE | FALSE |
| 3 | 32022399 | 94 | 0.81 | 92 | 0.96 | 49 | 1.13 | OSBPL10 | -0.15 | FALSE | FALSE |
| 3 | 32022410 | 87 | 0.78 | 93 | 1.03 | 49 | 1.13 | OSBPL10 | -0.26 | FALSE | FALSE |
| 3 | 32022501 | 87 | 1.15 | 68 | 0.78 | 56 | 0.98 | OSBPL10 | 0.37 | FALSE | FALSE |
| 3 | 32022574 | 80 | 0.74 | 85 | 1.08 | 51 | 0.92 | OSBPL10 | -0.34 | FALSE | FALSE |
| 3 | 32022598 | 70 | 1.23 | 95 | 0.99 | 50 | 0.86 | OSBPL10 | 0.24 | FALSE | FALSE |
| 3 | 49161164 | 81 | 1.14 | 85 | 1.11 | 63 | 1.17 | LAMB2 | 0.03 | FALSE | FALSE |
| 3 | 65365008 | 56 | 0.99 | 65 | 1.07 | 46 | 1.15 | MAGI1 | -0.08 | FALSE | FALSE |
| 3 | 67049550 | 63 | 1.04 | 74 | 0.94 | 74 | 0.91 | KBTBD8 | 0.10 | FALSE | FALSE |
| 3 | 78766462 | 55 | 1.00 | 50 | 1.15 | 55 | 1.11 | ROBO1 | -0.14 | FALSE | FALSE |
| 3 | 108836887 | 85 | 0.84 | 74 | 1.02 | 55 | 0.97 | MORC1 | -0.18 | FALSE | FALSE |
| 3 | 110852566 | 78 | 1.02 | 66 | 0.90 | 70 | 1.05 | PVRL3 | 0.12 | FALSE | FALSE |
| 3 | 146233867 | 77 | 1.28 | 73 | 0.98 | 83 | 1.01 | PLSCR1 | 0.29 | FALSE | FALSE |
| 3 | 185003424 | 81 | 0.91 | 78 | 1.00 | 83 | 0.91 | MAP3K13 | -0.09 | FALSE | FALSE |
| 3 | 186502405 | 73 | 0.98 | 89 | 1.03 | 70 | 1.05 | EIF4A2 | -0.05 | FALSE | FALSE |
| 3 | 186502407 | 73 | 0.98 | 92 | 1.04 | 74 | 1.10 | EIF4A2 | -0.06 | FALSE | FALSE |
| 3 | 186649184 | 71 | 1.35 | 93 | 0.92 | 47 | 1.13 | ST6GAL1 | 0.43 | FALSE | FALSE |
| 3 | 189590731 | 75 | 1.17 | 82 | 1.17 | 61 | 1.00 | TP63 | 0.00 | FALSE | FALSE |
| 4 | 5461897 | 60 | 0.75 | 53 | 0.47 | 47 | 0.48 | STK32B | 0.28 | FALSE | FALSE |
| 4 | 23830210 | 56 | 0.79 | 59 | 0.57 | 44 | 0.56 | PPARGC1A | 0.23 | FALSE | FALSE |
| 4 | 143326366 | 60 | 1.13 | 85 | 1.16 | 62 | 0.96 | INPP4B | -0.03 | FALSE | FALSE |
| 4 | 156632039 | 66 | 0.84 | 81 | 0.88 | 68 | 0.81 | GUCY1A3 | -0.05 | FALSE | FALSE |
| 5 | 35039555 | 77 | 1.12 | 86 | 0.95 | 63 | 1.07 | AGXT2 | 0.16 | FALSE | FALSE |
| 5 | 76129198 | 101 | 0.89 | 80 | 0.95 | 62 | 0.92 | F2RL1 | -0.05 | FALSE | FALSE |
| 5 | 124079788 | 67 | 1.13 | 60 | 1.30 | 50 | 1.02 | ZNF608 | -0.17 | FALSE | FALSE |
| 5 | 140201541 | 83 | 0.99 | 90 | 0.91 | 56 | 0.98 | PCDHA5 | 0.08 | FALSE | FALSE |
| 6 | 10695270 | 74 | 1.05 | 65 | 1.23 | 44 | 1.25 | PAK1IP1 | -0.18 | FALSE | FALSE |
| 6 | 26056356 | 88 | 0.95 | 76 | 0.78 | 47 | 1.00 | HIST1H1C | 0.17 | FALSE | FALSE |
| 6 | 26156827 | 77 | 0.98 | 84 | 1.05 | 59 | 0.97 | HIST1H1E | -0.06 | FALSE | FALSE |
| 6 | 26216528 | 73 | 0.95 | 80 | 1.05 | 75 | 1.17 | HIST1H2BG | -0.10 | FALSE | FALSE |
| 6 | 26234827 | 69 | 1.01 | 100 | 0.88 | 57 | 1.07 | HIST1H1D | 0.13 | FALSE | FALSE |
| 6 | 27792128 | 66 | 0.93 | 99 | 0.76 | 43 | 0.71 | HIST1H4J | 0.17 | FALSE | FALSE |
| 6 | 31324560 | 74 | 0.91 | 96 | 1.09 | 53 | 0.96 | HLA-B | -0.17 | FALSE | FALSE |
| 6 | 31324659 | 65 | 0.85 | 88 | 1.05 | 48 | 0.77 | HLA-B | -0.20 | FALSE | FALSE |
| 6 | 31324660 | 69 | 0.89 | 86 | 1.07 | 48 | 0.85 | HLA-B | -0.18 | FALSE | FALSE |
| 6 | 32024572 | 80 | 1.05 | 86 | 0.74 | 47 | 0.74 | TNXB | 0.31 | FALSE | FALSE |
| 6 | 34213241 | 81 | 1.06 | 81 | 1.14 | 58 | 0.99 | HMGA1 | -0.08 | FALSE | FALSE |
| 6 | 37138609 | 90 | 0.80 | 88 | 0.86 | 51 | 0.80 | PIM1 | -0.06 | FALSE | FALSE |
| 6 | 43155568 | 57 | 0.22 | 70 | 0.41 | 55 | 0.07 | CUL9 | -0.19 | FALSE | FALSE |
| 6 | 55119985 | 82 | 1.05 | 85 | 0.87 | 75 | 0.98 | HCRTR2 | 0.18 | FALSE | FALSE |
| 6 | 111288748 | 64 | 0.96 | 98 | 0.96 | 83 | 1.08 | GTF3C6 | 0.00 | FALSE | FALSE |
| 6 | 134492852 | 73 | 0.90 | 90 | 1.09 | 67 | 0.85 | SGK1 | -0.19 | FALSE | FALSE |
| 6 | 134495673 | 79 | 1.09 | 74 | 0.97 | 63 | 0.84 | SGK1 | 0.12 | FALSE | FALSE |
| 6 | 134495688 | 77 | 1.06 | 78 | 1.10 | 62 | 1.19 | SGK1 | -0.04 | FALSE | FALSE |
| 6 | 138413252 | 72 | 0.88 | 91 | 0.94 | 60 | 0.95 | PERP | -0.06 | FALSE | FALSE |
| 7 | 5569230 | 81 | 1.06 | 111 | 1.09 | 63 | 0.84 | ACTB | -0.03 | FALSE | FALSE |
| 7 | 12666276 | 80 | 1.02 | 88 | 1.00 | 68 | 1.08 | SCIN | 0.02 | FALSE | FALSE |
| 7 | 42949991 | 71 | 0.86 | 80 | 1.07 | 55 | 1.04 | C7orf25 | -0.21 | FALSE | FALSE |
| 7 | 73922478 | 71 | 0.98 | 74 | 0.97 | 59 | 0.97 | GTF2IRD1 | 0.01 | FALSE | FALSE |
| 7 | 97823439 | 74 | 1.19 | 76 | 0.92 | 51 | 0.80 | LMTK2 | 0.27 | FALSE | FALSE |
| 7 | 107621137 | 75 | 0.90 | 69 | 0.89 | 59 | 1.00 | LAMB1 | 0.01 | FALSE | FALSE |

| | | | | | | | | | | | |
|---|---|---|---|---|---|---|---|---|---|---|---|
| 8 | 8749370 | 70 | 0.91 | 101 | 0.93 | 49 | 0.96 | MFHAS1 | -0.03 | FALSE | FALSE |
| 8 | 20107856 | 72 | 0.88 | 84 | 1.07 | 50 | 0.94 | LZTS1 | -0.19 | FALSE | FALSE |
| 8 | 67525070 | 95 | 0.93 | 86 | 0.98 | 45 | 0.91 | MYBL1 | -0.05 | FALSE | FALSE |
| 8 | 119296569 | 79 | 1.11 | 90 | 1.02 | 65 | 1.01 | AC023590.1 | 0.09 | FALSE | FALSE |
| 8 | 124219609 | 78 | 1.02 | 80 | 0.89 | 36 | 0.96 | FAM83A | 0.13 | FALSE | FALSE |
| 8 | 128750574 | 112 | 1.39 | 101 | 1.24 | 57 | 1.19 | MYC | 0.15 | FALSE | FALSE |
| 8 | 128750608 | 97 | 1.41 | 97 | 1.27 | 60 | 1.17 | MYC | 0.15 | FALSE | FALSE |
| 8 | 128750613 | 102 | 1.44 | 94 | 1.26 | 59 | 1.11 | MYC | 0.18 | FALSE | FALSE |
| 8 | 128750625 | 102 | 1.38 | 97 | 1.22 | 67 | 1.22 | MYC | 0.16 | FALSE | FALSE |
| 8 | 128750628 | 101 | 1.36 | 100 | 1.21 | 65 | 1.19 | MYC | 0.15 | FALSE | FALSE |
| 8 | 128750637 | 97 | 1.37 | 92 | 1.20 | 64 | 1.28 | MYC | 0.17 | FALSE | FALSE |
| 8 | 128750677 | 97 | 1.39 | 77 | 1.01 | 71 | 1.39 | MYC | 0.38 | FALSE | FALSE |
| 8 | 128750953 | 99 | 0.12 | 87 | 0.16 | 67 | 0.28 | MYC | -0.04 | FALSE | FALSE |
| 8 | 128751056 | 98 | 1.23 | 85 | 0.89 | 63 | 1.15 | MYC | 0.34 | FALSE | FALSE |
| 8 | 128751226 | 111 | 1.25 | 97 | 0.89 | 64 | 1.39 | MYC | 0.37 | FALSE | FALSE |
| 8 | 128751251 | 108 | 1.08 | 85 | 0.84 | 65 | 1.26 | MYC | 0.24 | FALSE | FALSE |
| 8 | 128751258 | 106 | 1.12 | 82 | 0.85 | 63 | 1.19 | MYC | 0.27 | FALSE | FALSE |
| 8 | 128752808 | 91 | 1.35 | 79 | 1.01 | 76 | 1.48 | MYC | 0.34 | FALSE | FALSE |
| 8 | 128753003 | 90 | 1.41 | 82 | 1.17 | 61 | 1.50 | MYC | 0.24 | FALSE | FALSE |
| 8 | 142161826 | 80 | 1.07 | 71 | 0.92 | 57 | 1.00 | DENND3 | 0.15 | FALSE | FALSE |
| 9 | 103046887 | 73 | 1.09 | 82 | 1.07 | 61 | 0.84 | INVS | 0.02 | FALSE | FALSE |
| 9 | 105767317 | 76 | 0.86 | 75 | 0.90 | 64 | 0.73 | CYLC2 | -0.04 | FALSE | FALSE |
| 9 | 127262554 | 87 | 1.18 | 97 | 0.97 | 74 | 0.97 | NR5A1 | 0.21 | FALSE | FALSE |
| 9 | 127618788 | 72 | 1.11 | 80 | 1.00 | 54 | 1.17 | WDR38 | 0.11 | FALSE | FALSE |
| 9 | 131904902 | 75 | 0.95 | 77 | 1.12 | 74 | 0.77 | PPP2R4 | -0.16 | FALSE | FALSE |
| 10 | 28151431 | 91 | 0.99 | 77 | 0.88 | 73 | 1.09 | ARMC4 | 0.11 | FALSE | FALSE |
| 10 | 63662036 | 91 | 0.97 | 69 | 1.16 | 64 | 0.86 | ARID5B | -0.19 | FALSE | FALSE |
| 10 | 83648964 | 71 | 1.01 | 81 | 0.93 | 65 | 0.91 | NRG3 | 0.07 | FALSE | FALSE |
| 10 | 101089332 | 66 | 0.96 | 89 | 1.06 | 41 | 1.05 | CNNM1 | -0.10 | FALSE | FALSE |
| 10 | 101380082 | 21 | 0.68 | 22 | 0.37 | 27 | 0.08 | SLC25A28 | 0.31 | FALSE | FALSE |
| 10 | 103909691 | 52 | 0.83 | 112 | 0.91 | 70 | 1.05 | PPRC1 | -0.09 | FALSE | FALSE |
| 11 | 830087 | 91 | 1.15 | 104 | 1.16 | 59 | 1.18 | EFCAB4A | -0.01 | FALSE | FALSE |
| 11 | 4929538 | 80 | 1.07 | 92 | 0.82 | 60 | 0.88 | OR51A7 | 0.25 | FALSE | FALSE |
| 11 | 56143539 | 42 | 0.34 | 52 | 0.08 | 30 | 0.00 | OR8U1 | 0.26 | FALSE | FALSE |
| 11 | 56143544 | 43 | 0.33 | 55 | 0.07 | 30 | 0.00 | OR8U1 | 0.26 | FALSE | FALSE |
| 11 | 56143556 | 43 | 0.33 | 53 | 0.08 | 32 | 0.00 | OR8U1 | 0.26 | FALSE | FALSE |
| 11 | 72851161 | 95 | 0.97 | 78 | 0.92 | 98 | 0.96 | FCHSD2 | 0.05 | FALSE | FALSE |
| 11 | 103075628 | 85 | 1.03 | 77 | 1.01 | 71 | 1.15 | DYNC2H1 | 0.03 | FALSE | FALSE |
| 11 | 103128449 | 72 | 0.88 | 85 | 1.25 | 74 | 1.08 | DYNC2H1 | -0.37 | FALSE | FALSE |
| 12 | 968478 | 81 | 0.96 | 70 | 1.02 | 64 | 0.96 | WNK1 | -0.06 | FALSE | FALSE |
| 12 | 7061307 | 98 | 0.77 | 83 | 1.01 | 54 | 0.98 | PTPN6 | -0.24 | FALSE | FALSE |
| 12 | 7069113 | 64 | 1.15 | 90 | 0.93 | 55 | 1.08 | PTPN6 | 0.22 | FALSE | FALSE |
| 12 | 12870982 | 73 | 0.98 | 88 | 0.98 | 53 | 1.00 | CDKN1B | 0.00 | FALSE | FALSE |
| 12 | 25398281 | 74 | 1.13 | 81 | 0.96 | 59 | 0.97 | KRAS | 0.17 | FALSE | FALSE |
| 12 | 56031495 | 79 | 1.01 | 78 | 0.84 | 57 | 1.11 | OR10P1 | 0.17 | FALSE | FALSE |
| 12 | 71946869 | 81 | 1.09 | 80 | 0.84 | 49 | 1.17 | LGR5 | 0.24 | FALSE | FALSE |
| 12 | 75807441 | 69 | 1.04 | 68 | 1.08 | 59 | 0.87 | GLIPR1L2 | -0.05 | FALSE | FALSE |
| 12 | 85497795 | 58 | 1.30 | 74 | 1.00 | 84 | 0.90 | LRRIQ1 | 0.31 | FALSE | FALSE |
| 12 | 89745630 | 70 | 0.15 | 84 | 0.05 | 64 | 0.10 | DUSP6 | 0.10 | FALSE | FALSE |
| 12 | 92538206 | 84 | 1.14 | 91 | 0.74 | 65 | 0.75 | BTG1 | 0.40 | FALSE | FALSE |
| 12 | 92538211 | 82 | 1.22 | 92 | 0.69 | 58 | 0.81 | BTG1 | 0.53 | FALSE | FALSE |
| 12 | 92538217 | 80 | 1.20 | 87 | 0.73 | 55 | 0.82 | BTG1 | 0.47 | FALSE | FALSE |
| 12 | 92539167 | 69 | 1.13 | 95 | 0.99 | 49 | 1.04 | BTG1 | 0.14 | FALSE | FALSE |
| 12 | 130833953 | 103 | 0.91 | 67 | 1.10 | 66 | 1.15 | PIWIL1 | -0.19 | FALSE | FALSE |
| 12 | 130898776 | 60 | 0.17 | 80 | 0.10 | 59 | 0.26 | RIMBP2 | 0.07 | FALSE | FALSE |
| 13 | 36744693 | 47 | 0.74 | 56 | 0.84 | 66 | 0.99 | CCDC169 | -0.10 | FALSE | FALSE |
| 14 | 31420123 | 91 | 1.08 | 71 | 1.18 | 82 | 0.85 | STRN3 | -0.10 | FALSE | FALSE |
| 14 | 47342665 | 88 | 1.12 | 79 | 1.17 | 76 | 0.99 | MDGA2 | -0.05 | FALSE | FALSE |
| 14 | 64596849 | 66 | 0.87 | 79 | 1.09 | 62 | 0.92 | SYNE2 | -0.22 | FALSE | FALSE |
| 14 | 73444911 | 70 | 1.02 | 67 | 1.22 | 72 | 1.13 | ZFYVE1 | -0.20 | FALSE | FALSE |
| 14 | 74035961 | 85 | 1.03 | 84 | 1.05 | 57 | 0.97 | ACOT2 | -0.01 | FALSE | FALSE |
| 14 | 74995356 | 70 | 0.76 | 92 | 1.00 | 65 | 1.10 | LTBP2 | -0.24 | FALSE | FALSE |
| 14 | 89076027 | 75 | 0.98 | 71 | 0.78 | 72 | 1.05 | ZC3H14 | 0.20 | FALSE | FALSE |
| 14 | 106329437 | 36 | 0.91 | 50 | 0.91 | 17 | 0.86 | IGHJ6 | 0.00 | FALSE | FALSE |
| 14 | 106330045 | 158 | 0.22 | 128 | 0.22 | 80 | 0.19 | IGHJ5 | 0.00 | FALSE | FALSE |
| 14 | 106330069 | 148 | 0.25 | 117 | 0.26 | 70 | 0.26 | IGHJ5 | -0.02 | FALSE | FALSE |
| 14 | 106330070 | 113 | 0.33 | 76 | 0.41 | 56 | 0.33 | IGHJ5 | -0.08 | FALSE | FALSE |
| 14 | 106725359 | 88 | 0.36 | 99 | 0.39 | 77 | 0.35 | IGHV3-23 | -0.02 | FALSE | FALSE |
| 14 | 106725391 | 90 | 0.50 | 116 | 0.50 | 83 | 0.49 | IGHV3-23 | 0.00 | FALSE | FALSE |
| 14 | 106725399 | 90 | 0.52 | 115 | 0.52 | 82 | 0.52 | IGHV3-23 | 0.00 | FALSE | FALSE |
| 14 | 106725401 | 91 | 0.52 | 114 | 0.51 | 79 | 0.51 | IGHV3-23 | 0.01 | FALSE | FALSE |
| 14 | 106725404 | 90 | 0.55 | 115 | 0.52 | 79 | 0.54 | IGHV3-23 | 0.03 | FALSE | FALSE |
| 14 | 106725427 | 90 | 0.64 | 127 | 0.61 | 89 | 0.56 | IGHV3-23 | 0.04 | FALSE | FALSE |
| 14 | 106725638 | 119 | 0.61 | 139 | 0.77 | 123 | 0.79 | IGHV3-23 | -0.16 | FALSE | FALSE |
| 14 | 107170002 | 108 | 0.79 | 121 | 0.74 | 86 | 0.84 | IGHV1-69 | 0.05 | FALSE | FALSE |
| 14 | 107170121 | 135 | 0.62 | 155 | 0.61 | 114 | 0.63 | IGHV1-69 | 0.01 | FALSE | FALSE |
| 14 | 107170160 | 156 | 0.64 | 163 | 0.75 | 117 | 0.83 | IGHV1-69 | -0.10 | FALSE | FALSE |

| | | | | | | | | | | | |
|---|---|---|---|---|---|---|---|---|---|---|---|
| 14 | 107170169 | 154 | 0.69 | 163 | 0.76 | 106 | 0.85 | IGHV1-69 | -0.07 | FALSE | FALSE |
| 14 | 107178945 | 57 | 0.90 | 51 | 0.88 | 35 | 0.71 | IGHV2-70 | 0.02 | FALSE | FALSE |
| 14 | 107178960 | 57 | 0.90 | 50 | 0.85 | 33 | 0.82 | IGHV2-70 | 0.04 | FALSE | FALSE |
| 14 | 107179064 | 51 | 0.67 | 61 | 0.91 | 28 | 0.81 | IGHV2-70 | -0.24 | FALSE | FALSE |
| 14 | 107179070 | 54 | 0.67 | 63 | 0.92 | 30 | 0.90 | IGHV2-70 | -0.24 | FALSE | FALSE |
| 15 | 31619569 | 36 | 0.97 | 37 | 1.27 | 17 | 1.08 | KLF13 | -0.31 | FALSE | FALSE |
| 15 | 42145192 | 101 | 0.87 | 95 | 0.95 | 64 | 1.05 | SPTBN5 | -0.08 | FALSE | FALSE |
| 15 | 42984185 | 73 | 1.12 | 74 | 1.16 | 59 | 1.04 | STARD9 | -0.04 | FALSE | FALSE |
| 15 | 45003766 | 67 | 0.98 | 79 | 1.11 | 77 | 0.88 | B2M | -0.14 | FALSE | FALSE |
| 15 | 45467525 | 78 | 0.89 | 99 | 1.01 | 67 | 0.85 | SHF | -0.12 | FALSE | FALSE |
| 15 | 80259981 | 78 | 1.04 | 82 | 0.90 | 65 | 0.00 | BCL2A1 | 0.15 | FALSE | FALSE |
| 15 | 80260012 | 75 | 0.98 | 82 | 0.95 | 71 | 0.00 | BCL2A1 | 0.04 | FALSE | FALSE |
| 16 | 1272090 | 69 | 0.83 | 78 | 1.13 | 51 | 0.80 | TPSG1 | -0.30 | FALSE | FALSE |
| 16 | 11348933 | 71 | 1.18 | 81 | 1.01 | 36 | 0.74 | SOCS1 | 0.17 | FALSE | FALSE |
| 16 | 11348951 | 77 | 1.06 | 92 | 1.04 | 40 | 0.77 | SOCS1 | 0.02 | FALSE | FALSE |
| 16 | 11873064 | 79 | 1.04 | 95 | 0.82 | 78 | 1.18 | ZC3H7A | 0.22 | FALSE | FALSE |
| 16 | 18896980 | 73 | 0.84 | 84 | 1.19 | 87 | 0.89 | SMG1 | -0.35 | FALSE | FALSE |
| 16 | 21086769 | 56 | 1.06 | 70 | 0.96 | 73 | 0.95 | DNAH3 | 0.09 | FALSE | FALSE |
| 16 | 24183630 | 71 | 1.15 | 67 | 0.89 | 60 | 1.05 | PRKCB | 0.27 | FALSE | FALSE |
| 16 | 66562903 | 76 | 0.94 | 84 | 1.05 | 59 | 0.93 | TK2 | -0.11 | FALSE | FALSE |
| 16 | 70557337 | 66 | 1.08 | 99 | 0.91 | 55 | 0.89 | COG4 | 0.18 | FALSE | FALSE |
| 16 | 83998897 | 77 | 0.96 | 76 | 0.94 | 53 | 0.89 | OSGIN1 | 0.01 | FALSE | FALSE |
| 16 | 89347080 | 74 | 1.02 | 89 | 0.99 | 45 | 1.00 | ANKRD11 | 0.03 | FALSE | FALSE |
| 17 | 7369066 | 75 | 2.04 | 102 | 2.04 | 62 | 1.97 | ZBTB4 | 0.00 | FALSE | FALSE |
| 17 | 76823312 | 45 | 0.41 | 44 | 0.05 | 30 | 0.14 | USP36 | 0.36 | FALSE | TRUE |
| 17 | 79203033 | 45 | 0.27 | 42 | 0.19 | 25 | 0.16 | ENTHD2 | 0.08 | FALSE | FALSE |
| 17 | 79479113 | 73 | 2.04 | 79 | 2.04 | 66 | 1.91 | ACTG1 | 0.00 | FALSE | FALSE |
| 17 | 79632245 | 74 | 2.04 | 88 | 2.04 | 47 | 1.99 | OXLD1 | 0.00 | FALSE | FALSE |
| 18 | 30350482 | 70 | 1.05 | 67 | 1.16 | 54 | 1.10 | KLHL14 | -0.11 | FALSE | FALSE |
| 18 | 31320250 | 62 | 1.06 | 86 | 1.07 | 62 | 1.09 | ASXL3 | -0.01 | FALSE | FALSE |
| 18 | 50942502 | 68 | 0.78 | 78 | 1.15 | 64 | 1.12 | DCC | -0.37 | FALSE | FALSE |
| 18 | 50942503 | 68 | 0.81 | 78 | 1.15 | 70 | 1.14 | DCC | -0.34 | FALSE | FALSE |
| 19 | 501767 | 104 | 0.33 | 115 | 0.12 | 75 | 0.16 | MADCAM1 | 0.21 | FALSE | FALSE |
| 19 | 6590112 | 80 | 1.36 | 68 | 1.11 | 40 | 0.92 | CD70 | 0.24 | FALSE | FALSE |
| 19 | 6665041 | 67 | 1.01 | 94 | 1.00 | 62 | 0.99 | TNFSF14 | 0.01 | FALSE | FALSE |
| 19 | 10655477 | 79 | 0.88 | 97 | 0.99 | 56 | 1.17 | ATG4D | -0.11 | FALSE | FALSE |
| 19 | 12902685 | 79 | 0.91 | 91 | 1.01 | 62 | 0.86 | JUNB | -0.11 | FALSE | FALSE |
| 19 | 12903583 | 71 | 0.84 | 96 | 0.96 | 60 | 1.02 | JUNB | -0.12 | FALSE | FALSE |
| 19 | 16436045 | 78 | 1.05 | 83 | 0.99 | 44 | 1.21 | KLF2 | 0.06 | FALSE | FALSE |
| 19 | 16436654 | 42 | 0.83 | 55 | 1.04 | 35 | 0.58 | KLF2 | -0.21 | FALSE | FALSE |
| 19 | 16436756 | 52 | 1.02 | 61 | 1.21 | 39 | 1.20 | KLF2 | -0.18 | FALSE | FALSE |
| 19 | 21299916 | 75 | 0.82 | 69 | 1.04 | 60 | 1.16 | ZNF714 | -0.22 | FALSE | FALSE |
| 19 | 38160872 | 82 | 1.30 | 74 | 0.94 | 73 | 1.15 | ZNF781 | 0.36 | FALSE | FALSE |
| 19 | 39412083 | 85 | 1.13 | 65 | 0.91 | 59 | 0.76 | SARS2 | 0.22 | FALSE | FALSE |
| 19 | 51171436 | 24 | 1.11 | 36 | 1.14 | 16 | 1.40 | SHANK1 | -0.03 | FALSE | FALSE |
| 19 | 54629913 | 68 | 0.96 | 86 | 0.90 | 61 | 0.87 | PRPF31 | 0.06 | FALSE | FALSE |
| 20 | 34459013 | 68 | 1.17 | 78 | 0.97 | 70 | 0.99 | PHF20 | 0.20 | FALSE | FALSE |
| 20 | 62194009 | 73 | 1.12 | 95 | 0.97 | 49 | 1.00 | HELZ2 | 0.15 | FALSE | FALSE |
| 21 | 45741670 | 96 | 0.89 | 96 | 0.87 | 60 | 1.02 | PFKL | 0.02 | FALSE | FALSE |
| 22 | 20103817 | 64 | 1.15 | 92 | 1.02 | 57 | 1.00 | RANBP1 | 0.13 | FALSE | FALSE |
| 22 | 22049289 | 80 | 0.92 | 87 | 0.96 | 63 | 1.13 | PPIL2 | -0.04 | FALSE | FALSE |
| 22 | 23223372 | 74 | 0.97 | 69 | 0.77 | 65 | 0.97 | IGLV3-1 | 0.20 | FALSE | FALSE |
| 22 | 23223389 | 75 | 1.09 | 66 | 0.74 | 60 | 0.95 | IGLV3-1 | 0.35 | FALSE | FALSE |
| 22 | 23223477 | 76 | 1.05 | 70 | 0.79 | 65 | 0.91 | IGLV3-1 | 0.26 | FALSE | FALSE |
| 22 | 23223520 | 72 | 1.02 | 68 | 0.69 | 63 | 0.88 | IGLV3-1 | 0.33 | FALSE | FALSE |
| 22 | 23230234 | 73 | 0.95 | 84 | 0.93 | 59 | 0.97 | IGLL5 | 0.03 | FALSE | FALSE |
| 22 | 37888774 | 59 | 1.04 | 98 | 1.00 | 60 | 0.88 | CARD10 | 0.04 | FALSE | FALSE |
| 22 | 42608072 | 59 | 0.97 | 83 | 1.16 | 55 | 0.97 | TCF20 | -0.19 | FALSE | FALSE |
| 22 | 50181290 | 89 | 0.97 | 99 | 0.89 | 72 | 0.77 | BRD1 | 0.08 | FALSE | FALSE |
| X | 12994426 | 100 | 0.84 | 70 | 1.08 | 74 | 0.99 | TMSB4X | -0.24 | FALSE | FALSE |
| X | 15554483 | 91 | 0.40 | 87 | 0.00 | 67 | 0.28 | BMX | 0.40 | FALSE | TRUE |
| X | 67937847 | 55 | 0.67 | 37 | 1.03 | 31 | 1.00 | STARD8 | -0.35 | FALSE | FALSE |
| X | 69479278 | 69 | 0.65 | 42 | 1.03 | 54 | 0.96 | P2RY4 | -0.37 | FALSE | FALSE |
| X | 70146015 | 65 | 0.36 | 37 | 0.00 | 40 | 0.13 | SLC7A3 | 0.36 | FALSE | FALSE |
| X | 83128586 | 68 | 0.42 | 40 | 0.00 | 46 | 0.23 | CYLC1 | 0.42 | FALSE | TRUE |
| X | 85997669 | 74 | 0.55 | 40 | 1.03 | 51 | 0.97 | DACH2 | -0.47 | FALSE | FALSE |
| X | 88008999 | 66 | 0.59 | 35 | 1.03 | 30 | 0.91 | CPXCR1 | -0.44 | FALSE | FALSE |
| X | 113965729 | 55 | 0.49 | 33 | 0.00 | 43 | 0.24 | HTR2C | 0.49 | FALSE | TRUE |
| X | 153627724 | 51 | 0.58 | 43 | 1.03 | 35 | 1.15 | RPL10 | -0.44 | FALSE | FALSE |
| 1 | 152323939 | 59 | 0.03 | 65 | 0.25 | 58 | 0.07 | FLG2 | -0.22 | FALSE | FALSE |
| 2 | 39164479 | 52 | 0.11 | 60 | 0.33 | 43 | 0.00 | ARHGEF33 | -0.22 | FALSE | FALSE |
| 4 | 121957535 | 74 | 0.08 | 105 | 0.58 | 71 | 0.43 | NDNF | -0.50 | TRUE | FALSE |
| 6 | 43155544 | 64 | 0.19 | 77 | 0.27 | 49 | 0.08 | CUL9 | -0.07 | FALSE | FALSE |
| 6 | 43155557 | 62 | 0.13 | 70 | 0.29 | 45 | 0.14 | CUL9 | -0.16 | FALSE | FALSE |
| 6 | 43155559 | 62 | 0.16 | 67 | 0.21 | 49 | 0.00 | CUL9 | -0.05 | FALSE | FALSE |
| 17 | 46608203 | 93 | 0.13 | 85 | 0.22 | 57 | 0.21 | HOXB1 | -0.08 | FALSE | FALSE |
| 18 | 57567669 | 80 | 0.00 | 93 | 0.11 | 67 | 0.03 | PMAIP1 | -0.11 | FALSE | FALSE |

| | | | | | | | | | | | |
|---|---|---|---|---|---|---|---|---|---|---|---|
| 19 | 51227736 | 74 | 0.08 | 101 | 0.16 | 57 | 0.18 | CLEC11A | -0.08 | FALSE | FALSE |
| 22 | 23235954 | 88 | 0.00 | 86 | 0.14 | 71 | 0.06 | IGLJ1 | -0.14 | FALSE | FALSE |
| 3 | 32022389 | 94 | 2.05 | 95 | 2.05 | 45 | 1.91 | OSBPL10 | 0.00 | FALSE | FALSE |
| 3 | 195512693 | 46 | 0.09 | 38 | 0.22 | 29 | 0.56 | MUC4 | -0.13 | FALSE | FALSE |
| 5 | 667958 | 54 | 0.08 | 79 | 0.18 | 48 | 0.34 | AC026740.1 | -0.11 | FALSE | FALSE |
| 6 | 116579808 | 74 | 0.00 | 63 | 0.03 | 68 | 0.33 | DSE | -0.03 | FALSE | FALSE |
| 6 | 116579835 | 68 | 0.03 | 69 | 0.00 | 70 | 0.35 | DSE | 0.03 | FALSE | FALSE |
| 14 | 106725244 | 91 | 0.47 | 85 | 0.35 | 85 | 0.50 | IGHV3-23 | 0.12 | FALSE | FALSE |
| 14 | 106725260 | 95 | 0.43 | 91 | 0.33 | 93 | 0.46 | IGHV3-23 | 0.10 | FALSE | FALSE |
| 14 | 106725266 | 95 | 0.54 | 89 | 0.38 | 92 | 0.51 | IGHV3-23 | 0.16 | FALSE | FALSE |
| 22 | 20708903 | 42 | 0.10 | 52 | 0.24 | 31 | 0.40 | FAM230A | -0.14 | FALSE | FALSE |
