## Supplementary Table 8 for "Dissecting intratumor heterogeneity of nodal B cell lymphomas on the transcriptional, genetic, and drug response level"

| | DLBCL1 | DLBCL2 | DLBCL3 | tFL1 | tFL2 | FL1 | FL2 | FL3 | FL4 | rLN1 | rLN2 | rLN3 |
|---|---|---|---|---|---|---|---|---|---|---|---|---|
| **Sequencing** | | | | | | | | | | | | |
| Number of Reads | 601,122,706 | 648,260,705 | 510,119,438 | 636,560,359 | 660,387,852 | 629,839,824 | 545,314,100 | 571,096,096 | 650,044,750 | 532,632,646 | 613,791,866 | 647,001,291 |
| Valid Barcodes | 97.20% | 97.60% | 96.70% | 97.30% | 97.20% | 97.10% | 97.10% | 97.10% | 96.90% | 97.10% | 97.40% | 97.20% |
| Sequencing Saturation | 77.50% | 89.80% | 85.80% | 94.50% | 95.30% | 94.20% | 93.80% | 91.50% | 95.60% | 88.50% | 94.40% | 95.70% |
| Q30 Bases in Barcode | 93.40% | 94.60% | 92.80% | 93.90% | 93.80% | 93.50% | 93.50% | 93.60% | 92.10% | 93.80% | 94.10% | 93.90% |
| Q30 Bases in RNA Read | 70.80% | 84.70% | 55.90% | 75.10% | 83.60% | 69.70% | 60.60% | 61.20% | 71.00% | 57.20% | 71.40% | 72.60% |
| Q30 Bases in UMI | 92.80% | 92.70% | 92.60% | 93.00% | 92.50% | 93.00% | 93.10% | 93.20% | 91.20% | 93.60% | 93.50% | 93.40% |
| **Mapping** | | | | | | | | | | | | |
| Reads Mapped to Genome | 81.00% | 89.40% | 77.50% | 80.40% | 89.90% | 81.10% | 77.50% | 79.80% | 84.90% | 75.60% | 84.80% | 86.20% |
| Reads Mapped Confidently to Genome | 77.20% | 84.60% | 73.80% | 76.60% | 86.30% | 76.60% | 72.50% | 74.90% | 80.60% | 71.60% | 80.30% | 82.10% |
| Reads Mapped Confidently to Intergenic Regions | 2.40% | 3.70% | 3.30% | 3.30% | 2.60% | 3.90% | 4.00% | 4.30% | 3.20% | 3.10% | 2.80% | 3.00% |
| Reads Mapped Confidently to Intronic Regions | 15.40% | 31.60% | 12.60% | 23.80% | 20.00% | 30.10% | 25.70% | 26.50% | 22.20% | 21.70% | 15.00% | 18.80% |
| Reads Mapped Confidently to Exonic Regions | 59.40% | 49.20% | 57.90% | 49.50% | 63.70% | 42.60% | 42.80% | 44.10% | 55.20% | 46.80% | 62.50% | 60.30% |
| Reads Mapped Confidently to Transcriptome | 56.20% | 46.40% | 55.20% | 46.70% | 60.80% | 39.90% | 40.20% | 41.60% | 52.60% | 44.20% | 59.80% | 57.60% |
| Reads Mapped Antisense to Gene | 1.10% | 1.30% | 0.60% | 1.20% | 0.90% | 1.30% | 1.30% | 1.20% | 0.90% | 1.10% | 0.70% | 0.90% |
| **Cells** | | | | | | | | | | | | |
| Estimated Number of Cells | 4,212 | 5,249 | 3,109 | 2,071 | 2,360 | 3,690 | 5,107 | 4,705 | 3,220 | 3,095 | 2,119 | 2,849 |
| Fraction Reads in Cells | 92.00% | 91.70% | 96.50% | 89.80% | 95.30% | 91.90% | 94.00% | 92.20% | 96.20% | 69.00% | 92.70% | 96.90% |
| Mean Reads per Cell | 142,716 | 123,501 | 164,078 | 307,368 | 279,825 | 170,688 | 106,777 | 121,380 | 201,877 | 172,094 | 289,661 | 227,097 |
| Median Genes per Cell | 3,275 | 1,488 | 2,450 | 1,451 | 1,583 | 1,103 | 834 | 1,300 | 1,126 | 1,486 | 1,227 | 1,100 |
| Total Genes Detected | 20,919 | 20,557 | 20,230 | 20,936 | 19,040 | 20,492 | 20,524 | 21,420 | 19,485 | 20,966 | 20,020 | 19,994 |
| Median UMI Counts per Cell | 14,791 | 3,922 | 8,267 | 3,562 | 4,975 | 2,282 | 1,768 | 2,971 | 2,838 | 3,759 | 4,021 | 3,113 |
